# Towards an integration of deep learning and neuroscience

**DOI:** 10.1101/058545

**Authors:** Adam H. Marblestone, Greg Wayne, Konrad P. Kording

## Abstract

Neuroscience has focused on the detailed implementation of computation, studying neural codes, dynamics and circuits. In machine learning, however, artificial neural networks tend to eschew precisely designed codes, dynamics or circuits in favor of brute force optimization of a cost function, often using simple and relatively uniform initial architectures. Two recent developments have emerged within machine learning that create an opportunity to connect these seemingly divergent perspectives. First, structured architectures are used, including dedicated systems for attention, recursion and various forms of short- and long-term memory storage. Second, cost functions and training procedures have become more complex and are varied across layers and over time. Here we think about the brain in terms of these ideas. We hypothesize that (1) the brain optimizes cost functions, (2) the cost functions are diverse and differ across brain locations and over development, and (3) optimization operates within a pre-structured architecture matched to the computational problems posed by behavior. In support of these hypotheses, we argue that a range of implementations of credit assignment through multiple layers of neurons are compatible with our current knowledge of neural circuitry, and that the brain’s specialized systems can be interpreted as enabling efficient optimization for specific problem classes. Such a heterogeneously optimized system, enabled by a series of interacting cost functions, serves to make learning data-efficient and precisely targeted to the needs of the organism. We suggest directions by which neuroscience could seek to refine and test these hypotheses.

## 1. Introduction

Machine learning and neuroscience speak different languages today. Brain science has discovered a dazzling array of brain areas (Solari and Stoner, 2015), cell types, molecules, cellular states, and mechanisms for computation and information storage. Machine learning, in contrast, has largely focused on instantiations of a single principle: function optimization. It has found that simple optimization objectives, like minimizing classification error, can lead to the formation of rich internal representations and powerful algorithmic capabilities in multilayer and recurrent networks (LeCun et al., 2015; Schmidhuber, 2015). Here we seek to connect these perspectives.

The artificial neural networks now prominent in machine learning were, of course, originally inspired by neuroscience (McCulloch and Pitts, 1943). While neuroscience has continued to play a role (Cox and Dean, 2014), many of the major developments were guided by insights into the mathematics of efficient optimization, rather than neuroscientific findings (Sutskever and Martens, 2013). The field has advanced from simple linear systems (Minsky and Papert, 1972), to nonlinear networks (Haykin, 1994), to deep and recurrent networks (Schmidhuber, 2015; LeCun et al., 2015). Backpropagation of error (Werbos, 1974, 1982; Rumelhart et al., 1986) enabled neural networks to be trained efficiently, by providing an efficient means to compute the gradient with respect to the weights of a multi-layer network. Methods of training have improved to include momentum terms, better weight initializations, conjugate gradients and so forth, evolving to the current breed of networks optimized using batch-wise stochastic gradient descent. These developments have little obvious connection to neuroscience.

We will argue here, however, that neuroscience and machine learning are again ripe for convergence. Three aspects of machine learning are particularly important in the context of this paper. First, machine learning has focused on the optimization of cost functions (**Figure 1A**).

**Fig 1:**
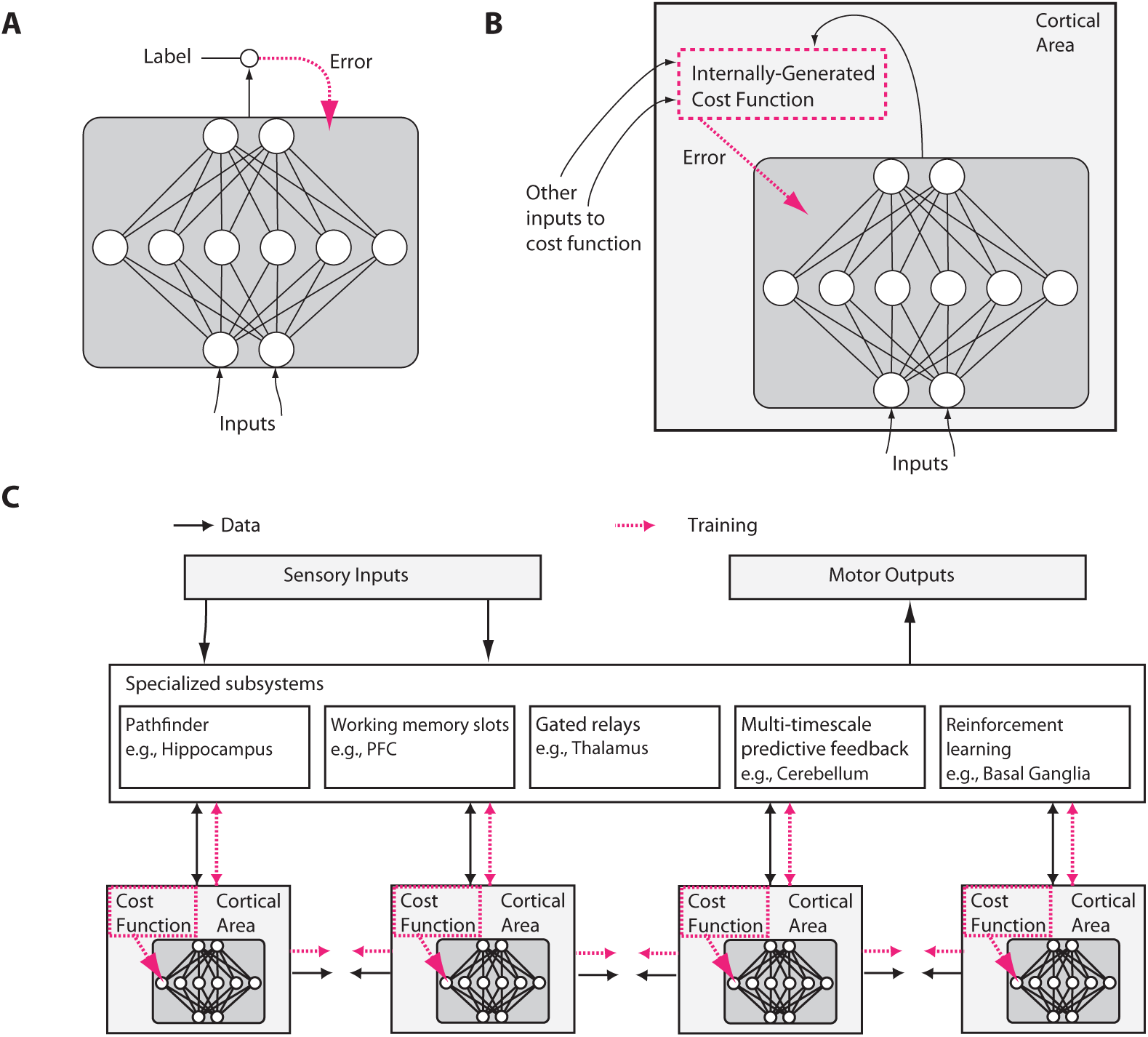
Putative differences between conventional and brain-like neural network designs. **A**) In conventional deep learning, supervised training is based on externally-supplied, labeled data. **B**) In the brain, supervised training of networks can still occur via gradient descent on an error signal, but this error signal must arise from internally generated cost functions. These cost functions are themselves computed by neural modules specified by both genetics and learning. Internally generated cost functions create heuristics that are used to bootstrap more complex learning. For example, an area which recognizes faces might first be trained to detect faces using simple heuristics, like the presence of two dots above a line, and then further trained to discriminate salient facial expressions using representations arising from unsupervised learning and error signals from other brain areas related to social reward processing. **C**) Internally generated cost functions and error-driven training of cortical deep networks form part of a larger architecture containing several specialized systems. Although the trainable cortical areas are schematized as feedforward neural networks here, LSTMs or other types of recurrent networks may be a more accurate analogy, and many neuronal and network properties such as spiking, dendritic computation, neuromodulation, adaptation and homeostatic plasticity, timing-dependent plasticity, direct electrical connections, transient synaptic dynamics, excitatory/inhibitory balance, spontaneous oscillatory activity, axonal conduction delays (Izhikevich, 2006) and others, will influence what and how such networks learn.

Second, recent work in machine learning has started to introduce complex cost functions, those that are not uniform across layers and time, and those that arise from interactions between different parts of a network. For example, introducing the objective of temporal coherence for lower layers (non-uniform cost function over space) improves feature learning (Sermanet and Kavukcuoglu, 2013), cost function schedules (non-uniform cost function over time) improve^1^ generalization (Saxe et al., 2013; Goodfellow et al., 2014b; Gülçehre and Bengio, 2016) and adversarial networks – an example of a cost function arising from internal interactions – allow gradient-based training of generative models (Goodfellow et al., 2014a)^2^. Networks that are easier to train are being used to provide “hints” to help bootstrap the training of more powerful networks (Romero et al., 2014).

Third, machine learning has also begun to diversify the architectures that are subject to optimization. It has introduced simple memory cells with multiple persistent states (Hochreiter and Schmidhuber, 1997; Chung et al., 2014), more complex elementary units such as “capsules” and other structures (Hinton et al., 2011; Livni et al., 2013; Delalleau and Bengio, 2011; Tang et al., 2012), content addressable (Weston et al., 2014; Graves et al., 2014) and location addressable memories (Graves et al., 2014), as well as pointers (Kurach et al., 2015) and hard-coded arithmetic operations (Neelakantan et al., 2015).

These three ideas have, so far, not received much attention in neuroscience. We thus formulate these ideas as three hypotheses about the brain, examine evidence for them, and sketch how experiments could test them. But first, let us state the hypotheses more precisely.

### 1.1 *Hypothesis 1* – The brain optimizes cost functions

The central hypothesis for linking the two fields is that biological systems, like many machine-learning systems, are able to optimize cost functions. The idea of cost functions means that neurons in a brain area can somehow change their properties, e.g., the properties of their synapses, so that they get better at doing whatever the cost function defines as their role. Human behavior sometimes approaches optimality in a domain, e.g., during movement (Körding, 2007), which suggests that the brain may have learned optimal strategies. Subjects minimize energy consumption of their movement system (Taylor and Faisal, 2011), and minimize risk and damage to their body, while maximizing financial and movement gains. Computationally, we now know that optimization of trajectories gives rise to elegant solutions for very complex motor tasks (Mordatch et al., 2012; Todorov and Jordan, 2002; Harris and Wolpert, 1998). We suggest that cost function optimization occurs much more generally in shaping the internal representations and processes used by the brain. Importantly, we also suggest that this requires the brain to have mechanisms for efficient credit assignment in multilayer and recurrent networks.

### 1.2 *Hypothesis 2* – Cost functions are diverse across areas and change over development

A second realization is that cost functions need not be global. Neurons in different brain areas may optimize different things, e.g., the mean squared error of movements, surprise in a visual stimulus, or the allocation of attention. Importantly, such a cost function could be locally generated. For example, neurons could locally evaluate the quality of their statistical model of their inputs (**Figure 1B**). Alternatively, cost functions for one area could be generated by another area. Moreover, cost functions may change over time, e.g., guiding young humans to understanding simple visual contrasts early on, and faces a bit later^3^. This could allow the developing brain to bootstrap more complex knowledge based on simpler knowledge. Cost functions in the brain are likely to be complex and to be arranged to vary across areas and over development.

### 1.3 *Hypothesis 3* – Specialized systems allow efficient solution of key computational problems

A third realization is that structure matters. The patterns of information flow seem fundamentally different across brain areas, suggesting that they solve distinct computational problems. Some brain areas are highly recurrent, perhaps making them predestined for short-term memory storage (Wang, 2012). Some areas contain cell types that can switch between qualitatively different states of activation, such as a persistent firing mode versus a transient firing mode, in response to particular neurotransmitters (Hasselmo, 2006). Other areas, like the thalamus appear to have the information from other areas flowing through them, perhaps allowing them to determine information routing (Sherman, 2005). Areas like the basal ganglia are involved in reinforcement learning and gating of discrete decisions (Sejnowski and Poizner, 2014; Doya, 1999). As every programmer knows, specialized algorithms matter for efficient solutions to computational problems, and the brain is likely to make good use of such specialization (**Figure 1C**).

These ideas are inspired by recent advances in machine learning, but we also propose that the brain has major differences from any of today’s machine learning techniques. In particular, the world gives us a relatively limited amount of information that we could use for supervised learning (Fodor and Crowther, 2002). There is a huge amount of information available for unsupervised learning, but there is no reason to assume that a *generic* unsupervised algorithm, no matter how powerful, would learn the precise things that humans need to know, in the order that they need to know it. The evolutionary challenge of making unsupervised learning solve the “right” problems is, therefore, to find a sequence of cost functions that will deterministically build circuits and behaviors according to prescribed developmental stages, so that in the end a relatively small amount of information suffices to produce the right behavior. For example, a developing duck imprints (Tinbergen, 1965) a template of its parent, and then uses that template to generate goal-targets that help it develop other skills like foraging.

Generalizing from this and from other studies (Ullman et al., 2012; Minsky, 1977), we propose that many of the brain’s cost functions arise from such an internal bootstrapping process. Indeed, we propose that biological development and reinforcement learning can, in effect, program the emergence of a sequence of cost functions that precisely anticipates the future needs faced by the brain’s internal subsystems, as well as by the organism as a whole. This type of developmentally programmed bootstrapping generates an internal infrastructure of cost functions which is diverse and complex, while simplifying the learning problems faced by the brain’s internal processes. Beyond simple tasks like familial imprinting, this type of bootstrapping could extend to higher cognition, e.g., internally generated cost functions could train a developing brain to properly access its memory or to organize its actions in ways that will prove to be useful later on. The potential bootstrapping mechanisms that we will consider operate in the context of unsupervised and reinforcement learning, and go well beyond the types of curriculum learning ideas used in today’s machine learning (Bengio et al., 2009).

In the rest of this paper, we will elaborate on these hypotheses. First, we will argue that both local and multi-layer optimization is, perhaps surprisingly, compatible with what we know about the brain. Second, we will argue that cost functions differ across brain areas and change over time and describe how cost functions interacting in an orchestrated way could allow bootstrapping of complex function. Third, we will list a broad set of specialized problems that need to be solved by neural computation, and the brain areas that have structure that seems to be matched to a particular computational problem. We then discuss some implications of the above hypotheses for research approaches in neuroscience and machine learning, and sketch a set of experiments to test these hypotheses. Finally, we discuss this architecture from the perspective of evolution.

## 2. The brain can optimize cost functions

Much of machine learning is based on efficiently optimizing functions, and, as we will detail below, the ability to use backpropagation of error (Werbos, 1974; Rumelhart et al., 1986) to calculate gradients of arbitrary parametrized functions has been a key breakthrough. In **Hypothesis 1**, we claim that the brain is also, at least in part^4^, an optimization machine. But what exactly does it mean to say that the brain can optimize cost functions? After all, many processes can be viewed as optimizations. For example, the laws of physics are often viewed as minimizing an action functional, while evolution optimizes the fitness of replicators over a long timescale. To be clear, our main claims are: that **a)** the brain has powerful mechanisms for credit assignment during learning that allow it to optimize global functions in multi-layer networks by adjusting the properties of each neuron to contribute to the global outcome, and that **b)** the brain has mechanisms to specify exactly which cost functions it subjects its networks to, i.e., that the cost functions are highly tunable, shaped by evolution and matched to the animal’s ethological needs. Thus, the brain uses cost functions as a key driving force of its development, much as modern machine learning systems do.

To understand the basis of these claims, we must now delve into the details of how the brain might efficiently perform credit assignment throughout large, multi-layered networks, in order to optimize complex functions. We argue that the brain uses several different types of optimization to solve distinct problems. In some structures, it may use genetic pre-specification of circuits for problems that require only limited learning based on data, or it may exploit local optimization to avoid the need to assign credit through many layers of neurons. It may also use a host of proposed circuit structures that would allow it to actually perform, in effect, backpropagation of errors through a multi-layer network, using biologically realistic mechanisms – a feat that had once been widely believed to be biologically implausible (Crick, 1989; Stork, 1989). Potential such mechanisms include circuits that literally backpropagate error derivatives in the manner of conventional backpropagation, as well as circuits that provide other efficient means of approximating the effects of backpropagation, i.e., of rapidly computing the approximate gradient of a cost function relative to any given connection weight in the network. Lastly, the brain may use algorithms that exploit specific aspects of neurophysiology – such as spike timing dependent plasticity, dendritic computation, local excitatory-inhibitory networks, or other properties – as well as the integrated nature of higher-level brain systems. Such mechanisms promise to allow learning capabilities that go even beyond those of current backpropagation networks.

### 2.1 Local self-organization and optimization without multi-layer credit assignment

Not all learning requires a general-purpose optimization mechanism like gradient descent^5^. Many theories of cortex (George and Hawkins, 2009; Kappel et al., 2014) emphasize potential self-organizing and unsupervised learning properties that may obviate the need for multi-layer backpropagation as such. Hebbian plasticity, which adjusts weights according to correlations in pre-synaptic and post-synaptic activity, is well established^6^. Various versions of Hebbian plasticity (Miller and MacKay, 1994), e.g., with nonlinearities (Brito and Gerstner, 2016), can give rise to different forms of correlation and competition between neurons, leading to the self-organized formation of ocular dominance columns, self-organizing maps and orientation columns (Ferster and Miller, 2003; Miller et al., 1989). Often these types of local self-organization can also be viewed as optimizing a cost function: for example, certain forms of Hebbian plasticity can be viewed as extracting the principal components of the input, which minimizes a reconstruction error (Pehlevan and Chklovskii, 2015).

To generate complex temporal patterns, the brain may also implement other forms of learning that do not require any equivalent of full back-propagation through a multilayer network. For example, “liquid-” (Maass et al., 2002) or “echo-state machines” (Jaeger and Haas, 2004) are randomly connected recurrent networks that form a basis set (also known as a “reservoir”) of random filters, which can be harnessed for learning with tunable readout weights. Variants exhibiting chaotic, spontaneous dynamics can even be trained by feeding back readouts into the network and suppressing the chaotic activity (Sussillo and Abbott, 2009). Learning only the readout layer makes the optimization problem much simpler (indeed, equivalent to regression for supervised learning). Additionally, echo state networks can be trained by reinforcement learning as well as supervised learning (Bush, 2007; Hoerzer et al., 2014). Reservoirs of random nonlinear filters are one interpretation of the diverse, high-dimensional, mixed-selectivity tuning properties of many neurons, e.g. in the prefrontal cortex (Enel et al., 2016). Other variants of learning rules that modify only a fraction of the synapses inside a random network are being developed as models of biological working memory and sequence generation (Rajan et al., 2016).

### 2.2 Biological implementation of optimization

We argue that the above mechanisms of local self-organization are likely insufficient to account for the brain’s powerful learning performance (Brea and Gerstner, 2016). To elaborate on the need for an efficient means of gradient computation in the brain, we will first place backpropagation into its computational context (Hinton, 1989; Baldi and Sadowski, 2015). Then we will explain how the brain could plausibly implement approximations of gradient descent.

#### 2.2.1 The need for efficient gradient descent in multi-layer networks

The simplest mechanism to perform cost function optimization is sometimes known as the “twiddle” algorithm or, more technically, as “serial perturbation”. This mechanism works by perturbing (i.e., “twiddling”), with a small increment, a single weight in the network, and verifying improvement by measuring whether the cost function has decreased compared to the network’s performance with the weight unperturbed. If improvement is noticeable, the perturbation is used as a direction of change to the weight; otherwise, the weight is changed in the opposite direction (or not changed at all). Serial perturbation is therefore a method of “coordinate descent” on the cost, but it is slow and requires global coordination: each synapse in turn is perturbed while others remain fixed.

Weight perturbation (or parallel perturbation) perturbs all of the weights in the network at once. It is able to optimize small networks to perform tasks but generally suffers from high variance. That is, the measurement of the gradient direction is noisy and changes drastically from perturbation to perturbation because a weight’s influence on the cost is masked by the changes of all other weights, and there is only one scalar feedback signal indicating the change in the cost^7^. Weight perturbation is dramatically inefficient for large networks. In fact, parallel and serial perturbation learn at approximately the same rate if the time measure counts the number of times the network propagates information from input to output (Werfel et al., 2005).

Some efficiency gain can be achieved by perturbing neural activities instead of synaptic weights, acknowledging the fact that any long-range effect of a synapse is mediated through a neuron. Like weight perturbation and unlike serial perturbation, minimal global coordination is needed: each neuron only needs to receive a feedback signal indicating the global cost. The variance of node perturbation’s gradient estimate is far smaller than that of weight perturbation under the assumptions that either all neurons or all weights, respectively, are perturbed and that they are perturbed at the same frequency. In this case, node perturbation’s variance is proportional to the number of cells in the network, not the number of synapses.

All of these approaches are slow either due to the time needed for serial iteration over all weights or the time needed for averaging over low signal-to-noise ratio gradient estimates. To their credit however, none of these approaches requires more than knowledge of local activities and the single global cost signal. Real neural circuits in the brain have mechanisms (e.g., diffusible neuromodulators) that appear to code the signals relevant to implementing those algorithms. In many cases, for example in reinforcement learning, the cost function, which is computed based on interaction with an unknown environment, cannot be differentiated directly, and an agent has no choice but to deploy clever twiddling to explore at some level of the system (Williams, 1992).

Backpropagation, in contrast, works by computing the sensitivity of the cost function to each weight based on the layered structure of the system. The derivatives of the cost function with respect to the last layer can be used to compute the derivatives of the cost function with respect to the penultimate layer, and so on, all the way down to the earliest layers^8^. Backpropagation can be computed rapidly, and for a single input-output pattern, it exhibits no variance in its gradient estimate. The backpropagated gradient has no more noise for a large system than for a small system, so deep and wide architectures with great computational power can be trained efficiently.

#### 2.2.2 Biologically plausible approximations of gradient descent

To permit biological learning with efficiency approaching that of machine learning methods, some provision for more sophisticated gradient propagation may be suspected. Contrary to what was once a common assumption, there are now many proposed “biologically plausible” mechanisms by which a neural circuit could implement optimization algorithms that, like back-propagation, can efficiently make use of the gradient. These include Generalized Recirculation (O’Reilly, 1996), Contrastive Hebbian Learning (Xie and Seung, 2003), random feedback weights together with synaptic homeostasis (Lillicrap et al., 2014; Liao et al., 2015), spike timing dependent plasticity (STDP) with iterative inference and target propagation (Scellier and Bengio, 2016; Bengio et al., 2015a), complex neurons with backpropagating action-potentials (Körding and König, 2000), and others (Balduzzi et al., 2014). While these mechanisms differ in detail, they all invoke feedback connections that carry error phasically. Learning occurs by comparing a prediction with a target, and the prediction error is used to drive top-down changes in bottom-up activity.

As an example, consider O’Reilly’s temporally eXtended Contrastive Attractor Learning (XCAL) algorithm (O’Reilly et al., 2012, 2014b). Suppose we have a multilayer neural network with an input layer, an output layer, and a set of hidden layers in between. O’Reilly showed that the same functionality as backpropagation can be implemented by a bidirectional network with the same weights but symmetric connections. After computing the outputs using the forward connections only, we set the output neurons to the values they should have. The dynamics of the network then cause the hidden layers’ activities to evolve toward a stable attractor state linking input to output. The XCAL algorithm performs a type of local modified Hebbian learning at each synapse in the network during this process (O’Reilly et al., 2012). The XCAL Hebbian learning rule compares the local synaptic activity (pre x post) during the early phase of this settling (before the attractor state is reached) to the final phase (once the attractor state has been reached), and adjusts the weights in a way that should make the early phase reflect the later phase more closely. These contrastive Hebbian learning methods even work when the connection weights are not precisely symmetric (O’Reilly, 1996). XCAL has been implemented in biologically plausible conductance-based neurons and basically implements the backpropagation of error approach.

Approximations to backpropagation could also be enabled by the millisecond-scale timing of of neural activities (O’Reilly et al., 2014b). Spike timing dependent plasticity (STDP) (Markram et al., 1997), for example, is a feature of some neurons in which the sign of the synaptic weight change depends on the precise millisecond-scale relative timing of pre-synaptic and post-synaptic spikes. This is conventionally interpreted as Hebbian plasticity that measures the potential for a causal relationship between the pre-synaptic and post-synaptic spikes: a pre-synaptic spike could have contributed to causing a post-synaptic spike only if it occurs shortly beforehand^9^. To enable a backpropagation mechanism, Hinton has suggested an alternative interpretation: that neurons could encode the types of error derivatives needed for backpropagation in the temporal derivatives of their firing rates (Hinton, 2007, 2016). STDP then corresponds to a learning rule that is sensitive to these error derivatives (Bengio et al., 2015b; Xie and Seung, 2000). In other words, in an appropriate network context, STDP learning could give rise to a biological implementation of backpropagation^10^.

Another possible mechanism, by which biological neural networks could approximate back-propagation, is “feedback alignment” (Lillicrap et al., 2014; Liao et al., 2015). There, the feedback pathway in backpropagation, by which error derivatives at a layer are computed from error derivatives at the subsequent layer, is replaced by a set of random feedback connections, with no dependence on the forward weights. Subject to the existence of a synaptic normalization mechanism and approximate sign-concordance between the feedforward and feedback connections (Liao et al., 2015), this mechanism of computing error derivatives works nearly as well as backpropagation on a variety of tasks. In effect, the forward weights are able to adapt to bring the network into a regime in which the random backwards weights actually carry the information that is useful for approximating the gradient. This is a remarkable and surprising finding, and is indicative of the fact that our understanding of gradient descent optimization, and specifically of the mechanisms by which backpropagation itself functions, are still incomplete. In neuroscience, meanwhile, we find feedback connections almost wherever we find feed-forward connections, and their role is the subject of diverse theories (Callaway, 2004; Maass et al., 2007). It should be noted that feedback alignment as such does not specify exactly how neurons represent and make use of the error signals; it only relaxes a constraint on the transport of the error signals. Thus, feedback alignment is more a primitive that can be used in fully biological (approximate) implementations of backpropagation, than a fully biological implementation in its own right. As such, it may be possible to incorporate it into several of the other schemes discussed here.

The above “biological” implementations of backpropagation still lack some key aspects of biological realism. For example, in the brain, neurons tend to be either excitatory or inhibitory but not both, whereas in artificial neural networks a single neuron may send both excitatory and inhibitory signals to its downstream neurons. Fortunately, this constraint is unlikely to limit the functions that can be learned (Parisien et al., 2008; Tripp and Eliasmith, 2016). Other biological considerations, however, need to be looked at in more detail: the highly recurrent nature of biological neural networks, which show rich dynamics in time, and the fact that most neurons in mammalian brains communicate via spikes. We now consider these two issues in turn.

**Temporal credit assignment:** The biological implementations of backpropagation proposed above, while applicable to feedforward networks, do not give a natural implementation of “backpropagation through time” (BPTT) (Werbos, 1990) for recurrent networks, which is widely used in machine learning for training recurrent networks on sequential processing tasks. BPTT “unfolds” a recurrent network across multiple discrete time steps and then runs backpropagation on the unfolded network to assign credit to particular units at particular time steps^11^. While the network unfolding procedure of BPTT itself does not seem biologically plausible, to our intuition, it is unclear to what extent temporal credit assignment is truly needed (Ollivier and Charpiat, 2015) for learning particular temporally extended tasks.

If the system is given access to appropriate memory stores and representations (Buonomano and Merzenich, 1995; Gershman et al., 2012, 2014) of temporal context, this could potentially mitigate the need for temporal credit assignment as such – in effect, memory systems could “spatialize” the problem of temporal credit assignment^12^. For example, memory networks (Weston et al., 2014) store everything by default up to a certain buffer size, eliminating the need to perform credit assignment over the write-to-memory events, such that the network only needs to perform credit assignment over the read-from-memory events. In another example, certain network architectures that are superficially very deep, but which possess particular types of “skip connections”, can actually be seen as ensembles of comparatively shallow networks (Veit et al., 2016); applied in the time domain, this could limit the need to propagate errors far backwards in time. Other, similar specializations or higher-levels of structure could, potentially, further ease the burden on credit assignment.

Can generic recurrent networks perform temporal credit assignment in in a way that is more biologically plausible than BPTT? Indeed, new discoveries are being made about the capacity for supervised learning in continuous-time recurrent networks with more realistic synapses and neural integration properties. In internal FORCE learning (Sussillo and Abbott, 2009), internally generated random fluctuations inside a chaotic recurrent network are adjusted to provide feedback signals that drive weight changes internal to the network while the outputs are clamped to desired patterns. This is made possible by a learning procedure that rapidly adjusts the network output to a state where it is close to the clamped values, and exerts continuous control to keep this difference small throughout the learning process^13^. This procedure is able to control and exploit the chaotic dynamical patterns that are spontaneously generated by the network.

Werbos has proposed in his “error critic” that an online approximation to BPTT can be achieved by learning to predict the backward-through-time gradient signal (costate) in a manner analogous to the prediction of value functions in reinforcement learning (Si, 2004). This kind of idea was recently applied in (Jaderberg et al., 2016) to allow decoupling of different parts of a network during training and to facilitate back-propagation through time. Broadly, we are only beginning to understand how neural activity can itself represent the time variable (Finnerty and Shadlen, 2015; Xu et al., 2014)^14^, and how recurrent networks can learn to generate trajectories of population activity over time (Liu and Buonomano, 2009). Moreover, as we discuss below, a number of cortical models also propose means, other than BPTT, by which networks could be trained on sequential prediction tasks, even in an online fashion (Cui et al., 2015; O’Reilly et al., 2014b; Brea et al., 2016). A broad range of ideas can be used to approximate BPTT in more realistic ways.

**Spiking networks:** It has been difficult to apply gradient descent learning directly to spiking neural networks^15^,^16^, although there do exist learning rules for doing so in specific representational contexts and network structures (Bekolay et al., 2013). A number of optimization procedures have been used to generate, indirectly, spiking networks which can perform complex tasks, by performing optimization on a continuous representation of the network dynamics and embedding variables into high-dimensional spaces with many spiking neurons representing each variable (Abbott et al., 2016; DePasquale et al., 2016; Komer and Eliasmith, 2016; Thalmeier et al., 2015). The use of recurrent connections with multiple timescales can remove the need for backpropagation in the direct training of spiking recurrent networks (Bourdoukan and Denève, 2015). Fast connections maintain the network in a state where slow connections have local access to a global error signal. While the biological realism of these methods is still unknown, they all allow connection weights to be learned in spiking networks.

These and other novel learning procedures illustrate the fact that we are only beginning to understand the connections between the temporal dynamics of biologically realistic networks, and mechanisms of temporal and spatial credit assignment. Nevertheless, we argue here that existing evidence suggests that biologically plausible neural networks can solve these problems – in other words, it is possible to efficiently optimize complex functions of temporal history in the context of spiking networks of biologically realistic neurons. In any case, there is little doubt that spiking recurrent networks using realistic population coding schemes can, with an appropriate choice of connection weights, compute complicated, cognitively relevant functions^17^. The question is how the developing brain efficiently learns such complex functions.

### 2.3 Other principles for biological learning

The brain has mechanisms and structures that could support learning mechanisms different from typical gradient-based optimization algorithms employed in artificial neural networks.

#### 2.3.1 Exploiting biological neural mechanisms

The complex physiology of individual biological neurons may not only help explain how some form of efficient gradient descent could be implemented within the brain, but also could provide mechanisms for learning that go beyond back-propagation. This suggests that the brain may have discovered mechanisms of credit assignment quite different from those dreamt up by machine learning.

One such biological primitive is dendritic computation, which could impact prospects for learning algorithms in several ways. First, real neurons are highly nonlinear (Antic et al., 2010), with the dendrites of each *single* neuron implementing^18^ something computationally similar to a three-layer neural network (Mel, 1992)^19^. Individual neurons thus should not be regarded as single “nodes” but as multi-component subnetworks. Second, when a neuron spikes, its action potential propagates back from the soma into the dendritic tree. However, it propagates more strongly into the branches of the dendritic tree that have been active (Williams and Stuart, 2000), potentially simplifying the problem of credit assignment (Körding and König, 2000). Third, neurons can have multiple somewhat independent dendritic compartments, as well as a somewhat independent somatic compartment, which means that the neuron should be thought of as storing more than one variable. Thus, there is the possibility for a neuron to store both its activation itself, and the error derivative of a cost function with respect to its activation, as required in backpropagation, and biological implementations of backpropagation based on this principle have been proposed (Körding and König, 2001; Schiess et al., 2016)^20^. Overall, the implications of dendritic computation for credit assignment in deep networks are only beginning to be considered^21^. But it is clear that the types of bi-directional, non-linear, multi-variate interactions that are possible *inside* a single neuron could support gradient descent learning or other powerful optimization mechanisms.

Beyond dendritic computation, diverse mechanisms (Marblestone and Boyden, 2014) like retrograde (post-synaptic to pre-synaptic) signals using cannabinoids (Wilson and Nicoll, 2001), or rapidly-diffusing gases such as nitric oxide (Arancio et al., 1996), are among many that could enable learning rules that go beyond conventional conceptions of backpropagation. Harris has suggested (Harris, 2008; Lewis and Harris, 2014) how slow, retroaxonal (i.e., from the outgoing synapses back to the parent cell body) transport of molecules like neurotrophins could allow neural networks to implement an analog of an exchangeable currency in economics, allowing networks to self-organize to efficiently provide information to downstream “consumer” neurons that are trained via faster and more direct error signals. The existence of these diverse mechanisms may call into question traditional, intuitive notions of “biological plausibility” for learning algorithms.

Another potentially important biological primitive is neuromodulation. The same neuron or circuit can exhibit different input-output responses and plasticity depending on a global circuit state, as reflected by the concentrations of various *neuromodulators* like dopamine, serotonin, norepinephrine, acetylcholine, and hundreds of different neuropeptides such as opiods (Bargmann and Marder, 2013; Bargmann, 2012). These modulators interact in complex and cell-type-specific ways to influence circuit function. Interactions with glial cells also play a role in neural signaling and neuromodulation, leading to the concept of “tripartite” synapses that include a glial contribution (Perea et al., 2009). Modulation could have many implications for learning. First, modulators can be used to gate synaptic plasticity on and off selectively in different areas and at different times, allowing precise, rapidly updated orchestration of where and when cost functions are applied. Furthermore, it has been argued that a single neural circuit can be thought of as multiple overlapping circuits with modulation switching between them (Bargmann and Marder, 2013; Bargmann, 2012). In a learning context, this could potentially allow sharing of synaptic weight information between overlapping circuits. Dayan (2012) discusses further computational aspects of neuromodulation. Overall, neuromodulation seems to expand the range of possible algorithms that could be used for optimization.

#### 2.3.2 Learning in the cortical sheet

A number of models attempt to explain cortical learning on the basis of specific architectural features of the 6-layered cortical sheet. These models generally agree that a primary function of the cortex is some form of unsupervised learning via prediction (O’Reilly et al., 2014b; Brea et al., 2016)^22^. Some cortical learning models are explicit attempts to map cortical structure onto the framework of message-passing algorithms for Bayesian inference (George and Hawkins, 2009; Dean, 2005; Lee and Mumford, 2003), while others start with particular aspects of cortical neurophysiology and seek to explain those in terms of a learning function, or in terms of a computational function, e.g., hierarchical clustering (Rodriguez et al., 2004). For example, the nonlinear and dynamical properties of cortical pyramidal neurons – the principal excitatory neuron type in cortex (Shepherd, 2014) – are of particular interest here, especially because these neurons have multiple dendritic zones that are targeted by different kinds of projections, which may allow the pyramidal neuron to make comparisons of top-down and bottom-up inputs^23^.

Other aspects of the laminar cortical architecture could be crucial to how the brain implements learning. Local inhibitory neurons targeting particular dendritic compartments of the L5 pyramidal could be used to exert precise control over when and how the relevant feedback signals and associative mechanisms are utilized. Notably, local inhibitory networks could also give rise to competition (Petrov et al., 2010) between different representations in the cortex, perhaps allowing one cortical column to suppress others nearby, or perhaps even to send more sophisticated messages to gate the state transitions of its neighbors (Bach and Herger, 2015). Moreover, recurrent connectivity with the thalamus, structured bursts of spiking, and cortical oscillations (not to mention other mechanisms like neuromodulation) could control the storage of information over time, to facilitate learning based on temporal prediction. These concepts begin to suggest preliminary, exploratory models for how the detailed anatomy and physiology of the cortex could be interpreted within a machine-learning framework that goes beyond backpropagation. But these are early days: we still lack detailed structural/molecular and functional maps of even a single local cortical microcircuit.

#### 2.3.3 One-shot learning

Human learning is often one-shot: it can take just a single exposure to a stimulus to never forget it, as well as to generalize from it to new examples. One way of allowing networks to have such properties is what is described by I-theory, in the context of learning invariant representations for object recognition (Anselmi et al., 2015). Instead of training via gradient descent, image templates are stored in the weights of simple-complex cell networks while objects undergo transformations, similar to the use of stored templates in HMAX (Serre et al., 2007). The theories then aim to show that you can invariantly and discriminatively represent objects using a single sample, even of a new class (Anselmi et al., 2015)^24^.

Additionally, the nervous system may have a way of quickly storing and replaying sequences of events. This would allow the brain to move an item from episodic memory into a long-term memory stored in the weights of a cortical network (Ji and Wilson, 2007), by replaying the memory over and over. This solution effectively uses many iterations of weight updating to fully learn a single item, even if one has only been exposed to it once. Alternatively, the brain could rapidly store an episodic memory and then retrieve it later without the need to perform slow gradient updates, which has proven to be useful for fast reinforcement learning in scenarios with limited available data (Blundell et al., 2016).

Finally, higher-level systems in the brain may be able to implement Bayesian learning of sequential programs, which is a powerful means of one-shot learning (Lake et al., 2015). This type of cognition likely relies on an interaction between multiple brain areas such as the prefrontal cortex and basal ganglia.

These potential substrates of one-shot learning rely on mechanisms other than simple gradient descent. It should be noted, though, that recent architectural advances, including specialized spatial attention and feedback mechanisms (Rezende et al., 2016), as well as specialized memory mechanisms (Santoro et al., 2016), do allow some types of one-shot generalization to be driven by backpropagation-based learning.

#### 2.3.4 Active learning

Human learning is often active and deliberate. It seems likely that, in human learning, actions are chosen so as to generate interesting training examples, and sometimes also to test specific hypotheses. Such ideas of active learning and “child as scientist” go back to Piaget and have been elaborated more recently (Gopnik et al., 2000). We want our learning to be based on maximally informative samples, and active querying of the environment (or of internal subsystems) provides a way route to this.

At some level of organization, of course, it would seem useful for a learning system to develop explicit representations of its uncertainty, since this can be used to guide the system to actively seek the information that would reduce its uncertainty most quickly. Moreover, there are population coding mechanisms that could support explicit probabilistic computations (Ma et al., 2006; Zemel and Dayan, 1997; Gershman and Beck, 2016; Eliasmith and Martens, 2011; Rao, 2004; Sahani and Dayan, 2003). Yet it is unclear to what extent and at what levels the brain uses an explicitly probabilistic framework, or to what extent probabilistic computations are emergent from other learning processes (Orhan and Ma, 2016)^25^ ^26^.

Standard gradient descent does not incorporate any such adaptive sampling mechanism, e.g., it does not deliberately sample data so as to maximally reduce its uncertainty. Interestingly, however, stochastic gradient descent can be used to generate a system that samples adaptively (Bouchard et al., 2015; Alain et al., 2015). In other words, a system can learn, by gradient descent, how to choose its own input data samples in order to learn most quickly from them by gradient descent.

Ideally, the learner learns to choose actions that will lead to the largest improvements in its prediction or data compression performance (Schmidhuber, 2010). In (Schmidhuber, 2010), this is done in the framework of reinforcement learning, and incorporates a mechanisms for the system to measure its own rate of learning. In other words, it is possible to reinforcement-learn a policy for selecting the most interesting inputs to drive learning. Adaptive sampling methods are also known in reinforcement learning that can achieve optimal Bayesian exploration of Markov Decision Process environments (Guez et al., 2012; Sun et al., 2011).

These approaches achieve optimality in an arbitrary, abstract environment. But of course, evolution may also encode its implicit knowledge of the organism’s natural environment, the behavioral goals of the organism, and the developmental stages and processes which occur inside the organism, as priors or heuristics^27^ which would further constrain the types of adaptive sampling that are optimal in practice. For example, simple heuristics like seeking certain perceptual signatures of novelty, or more complex heuristics like monitoring situations that other people seem to find interesting, might be good ways to bias sampling of the environment so as to learn more quickly. Other such heuristics might be used to give internal brain systems the types of training data that will be most useful to those particular systems at any given developmental stage.

We are only beginning to understand how active learning might be implemented in the brain. We speculate that multiple mechanisms, specialized to different brain systems and spatio-temporal scales, could be involved. The above examples suggest that at least some such mechanisms could be understood from the perspective of optimizing cost functions.

### 2.4 Differing biological requirements for supervised and reinforcement learning

We have suggested ways in which the brain could implement learning mechanisms of comparable power to backpropagation. But in many cases, the system may be more limited by the available training signals than by the optimization process itself. In machine learning, one distinguishes supervised learning, reinforcement learning and unsupervised learning, and the training data limitation manifests differently in each case.

Both supervised and reinforcement learning require some form of teaching signal, but the nature of the teaching signal in supervised learning is different from that in reinforcement learning. In supervised learning, the trainer provides the entire vector of errors for the output layer and these are back-propagated to compute the gradient: a locally optimal direction in which to update all of the weights of a potentially multilayer and/or recurrent network. In reinforcement learning, however, the trainer provides a scalar evaluation signal, but this is not sufficient to derive a low-variance gradient. Hence, some form of trial and error twiddling must be used to discover how to increase the evaluation signal. Consequently, reinforcement learning is generally much less efficient than supervised learning.

Reinforcement learning in shallow networks is simple to implement biologically. For reinforcement learning of a deep network to be biologically plausible, however, we need a more powerful learning mechanism, since we are learning based on a more limited evaluation signal than in the supervised case: we do not have the full target pattern to train towards. Nevertheless, approximations of gradient descent can be achieved in this case, and there are cases in which the scalar evaluation signal of reinforcement learning can be used to efficiently update a multi-layer network by gradient descent. The “attention-gated reinforcement learning” (AGREL) networks of (Stnior et al., 2013; Brosch et al., 2015; Roelfsema and van Ooyen, 2005), and variants like KickBack (Balduzzi, 2014), give a way to compute an approximation to the full gradient in a reinforcement learning context using a feedback-based attention mechanism for credit assignment within the multi-layer network. The feedback pathway, together with a diffusible reward signal, together gate plasticity. For networks with more than three layers, this gives rise to a model based on columns containing parallel feedforward and feedback pathways (Roelfsema and van Ooyen, 2005), and for recurrent networks that settle into attractor states it gives a reinforcement-trained version (Brosch et al., 2015) of the Almeida/Pineda recurrent backpropagation algorithm (Pineda, 1987). The process is still not as efficient or generic as backpropagation, but it seems that this form of feedback can make reinforcement learning in multi-layer networks more efficient than a naive node perturbation or weight perturbation approach.

The machine-learning field has recently been tackling the question of credit assignment in deep reinforcement learning. Deep Q-learning (Mnih et al., 2015) demonstrates reinforcement learning in a deep network, wherein most of the network is trained via backpropagation. In regular Q learning, we define a function Q, which estimates the best possible sum of future rewards (the return) if we are in a given state and take a given action. In deep Q learning, this function is approximated by a neural network that, in effect, estimates action-dependent returns in a given state. The network is trained using backpropagation of local errors in Q estimation, using the fact that the return decomposes into the current reward plus the discounted estimate of future return at the next moment. During training, as the agent acts in the environment, a series of loss functions is generated at each step, defining target patterns that can be used as the supervision signal for back-propagation. As Q is a highly nonlinear function of the state, tricks are needed to make deep Q learning efficient and stable, including experience replay and a particular type of mini-batch training. It is also necessary to store the outputs from the previous iteration (or clone the entire network) in evaluating the loss function for the subsequent iteration^28^.

This process for generating learning targets provides a kind of bridge between reinforcement learning and efficient backpropagation-based gradient descent learning^29^. Importantly, only temporally local information is needed making the approach relatively compatible with what we know about the nervous system.

Even given these advances, a key remaining issue in reinforcement learning is the problem of long timescales, e.g., learning the many small steps needed to navigate from London to Chicago. Many of the formal guarantees of reinforcement learning (Williams and Baird, 1993), for example, suggest that the difference between an optimal policy and the learned policy becomes increasingly loose as the discount factor shifts to take into account reward at longer timescales. Although the degree of optimality of human behavior is unknown, people routinely engage in adaptive behaviors that can take hours or longer to carry out, by using specialized processes like *prospective memory* to “remember to remember” relevant variables at the right times, permitting extremely long timescales of coherent action. Machine learning has not yet developed methods to deal with such a wide range of timescales and scopes of hierarchical action. Below we discuss ideas of hierarchical reinforcement learning that may make use of callable procedures and sub-routines, rather than operating explicitly in a time domain.

As we will discuss below, some form of deep reinforcement learning may be used by the brain for purposes beyond optimizing global rewards, including the training of local networks based on diverse internally generated cost functions. Scalar reinforcement-like signals are easy to compute, and easy to deliver to other areas, making them attractive mechanistically. If the brain does employ internally computed scalar reward-like signals as a basis for cost functions, it seems likely that it will have found an efficient means of reinforcement-based training of deep networks, but it is an open question whether an analog of deep Q networks, AGREL, or some other mechanism entirely, is used in the brain for this purpose. Moreover, as we will discuss further below, it is possible that reinforcement-type learning is made more efficient in the context of specialized brain systems like short term memories, replay mechanisms, and hierarchically organized control systems. These specialized systems could reduce reliance on a need for powerful credit assignment mechanisms for reinforcement learning. Finally, if the brain uses a diversity of scalar reward-like signals to implement different cost functions, then it may need to mediate delivery of those signals via a comparable diversity of molecular substrates. The great diversity of neuromodulatory signals, e.g., neuropeptides, in the brain (Bargmann, 2012; Bargmann and Marder, 2013) makes such diversity quite plausible, and moreover, the brain may have found other, as yet unknown, mechanisms of diversifying reward-like signaling pathways and enabling them to act independently of one another.

## 3. The cost functions are diverse across brain areas and time

In the last section, we argued that the brain can optimize functions. This raises the question of what functions it optimizes. Of course, in the brain, a cost function will itself be created (explicitly or implicitly) by a neural network shaped by the genome. Thus, the cost function used to train a given sub-network in the brain is a key innate property that can be built into the system by evolution. It may be much cheaper in biological terms to specify a cost function that allows the rapid learning of the solution to a problem than to specify the solution itself.

In **Hypothesis 2**, we proposed that the brain optimizes not a single “end-to-end” cost function, but rather a diversity of internally generated cost functions specific to particular brain functions^30^. To understand how and why the brain may use a diversity of cost functions, it is important to distinguish the differing types of cost functions that would be needed for supervised, unsupervised and reinforcement learning. We can also seek to identify types of cost functions that the brain may need to generate from a functional perspective, and how each may be implemented as supervised, unsupervised, reinforcement-based or hybrid systems.

### 3.1 How cost functions may be represented and applied

What additional circuitry is required to actually impose a cost function on an optimizing network? In the most familiar case, supervised learning may rely on computing a vector of errors at the output of a network, which will rely on some comparator circuitry^31^ to compute the difference between the network outputs and the target values. This difference could then be backpropagated to earlier layers. An alternative way to impose a cost function is to “clamp” the output of the network, forcing it to occupy a desired target state. Such clamping is actually assumed in some of the putative biological implementations of back-propagation described above, such as XCAL and target propagation. Alternatively, as described above, scalar reinforcement signals are attractive as internally-computed cost functions, but using them in deep networks requires special mechanisms for credit assignment.

In unsupervised learning, cost functions may not take the form of externally supplied training or error signals, but rather can be built into the dynamics inherent to the network itself, i.e., there may be no need for a *separate* circuit to compute and impose a cost function on the network. For example, specific spike-timing-dependent and homeostatic plasticity rules have been shown to give rise to gradient descent on a prediction error in recurrent neural networks (Galtier and Wainrib, 2013). Thus, specific unsupervised objectives could be implemented implicitly through specific local network dynamics^32^ and plasticity rules inside a network without explicit computation of cost function, nor explicit propagation of error derivatives.

Alternatively, explicit cost functions could be computed, delivered to an optimizing network, and used for unsupervised learning, following a variety of principles being discovered in machine learning (e.g., (Radford et al., 2015; Lotter et al., 2015)). These networks rely on backpropagation as the sole learning rule, and typically find a way to encode the desired cost function into the error derivatives which are backpropagated. For example, prediction errors naturally give rise to error signals for unsupervised learning, as do reconstruction errors in autoencoders, and these error signals can also be augmented with additional penalty or regularization terms that enforce objectives like sparsity or continuity, as described below. Then these error derivatives can be propagated throughout the network via standard back-propagation. In such systems, the objective function and the optimization mechanism can thus be mixed and matched modularly. In the next sections, we elaborate on these and other means of specifying and delivering cost functions in different learning contexts.

### 3.2 Cost functions for unsupervised learning

There are many objectives that can be optimized in an unsupervised context, to accomplish different kinds of functions or guide a network to form particular kinds of representations.

#### 3.2.1 Matching the statistics of the input data using generative models

In one common form of unsupervised learning, higher brain areas attempt to produce samples that are statistically similar to those actually seen in lower layers. For example, the wake-sleep algorithm (Hinton et al., 1995) requires the sleep mode to sample potential data points whose distribution should then match the observed distribution. Unsupervised pre-training of deep networks is an instance of this (Erhan and Manzagol, 2009), typically making use of a stacked auto-encoder framework. Similarly, in target propagation (Bengio, 2014), a top-down circuit, together with lateral information, has to produce data that directs the local learning of a bottom-up circuit and vice-versa. Ladder autoencoders make use of lateral connections and local noise injection to introduce an unsupervised cost function, based on internal reconstructions, that can be readily combined with supervised cost functions defined on the networks top layer outputs (Valpola, 2015). Compositional generative models generate a scene from discrete combinations of template parts and their transformations (Wang and Yuille, 2014), in effect performing a rendering of a scene based on its structural description. Hinton and colleagues have also proposed cortical “capsules” (Tang et al., 2013, 2012; Hinton et al., 2011) for compositional inverse rendering. The network can thus implement a statistical goal that embodies some understanding of the way that the world produces samples^33^.

Learning rules for generative models have historically involved local message passing of a form quite different from backpropagation, e.g., in a multi-stage process that first learns one layer at a time and then fine-tunes via the wake-sleep algorithm (Hinton et al., 2006). Message-passing implementations of probabilistic inference have also been proposed as an explanation and generalization of deep convolutional networks (Patel et al., 2015; Chen et al., 2014). Various mappings of such processes onto neural circuitry have been attempted (Sountsov and Miller, 2015; George and Hawkins, 2009; Lee and Yuille, 2011), and related models (Makin et al., 2013, 2016) have been used to account for optimal multi-sensory integration in the brain. Feedback connections tend to terminate in distinct layers of cortex relative to the feedforward ones (Felleman and Van Essen, 1991; Callaway, 2004) making the idea of separate but interacting networks for recognition and generation potentially attractive^34^. Interestingly, such sub-networks might even be part of the same neuron and map onto “apical” versus “basal” parts of the dendritic tree (Körding and König, 2001; Urbanczik and Senn, 2014).

Generative models can also be trained via backpropagation. Recent advances have shown how to perform variational approximations to Bayesian inference inside backpropagation-based neural networks (Kingma and Welling, 2013), and how to exploit this to create generative models (Eslami et al., 2016; Gregor et al., 2015; Goodfellow et al., 2014a; Radford et al., 2015). Through either explicitly statistical or gradient descent based learning, the brain can thus obtain a probabilistic model that simulates features of the world.

#### 3.2.2 Cost functions that approximate properties of the world

A perceiving system should exploit statistical regularities in the world that are not present in an arbitrary dataset or input distribution. For example, objects are sparse, at least in certain representations: there are far fewer objects than there are potential places in the world, and of all possible objects there is only a small subset visible at any given time. As such, we know that the output of an object recognition system must have sparse activations. Building the assumption of sparseness into simulated systems replicates a number of representational properties of the early visual system (Olshausen and Field, 1997; Rozell et al., 2008), and indeed the original paper on sparse coding obtained sparsity by gradient descent optimization of a cost function (Olshausen and Field, 1996). A range of unsupervised machine learning techniques, such as the sparse autoencoders (Le et al., 2011) used to discover cats in YouTube videos, build sparseness into neural networks. Building in such spatio-temporal sparseness priors should serve as an “inductive bias” (Mitchell, 1980) that can accelerate learning.

But we know much more about the regularities of objects. As young babies, we already know (Bremner et al., 2015) that objects tend to persist over time. The emergence or disappearance of an object from a region of space is a rare event. Moreover, object locations and configurations tend to be coherent in time. We can formulate this prior knowledge as a cost function, for example by penalizing representations which are not temporally continuous. This idea of continuity is used in a great number of artificial neural networks and related models (Mobahi et al., 2009; Wiskott and Sejnowski, 2002; Földiák, 2008). Imposing continuity within certain models gives rise to aspects of the visual system including complex cells (Körding et al., 2004), specific properties of visual invariance (Isik et al., 2012), and even other representational properties such as the existence of place cells (Wyss et al., 2006; Franzius et al., 2007). Unsupervised learning mechanisms that maximize temporal coherence or slowness are increasingly used in machine learning^35^.

We also know that objects tend to undergo predictable sequences of transformations, and it is possible to build this assumption into unsupervised neural learning systems (George and Hawkins, 2009). The minimization of prediction error explains a number of properties of the nervous system (Friston and Stephan, 2007; Huang and Rao, 2011), and biologically plausible theories are available for how cortex could learn using prediction errors by exploiting temporal differences (O’Reilly et al., 2014b) or top-down feedback (George and Hawkins, 2009). In one implementation, a system can simply predict the next input delivered to the system and can then use the difference between the actual next input and the predicted next input as a full vectorial error signal for supervised gradient descent. Thus, rather than optimization of prediction error being implicitly implemented by the network dynamics, the prediction error is used as an explicit cost function in the manner of supervised learning, leading to error derivatives which can be back-propagated. Then, no special learning rules beyond simple backpropagation are needed. This approach has recently been advanced within machine learning (Lotter et al., 2015, 2016). Recently, combining such prediction-based learning with a specific gating mechanism has been shown to lead to unsupervised learning of disentangled representations (Whitney et al., 2016). Neural networks can also be designed to learn to invert spatial transformations (Jaderberg et al., 2015b). Statistically describing transformations or sequences is thus an unsupervised way of learning representations.

Furthermore, there are multiple modalities of input to the brain. Each sensory modality is primarily connected to one part of the brain^36^. But higher levels of cortex in each modality are heavily connected to the other modalities. This can enable forms of self-supervised learning: with a developing visual understanding of the world we can predict its sounds, and then test those predictions with the auditory input, and vice versa. The same is true about multiple parts of the same modality: if we understand the left half of the visual field, it tells us an awful lot about the right. Indeed, we can use observations of one part of a visual scene to predict the contents of other parts (van den Oord et al., 2016; Noroozi and Favaro, 2016), and optimize a cost function that reflects the discrepancy. Maximizing mutual information is a natural way of improving learning (Becker and Hinton, 1992; Mohamed and Rezende, 2015), and there are many other ways in which multiple modalities or processing streams could mutually train one another. This way, each modality effectively produces training signals for the others^37^. Evidence from psychophysics suggests that some kind of training via detection of sensory conflicts may be occurring in children (Nardini et al., 2010).

### 3.3 Cost functions for supervised learning

In what cases might the brain use supervised learning, given that it requires the system to “already know” the exact target pattern to train towards? One possibility is that the brain can store records of states that led to good outcomes. For example, if a baby reaches for a target and misses, and then tries again and successfully hits the target, then the difference in the neural representations of these two tries reflects the direction in which the system should change. The brain could potentially use a comparator circuit to directly compute this vectorial difference in the neural population codes and then apply this difference vector as an error signal.

Another possibility is that the brain uses supervised learning to implement a form of “chunking”, i.e., a consolidation of something the brain already knows how to do: routines that are initially learned as multi-step, deliberative procedures could be compiled down to more rapid and automatic functions by using supervised learning to train a network to mimic the overall input-output behavior of the original multi-step process. Such a process is assumed to occur in cognitive models like ACT-R (Servan-Schreiber and Anderson, 1990), and methods for compressing the knowledge in neural networks into smaller networks are also being developed (Ba and Caruana, 2014). Thus supervised learning can be used to train a network to do in “one step” what would otherwise require long-range routing and sequential recruitment of multiple systems.

### 3.4 Repurposing reinforcement learning for diverse internal cost functions

Certain generalized forms of reinforcement learning may be ubiquitous throughout the brain. Such reinforcement signals may be repurposed to optimize diverse internal cost functions. These internal cost functions could be specified at least in part by genetics.

Some brain systems such as in the striatum appear to learn via some form of temporal difference reinforcement learning (Tesauro, 1995; Foster et al., 2000). This is reinforcement learning based on a global value function (O’Reilly et al., 2014a) that predicts total future reward or utility for the agent. Reward-driven signaling is not restricted to the striatum, and is present even in primary visual cortex (Chubykin et al., 2013; Stnior et al., 2013). Remarkably, the reward signaling in primary visual cortex is mediated in part by glial cells (Takata et al., 2011), rather than neurons, and involves the neurotransmitter acetylcholine (Chubykin et al., 2013; Hangya et al., 2015). On the other hand, some studies have suggested that visual cortex learns the basics of invariant object recognition in the absence of reward (Li and Dicarlo, 2012), perhaps using reinforcement only for more refined perceptual learning (Roelfsema et al., 2010).

But beyond these well-known global reward signals, we argue that the basic mechanisms of reinforcement learning may be widely re-purposed to train local networks using a variety of internally generated error signals. These internally generated signals may allow a learning system to go beyond what can be learned via standard unsupervised methods, effectively guiding or steering the system to learn specific features or computations (Ullman et al., 2012).

#### 3.4.1 Cost functions for bootstrapping learning in the human environment

Special, internally-generated signals are needed specifically for learning problems where standard unsupervised methods – based purely on matching the statistics of the world, or on optimizing simple mathematical objectives like temporal continuity or sparsity – will fail to discover properties of the world which are statistically weak in an objective sense but nevertheless have special significance to the organism (Ullman et al., 2012). Indigo bunting birds, for example, learn a template for the constellations of the night sky long before ever leaving the nest to engage in navigation-dependent tasks (Emlen, 1967). This memory template is directly used to determine the direction of flight during migratory periods, a process that is modulated hormonally so that winter and summer flights are reversed. Learning is therefore a multi-phase process in which navigational cues are memorized prior to the acquisition of motor control.

In humans, we suspect that similar multistage bootstrapping processes are arranged to occur. Humans have innate specializations for social learning. We need to be able to read one another’s expressions as indicated with hands and faces. Hands are important because they allow us to learn about the set of actions that can be produced by agents (Ullman et al., 2012). Faces are important because they give us insight into what others are thinking. People have intentions and personalities that differ from one another, and their feelings are important. How could we hack together cost functions, built on simple genetically specifiable mechanisms, to make it easier for a learning system to discover such behaviorally relevant variables?

Some preliminary studies are beginning to suggest specific mechanisms and heuristics that humans may be using to bootstrap more sophisticated knowledge. In a groundbreaking study, Ullman et al. (2012) asked how could we explain hands, to a system that does not already know about them, in a cheap way, without the need for labeled training examples? Hands are common in our visual space and have special roles in the scene: they move objects, collect objects, and caress babies. Building these biases into an area specialized to detect hands could guide the right kind of learning, by providing a downstream learning system with many likely positive examples of hands on the basis of innately-stored, heuristic signatures about how hands tend to look or behave (Ullman et al., 2012). Indeed, an internally supervised learning algorithm containing specialized, hard-coded biases to detect hands, on the basis of their typical motion properties, can be used to bootstrap the training of an image recognition module that learns to recognize hands based on their appearance. Thus, a simple, hard-coded module bootstraps the training of a much more complex algorithm for visual recognition of hands.

Ullman et al. (2012) then further exploits a combination of hand and face detection to bootstrap a predictor for gaze direction, based on the heuristic that faces tend to be looking towards hands. Of course, given a hand detector, it also becomes much easier to train a system for reaching, crawling, and so forth. Efforts are underway in psychology to determine whether the heuristics discovered to be useful computationally are, in fact, being used by human children during learning (Fausey et al., 2016; Yu and Smith, 2013).

Ullman refers to such primitive, inbuilt detectors as innate “proto-concepts” (Ullman et al., 2012). Their broader claim is that such prespecification of mutual supervision signals can make learning the relevant features of the world far easier, by giving an otherwise unsupervised learner the right kinds of hints or heuristic biases at the right times. Here we call these approximate, heuristic cost functions “bootstrap cost functions”. The purpose of the bootstrap cost functions is to reduce the amount of data required to learn a specific feature or task, but at the same time to avoid a need for fully unsupervised learning.

Could the neural circuitry for such a bootstrap hand-detector be pre-specified genetically? The precedent from other organisms is strong: for example, it is famously known that the frog retina contains circuitry sufficient to implement a kind of “bug detector” (Lettvin et al., 1959). Ullman’s hand detector, in fact, operates via a simple local optical flow calculation to detect “mover” events. This type of simple, local calculation could potentially be implemented in genetically-specified and/or spontaneously self-organized neural circuitry in the retina or early dorsal visual areas (Biilthoff et al., 1989), perhaps similarly to the frog’s “bug detector”.

How could we explain faces without any training data? Faces tend to have two dark dots in their upper half, a line in the lower half and tend to be symmetric about a vertical axis. Indeed, we know that babies are very much attracted to things with these generic features of upright faces starting from birth, and that they will acquire face-specific cortical areas^38^ in their first few years of life if not earlier (McKone et al., 2009). It is easy to define a local rule that produces a kind of crude face detector (e.g., detecting two dots on top of a horizontal line), and indeed some evidence suggests that the brain can rapidly detect faces without even a single feed-forward pass through the ventral visual stream (Crouzet and Thorpe, 2011). The crude detection of human faces used together with statistical learning should be analogous to semi-supervised learning (Sukhbaatar et al., 2014) and could allow identifying faces with high certainty.

Humans have areas devoted to emotional processing, and the brain seems to embody prior knowledge about the structure of emotional expressions and how they relate to causes in the world: emotions should have specific types of strong couplings to various other higher-level variables such as goal-satisfaction, should be expressed through the face, and so on (Skerry and Spelke, 2014; Lyons and Cheries, 2016; Baillargeon et al., 2016; Phillips et al., 2002). What about agency? It makes sense to describe, when dealing with high-level thinking, other beings as optimizers of their own goal functions. It appears that heuristically specified notions of goals and agency are infused into human psychological development from early infancy and that notions of agency are used to bootstrap heuristics for ethical evaluation (Skerry and Spelke, 2014; Hamlin et al., 2007). Algorithms for establishing more complex, innately-important social relationships such as joint attention are under study (Gao et al., 2014), building upon more primitive proto-concepts like face detectors and Ullman’s hand detectors (Ullman et al., 2012). The brain can thus use innate detectors to create cost functions and training procedures to train the next stages of learning. This prior knowledge, encoded into brain structure via evolution, could allow learning signals to come from the right places and to appear developmentally at the right times.

It is intuitive to ask whether this type of bootstrapping poses a kind of “chicken and egg” problem: if the brain already has an inbuilt heuristic hand detector, how can it be used to train a detector that performs any better than those heuristics? After all, isn’t a trained system only as good as its training data? The work of Ullman et al. (2012) illustrates why this is not the case. First, the “innate detector” can be used to train a downstream detector that operates based on different cues: for example, based on the spatial and body context of the hand, rather than its motion. Second, once multiple such pathways of detection come into existence, they can be used to improve each other. In Ullman et al. (2012), appearance, body context, and mover motion are all used to bootstrap off of one another, creating a detector that is better than any of its training heuristics. In effect, the innate detectors are used not as supervision signals per se, but rather to guide or steer the learning process, enabling it to discover features that would otherwise be difficult. If such affordances can be found in other domains, it seems likely that the brain would make extensive use of them to ensure that developing animals learn the precise patterns of perception and behavior needed to ensure their later survival and reproduction.

Thus, generalizing previous ideas (Ullman et al., 2012; Poggio, 2015), we suggest that the brain uses optimization with respect to internally generated heuristic^39^ detection signals to bootstrap learning of biologically relevant features which would otherwise be missed by an unsupervised learner. In one possible implementation, such bootstrapping may occur via reinforcement learning, using the outputs of the innate detectors as local reinforcement signals, and perhaps using mechanisms similar to (Stnior et al., 2013; Rombouts et al., 2015; Brosch et al., 2015; Roelfsema and van Ooyen, 2005) to perform reinforcement learning through a multi-layer network. It is also possible that the brain could use such internally generated heuristic detectors in other ways, for example to bias the inputs delivered to an unsupervised learning network towards entities of interest to humans via an attentional process (Joscha Bach, personal communication), to bias hippocampal replay (Kumaran et al., 2016) or other aspects of memory access, or to directly train simple classifiers (Ullman et al., 2012).

#### 3.4.2 Cost functions for learning by imitation and through social feedback

It has been widely observed that the capacity for imitation and social learning may be a feature that is uniquely human, and that enables other human traits Ramachandran (2000). Humans need to learn more from the environment by than trial and error can provide for, and more than genetically orchestrated internal bootstrapping signals can effectively guide. Hence, babies spend a long time watching adults, especially adults they are attached to (Meltzoff, 1999), and later use specific kinds of social cues from their parents to shape their development. Babies and children learn about cause and effect through models based on goals, outcomes and agents, not just pure statistical inference. For example, young children make inferences about causality selectively in situations where a human is trying to achieve an outcome (Meltzoff et al., 2012, 2013). Minsky (2006) discusses how we derive not just skills but also goals from our attachment figures, through socially induced emotions like pride and shame. To do all this requires a powerful infrastructure of mental abilities: we must attribute social feedback to particular aspects of our goals or actions, and hence we need to signal to each other positively and negatively, to draw attention to these aspects. Minsky speculates (Minsky, 2006) that the development of such “learning by being told” led to language by selecting for the development of increasingly precise parsing of synatatic structures in relation to our representations of agents and action-plans.

How does this connect with cost functions? The idea of goals is central here, as we need to be able to identify the goals of others, update our own goals based on feedback, and measure the success of actions relative to goals. It has been proposed that human intrinsically use a model based on abstract goal and costs to underpin learning about the social world (Jara-Ettinger et al., 2016). Perhaps we even learn about our “selves” by inferring a model of our own goals and cost functions. Relatedly, machine learning in some settings can infer their cost functions from samples of behavior (Ho and Ermon, 2016).

#### 3.4.3 Cost functions for story generation and understanding

It has been widely noticed in cognitive science and AI that the generation and understanding of stories are crucial to human cognition. Researchers such as Winston have framed story understanding as the key to human-like intelligence (Winston, 2011). Stories consist of a linear sequence of episodes, in which one episode refers to another through cause and effect relationships, with these relationships often involving the implicit goals of agents. Many other cognitive faculties, such as conceptual grounding of language, could conceivably emerge from an underlying internal representation in terms of stories.

Perhaps the ultimate series of bootstrap cost functions would be those which would direct the brain to utilize its learning networks and specialized systems so as to construct representations that are specifically useful as components of stories, to spontaneously chain these representations together, and to update them through experience and communication. How could such cost functions arise? One possibility is that they are bootstrapped through imitation and communication, where a child learns to mimic the storytelling behavior of others. Another possibility is that useful representations and primitives for stories emerge spontaneously from mechanisms for learning state and action chunking in hierarchical reinforcement learning and planning. Yet another is that stories emerge from learned patterns of saliency-directed memory storage and recall (e.g., (Xiong et al., 2016)). In addition, priors that direct the developing child’s brain to learn about and attend to social agency seem to be important for stories.

In this section, we have seen how cost functions can be specified that could lead to the learning of increasingly sophisticated mental abilities in a biologically plausible manner. Importantly, however, cost functions and optimization are not the whole story. To achieve more complex forms of optimization, e.g., for learning to understand complex patterns of cause and effect over long timescales, to plan and reason prospectively, or to effectively coordinate many widely distributed brain resources, the brain seems to invoke specialized, pre-constructed data structures, algorithms and communication systems, which in turn facilitate specific kinds of optimization. Moreover, optimization occurs in a tightly orchestrated multi-stage process, and specialized, pre-structured brain systems need to be invoked to account for this meta-level of control over when, where and how each optimization problem is set up. We now turn to how these pre-specialized systems may orchestrate and facilitate optimization.

## 4. Optimization occurs in the context of specialized structures

Optimization of initially unstructured “blank slate” networks is not sufficient to generate complex cognition in the brain, we argue, even given a diversity of powerful genetically-specified cost functions and local learning rules, as we have posited above. Instead, in **Hypothesis 3**, we suggest that specialized, pre-structured architectures are needed for at least two purposes.

First, pre-structured architectures are needed to allow the brain to find efficient solutions to certain types of problems. When we write computer code, there are a broad range of algorithms and data structures employed for different purposes: we may use dynamic programming to solve planning problems, trees to efficiently implement nearest neighbor search, or stacks to implement recursion. Having the right kind of algorithm and data structure in place to solve a problem allows it to be solved efficiently, robustly and with a minimum amount of learning or optimization needed. This observation is concordant with the increasing use of pre-specialized architectures and specialized computational components in machine learning (Graves et al., 2014; Weston et al., 2014; Neelakantan et al., 2015). In particular, to enable the learning of efficient computational solutions, the brain may need pre-specialized systems for planning and executing sequential multi-step processes, for accessing memories, and for forming and manipulating compositional and recursive structures^40^.

Second, the training of optimization modules may need to be coordinated in a complex and dynamic fashion, including delivering the right training signals and activating the right learning rules in the right places and at the right times. To allow this, the brain may need specialized systems for storing and routing data, and for flexibly routing training signals such as target patterns, training data, reinforcement signals, attention signals, and modulatory signals. These mechanisms may need to be at least partially in place in advance of learning.

Looking at the brain, we indeed seem to find highly conserved structures, e.g., cortex, where it is theorized that a similar type of learning and/or computation is happening in multiple places (Douglas and Martin, 2004; Braitenberg and Schutz, 1991). But we also see a large number of specialized structures, including thalamus, hippocampus, basal ganglia and cerebellum (Solari and Stoner, 2011). These structures evolutionarily pre-date (Lee et al., 2015) the cortex, and hence the cortex may have evolved to work in the context of such specialized mechanisms. For example, the cortex may have evolved as a trainable module for which the training is orchestrated by these older structures.

Even within the cortex itself, microcircuitry within different areas may be specialized: tinkered variations on a common ancestral micro-circuit scaffold could potentially allow different cortical areas, such as sensory areas vs. pre-frontal areas, to be configured to adopt a number of qualitatively distinct computational and learning configurations (Marcus et al., 2014a, b; Yuste et al., 2005), even while sharing a common gross physical layout and communication interface. Within cortex, over forty distinct cell types – differing in such aspects as dendritic organization, distribution throughout the six cortical layers, connectivity pattern, gene expression, and electrophysiological properties – have already been found (Zeisel et al., 2015; Markram et al., 2015). Central pattern generator circuits provide an example of the kinds of architectures that can be pre-wired into neural microcircuitry, and may have evolutionary relationships with cortical circuits (Yuste et al., 2005). Thus, while the precise degree of architectural specificity of particular cortical regions is still under debate (Marcus et al., 2014a, b), various mechanisms could offer pre-specified heterogeneity.

In this section, we explore the kinds of computational problems for which specialized structures may be useful, and attempt to map these to putative elements within the brain. Our preliminary sketch of a functional decomposition can be viewed as a summary of suggestions for specialized functions that have been made throughout the computational neuroscience literature, and is influenced strongly by the models of O’Reilly, Eliasmith, Grossberg, Marcus, Hayworth and others (O’Reilly, 2006; Eliasmith et al., 2012; Marcus, 2001; Grossberg, 2013; Hayworth, 2012). The correspondence between these models and actual neural circuitry is, of course, still the subject of extensive debate.

Many of the computational and neural concepts sketched here are preliminary and will need to be made more rigorous through future study. Our knowledge of the functions of particular brain areas, and thus our proposed mappings of certain computations onto neuroanatomy, also remains tentative. Finally, it is still far from established which processes in the brain emerge from optimization of cost functions, which emerge from other forms of self-organization, which are pre-structured through genetics and development, and which rely on an interplay of all these mechanisms^41^. Our discussion here should therefore be viewed as a sketch of potential directions for further study.

### 4.1 Structured forms of memory

One of the central elements of computation is memory. Importantly, multiple different kinds of memory are needed (Squire, 2004). For example, we need memory that is stored for a long period of time and that can be retrieved in a number of ways, such as in situations similar to the time when the memory was first stored (content addressable memory). We also need memory that we can keep for a short period of time and that we can rapidly rewrite (working memory). Lastly, we need the kind of implicit memory that we cannot explicitly recall, similar to the kind of memory that is classically learned using gradient descent on errors, i.e., sculpted into the weight matrix of a neural network.

#### 4.1.1 Content addressable memories

Content addressable memories^42^ are classic models in neuroscience (Hopfield, 1982). Most simply, they allow us to recognize a situation similar to one that we have seen before, and to “fill in” stored patterns based on partial or noisy information, but they may also be put to use as sub-components of many other functions. Recent research has shown that including such memories allows deep networks to learn to solve problems that previously were out of reach, even of LSTM networks that already have a simpler form of local memory and are already capable of learning long-term dependencies (Weston et al., 2014; Graves et al., 2014). Hippocampal area CA3 may act as an auto-associative memory^43^ capable of content-addressable pattern completion, with pattern separation occurring in the dentate gyrus (Rolls, 2013). If no similar pattern is available, an unfamiliar input will be stored as a new memory (Kumaran et al., 2016). Such systems could permit the retrieval of complete memories from partial cues, enabling networks to perform operations similar to database retrieval or to instantiate lookup tables of historical stimulus-response mappings, among numerous other possibilities.

Of course, memory systems may be organized – through cost function optimization or other mechanisms – into higher-order structures. Cost functions might be used to bias memory representations to adopt particular structures, e.g., to be organized into data structures like like Minskys frames and trans-frames (Minsky, 2006).

#### 4.1.2 Working memory buffers

Cognitive science has long characterized properties of the working memory. Its capacity is somewhat limited, with the old idea being that verbal working memory has a capacity of “seven plus or minus two” (Miller, 1956), while visual working memory has a capacity of four (Luck and Vogel, 1997) (or, other authors defend, one). There are many models of working memory (Wang, 2012; Singh and Eliasmith, 2006; O’Reilly and Frank, 2006; Buschman and Miller, 2014; Warden and Miller, 2007), some of which attribute it to persistent, self-reinforcing patterns of neural activation (Goldman et al., 2003) in the recurrent networks of the prefrontal cortex. Prefrontal working memory appears to be made up of multiple functionally distinct subsystems (Markowitz et al., 2015). Neural models of working memory can store not only scalar variables (Seung, 1998), but also high-dimensional vectors (Eliasmith and Anderson, 2004; Eliasmith et al., 2012) or sequences of vectors (Choo and Eliasmith, 2010). Working memory buffers seem crucial for human-like cognition, e.g., reasoning, as they allow short-term storage while also – in conjunction with other mechanisms – enabling generalization of operations across anything that can fill the buffer.

#### 4.1.3 Storing state in association with saliency

Saliency, or interestingness, measures can be used to tag the importance of a memory (Gonzalez Andino and Grave de Peralta Menendez, 2012). This can allow removal of the boring data from the training set, allowing a mechanism that is more like optimal experimentation. Moreover, saliency can guide memory replay or sampling from generative models, to generate more training data drawn from a distribution useful for learning (Mnih et al., 2015; Ji and Wilson, 2007). Conceivably, hippocampal replay could allow a batch-like training process, similar to how most machine learning systems are trained, rather than requiring all training to occur in an online fashion. Plasticity mechanisms in memory systems which are gated by saliency are starting to be uncovered in neuroscience (Dudman et al., 2007). Importantly, the notions of “saliency” computed by the brain could be quite intricate and multi-faceted, potentially leading to complex schemes by which specific kinds of memories would be tagged for later context-dependent retrieval. As a hypothetical example, representations of both timing and importance associated with memories could perhaps allow retrieval only of important memories that happened within a certain window of time (MacDonald et al., 2011; Kraus et al., 2013; Rubin et al., 2015). Storing and retrieving information selectively based on specific properties of the information itself, or of “tags” appended to that information, is a powerful computational primitive that could enable learning of more complex tasks. Relatedly, we know that certain pathways become associated with certain kinds of memories, e.g., specific pathways for fear-related memory in mice.

### 4.2 Structured routing systems

To use its information flexibly, the brain needs structured systems for routing data. Such systems need to address multiple temporal and spatial scales, and multiple modalities of control. Thus, there are several different kinds of information routing systems in the brain which operate by different mechanisms and under different constraints.

#### 4.2.1 Attention

If we can focus on one thing at a time, we may be able to allocate more computational resources to processing it, make better use of scarce data to learn about it, and more easily store and retrieve it from memory^44^. Notably in this context, attention allows improvements in learning: if we can focus on just a single object, instead of an entire scene, we can learn about it more easily using limited data. Formal accounts in a Bayesian framework talk about attention reducing the sample complexity of learning (Chikkerur et al., 2010). Likewise, in models, the processes of applying attention, and of effectively making use of incoming attentional signals to appropriately modulate local circuit activity, can themselves be learned by optimizing cost functions (Mnih et al., 2014; Jaramillo and Pearlmutter, 2004). The right kinds of attention make processing and learning more efficient, and also allow for a kind of programmatic control over multi-step perceptual tasks.

How does the brain determine where to allocate attention, and how is the attentional signal physically mediated? Answering this question is still an active area of neuroscience. Higher-level cortical areas may be specialized in allocating attention. The problem is made complex by the fact that there seem to be many different types of attention – such as object-based, feature-based and spatial attention in vision – that may be mediated by interactions between different brain areas. The frontal eye fields (area FEF), for example, are important in visual attention, specifically for controlling saccades of the eyes to attended locations. Area FEF contains “retinotopic” spatial maps whose activation determines the saccade targets in the visual field. Other prefrontal areas such as the dorsolateral prefrontal cortex and inferior frontal junction are also involved in maintaining representations that specify the targets of certain types of attention. Certain forms of attention may require a complex interaction between brain areas, e.g., to determine targets of attention based on higher-level properties that are represented across multiple areas, like the identity and spatial location of a specific face (Baldauf and Desimone, 2014).

There are many proposed neural mechanisms of attention, including the idea that synchrony plays a role (Baldauf and Desimone, 2014), perhaps by creating resonances that facilitate the transfer of information between synchronously oscillating neural populations in different areas^45^. Other proposed mechanisms include specific circuits for attention-dependent signal routing (Anderson and Van Essen, 1987; Olshausen et al., 1993). Various forms of attention also have specific neurophysiological signatures, such as enhancements in synchrony among neural spikes and with the ambient local field potential, changes in the sharpness of neural tuning curves, and other properties. These diverse effects and signatures of attention may be consequences of underlying pathways that wire up to particular elements of cortical microcircuits to mediate different attentional effects (Bobier et al., 2014).

#### 4.2.2 Buffers

One possibility is that the brain uses distinct groups of neurons, which we can call “buffers”, to store distinct variables, such as the subject or object in a sentence (Frankland and Greene, 2015). Having memory buffers allows the abstraction of a variable.

Once we establish that the brain has a number of memory buffers, we need ways for those buffers to interact. We need to be able to take a buffer, do a computation on its contents and store the output into another buffer. But if the representations in each of two groups of neurons are learned, and hence are coded differently, how can the brain “copy and paste” information between these groups of neurons? Malsburg argued that such a system of separate buffers is impossible because the neural pattern for “chair” in buffer 1 has nothing in common with the neural pattern for “chair” in buffer 2 – any learning that occurs for the contents of buffer 1 would not automatically be transferable to buffer 2. Various mechanisms have been proposed to allow such transferability, which focus on ways in which all buffers could be trained jointly and then later separated so that they can work independently when they need to^46^.

#### 4.2.3 Discrete gating of information flow between buffers

Dense connectivity is only achieved locally, but it would be desirable to have a way for any two cortical units to talk to one another, if needed, regardless of their distance from one another, and without introducing crosstalk^47^. It is therefore critical to be able to dynamically turn on and off the transfer of information between different source and destination regions, in much the manner of a switchboard. Together with attention, such dedicated routing systems can make sure that a brain area receives exactly the information it needs. Such a discrete routing system is, of course, central to cognitive architectures like ACT-R (Anderson, 2007). The key feature of ACT-R is the ability to evaluate the IF clauses of tens of thousands of symbolic rules (called “productions”), in parallel, approximately every 50 milliseconds. Each rule requires equality comparisons between the contents of many constant and variable memory buffers, and the execution of a rule leads to the conditional routing of information from one buffer to another.

What controls which long-range routing operations occur when, i.e., where is the switchboad and what controls it? Several models, including ACT-R, have attributed such parallel rule-based control of routing to the action selection circuitry (Gurney et al., 2001; Terrence Stewart, 2010) of the basal ganglia (BG) (Stocco et al., 2010; O’Reilly and Frank, 2006), and its interaction with working memory buffers in the prefrontal cortex. In conventional models of thalamo-cortico-striatal loops, competing actions of the direct and indirect pathways through the basal ganglia can inhibit or disinhibit an area of motor cortex, thereby gating a motor action^48^. Models like (Stocco et al., 2010; O’Reilly and Frank, 2006; Terrence Stewart, 2010) propose further that the basal ganglia can gate not just the transfer of information from motor cortex to downstream actuators, but also the transfer of information between cortical areas. To do so, the basal ganglia would dis-inhibit a thalamic relay (Sherman, 2005, 2007) linking two cortical areas. Dopamine-related activity is thought to lead to temporal difference reinforcement learning of such gating policies in the basal ganglia (Frank and Badre, 2012). Beyond the basal ganglia, there are also other, separate pathways involved in action selection, e.g., in the prefrontal cortex (Daw et al., 2006). Thus, multiple systems including basal ganglia and cortex could control the gating of long-range information transfer between cortical areas, with the thalamus perhaps largely constituting the switchboard itself.

How is such routing put to use in a learning context? One possibility is that the basal ganglia acts to orchestrate the training of the cortex. The basal ganglia may exert tight control^49^ over the cortex, helping to determine when and how it is trained. Indeed, because the basal ganglia pre-dates the cortex evolutionarily, it is possible that the cortex evolved as a flexible, trainable resource that could be harnessed by existing basal ganglia circuitry. All of the main regions and circuits of the basal ganglia are conserved from our common ancestor with the lamprey more than five hundred million years ago. The major part of the basal ganglia even seems to be conserved from our common ancestor with insects (Strausfeld and Hirth, 2013). Thus, in addition to its real-time action selection and routing functions, the basal ganglia may sculpt how the cortex learns.

### 4.3 Structured state representations to enable efficient algorithms

Certain algorithmic problems benefit greatly from particular types of representation and transformation, such as a grid-like representation of space. In some cases, rather than just waiting for them to emerge via gradient descent optimization of appropriate cost functions, the brain may be pre-structured to facilitate their creation.

#### 4.3.1 Continuous predictive control

We often have to plan and execute complicated sequences of actions on the fly, in response to a new situation. At the lowest level, that of motor control, our body and our immediate environment change all the time. As such, it is important for us to maintain knowledge about this environment in a continuous way. The deviations between our planned movements and those movements that we actually execute continuously provide information about the properties of the environment. Therefore it seems important to have a specialized system, optimized for high-speed continuous processing, that takes all our motor errors and uses them to update a dynamical model of our body and our immediate environment that can predict the delayed sensory results of our motor actions (McKinstry et al., 2006).

It appears that the cerebellum is such a structure, and lesions to it abolish our way of dealing successfully with a changing body. Incidentally, the cerebellum has more connections than the rest of the brain taken together, apparently in a largely feedforward architecture, and the tiny cerebellar granule cells, which may form a randomized high-dimensional input representation (Marr, 1969; Jacobson and Friedrich, 2013), outnumber all other neurons. The brain clearly needs a dedicated way of quickly and continuously correcting movements to minimize errors, without needing to rely on slow and complex association learning in the neocortex in order to do so.

Newer research shows that the cerebellum is involved in a broad range of cognitive problems (Moberget et al., 2014) as well, potentially because they share computational problems with motor control. For example, when subjects estimate time intervals, which are naturally important for movement, it appears that the brain uses the cerebellum even if no movements are involved (Gooch et al., 2010). Even individual cerebellar Purkinjie cells may learn to generate precise timings of their outputs (Johansson et al., 2014). The brain also appears to use inverse models to rapidly predict motor activity that would give rise to a given sensory target (Giret et al., 2014; Hanuschkin et al., 2013). Such mechanisms could be put to use far beyond motor control, in bootstrapping the training of a larger architecture by exploiting continuously changing error signals to update a real-time model of the system state.

#### 4.3.2 Hierarchical control

Importantly, many of the control problems we appear to be solving are hierarchical. We have a spinal cord, which deals with the fast signals coming from our muscles and proprioception. Within neuroscience, it is generally assumed that this system deals with fast feedback loops and that this behavior is learned to optimize its own cost function. The nature of cost functions in motor control is still under debate. In particular, the timescale over which cost functions operate remains unclear: motor optimization may occur via real-time responses to a cost function that is computed and optimized online, or via policy choices that change over time more slowly in response to the cost function (Körding, 2007). Nevertheless, the effect is that central processing in the brain has an effectively simplified physical system to control, e.g., one that is far more linear. So the spinal cord itself already suggests the existence of two levels of a hierarchy, each trained using different cost functions.

However, within the computational motor control literature (see e.g., (DeWolf and Eliasmith, 2011)), this idea can be pushed far further, e.g., with a hierarchy including spinal cord, M1, PMd, frontal, prefrontal areas. A low level may deal with muscles, the next level may deal with getting our limbs to places or moving objects, a next layer may deal with solving simple local problems (e.g., navigating across a room) while the highest levels may deal with us planning our path through life. This factorization of the problem comes with multiple aspects: First, each level can be solved with its own cost functions, and second, every layer has a characteristic timescale. Some levels, e.g., the spinal cord, must run at a high speed. Other levels, e.g., high-level planning, only need to be touched much more rarely. Converting the computationally hard optimal control problem into a hierarchical approximation promises to make it dramatically easier.

Does the brain solve control problems hierarchically? There is evidence that the brain uses such a strategy (Botvinick and Weinstein, 2014; Botvinick et al., 2009), beside neural network demonstrations (Wayne and Abbott, 2014). The brain may use specialized structures at each hierarchical level to ensure that each operates efficiently given the nature of its problem space and available training signals. At higher levels, these systems may use an abstract syntax for combining sequences of actions in pursuit of goals (Allen et al., 2010). Subroutines in such processes could be derived by a process of chunking sequences of actions into single actions (Botvinick and Weinstein, 2014; Graybiel, 1998). Some brain areas like Broca’s area, known for its involvement in language, also appear to be specifically involved in processing the hierarchical structure of behavior, as such, as opposed to its detailed temporal structure (Koechlin and Jubault, 2006).

At the highest level of the decision making and control hierarchy, human reward systems reflect changing goals and subgoals, and we are only beginning to understand how goals are actually coded in the brain, how we switch between goals, and how the cost functions used in learning depend on goal state (O’Reilly et al., 2014b; Buschman and Miller, 2014; Pezzulo et al., 2014). Goal hierarchies are beginning to be incorporated into deep learning (Kulkarni et al., 2016).

Given this hierarchical structure, the optimization algorithms can be fine-tuned. For the low levels, there is sheer unlimited training data. For the high levels, a simulation of the world may be simple, with a tractable number of high-level actions to choose from. Finally, each area needs to give reinforcement to other areas, e.g., high levels need to punish lower levels for making planning complicated. Thus this type of architecture can simplify the learning of control problems.

Progress is being made in both neuroscience and machine learning on finding potential mechanisms for this type of hierarchical planning and goal-seeking. This is beginning to reveal mechanisms for chunking goals and actions and for searching and pruning decision trees (Balaguer et al., 2016; Krishnamurthy et al., 2016; Tamar et al., 2016; O’Reilly et al., 2014a; Huys et al., 2015). The study of model-based hierarchical reinforcement learning and prospective optimization (Sejnowski and Poizner, 2014), which concerns the planning and evaluation of nested sequences of actions, implicates a network coupling the dorsolateral prefontral and orbitofrontal cortex, and the ventral and dorsolateral striatum (Botvinick et al., 2009). Hierarchical RL relies on a hierarchical representation of state and action spaces, and it has been suggested that error-driven learning of an optimal such representation in the hippocampus^50^ gives rise to place and grid cell properties (Stachenfeld, 2014), with goal representations themselves emerging in the amygdala, prefrontal cortex and other areas (O’Reilly et al., 2014a).

The question of how control problems can be successfully divided into component problems remains one of the central questions in neuroscience (Wolpert and Flanagan, 2016) and machine learning (Kulkarni et al., 2016), and the cost functions involved in learning to create such decompositions are still unknown. These considerations may begin to make plausible, however, how the brain could not only achieve its remarkable feats of motor learning – such as generating complex “innate” motor programs, like walking in the newborn gazelle almost immediately after birth – but also the kind of planning that allows a human to prepare a meal or travel from London to Chicago.

#### 4.3.3 Spatial planning

Spatial planning requires solving shortest-path problems subject to constraints. If we want to get from one location to another, there are an arbitrarily large number of simple paths that could be taken. Most naive implementations of such shortest paths problems are grossly inefficient. It appears that, in animals, the hippocampus aids – at least in part through “place cell” and “grid cell” systems – in efficient learning about new environments and in targeted navigation in such environments (Brown et al., 2016). Interestingly, once an environment becomes familiar, it appears that areas of the neocortex can take over the role of navigation (Hasselmo and Stern, 2015).

In some simple models, targeted navigation in the hippocampus is achieved via the dynamics of “bump attractors” or propagating waves in a place cell network with Hebbian plasticity and adaptation (Buzsáki and Moser, 2013; Hopfield, 2009; Ponulak and Hopfield, 2013), which allows the network to effectively chart out a path in the space of place cell representations. Other navigation models make use of the grid cell system. The place cell network may^51^ take input from a grid cell network that computes precise distances and directions, perhaps by integrating head direction and velocity signals – grid cells fire when the animal is on any node of a regularly spaced hexagonal grid. Different parts of the entorhinal cortex contain grid cells with different grid spacings, and place cells may combine information from multiple such grids in order to build up responses to particular single positions. These systems are highly structured temporally, e.g., containing nested gamma and theta oscillation structures that are phased locked to sequences of place-cell responses, interfering oscillators frequency-shifted by the animal’s motion velocity (Zilli and Hasselmo, 2010), tuned cellular resonances (Buzsáki, 2010; Giocomo et al., 2007), and other neural phenomena that lie far outside a conventional artificial neural network description. It seems that an intricate interplay of spatial and temporal network structures may be essential for encoding sequences of spatiotemporal events across multiple scales, and using them to drive multiple forms of learning, e.g., supporting forward and reverse sequence replay with various temporal compression factors (Buzsáki, 2010).

Higher-level cognitive tasks such as prospective planning appear to share computational sub-problems with path-finding (Hassabis and Maguire, 2009)^52^. Interaction between hippocampus and prefrontal cortex could perhaps support a more abstract notion of “navigation” in a space of goals and sub-goals. Interestingly, there is preliminary evidence from fMRI that abstract concepts are also represented according to grid-cell-like hexagonal grid structures in humans (Constantinescu et al., 2016), as well as preliminary evidence that social relationships may also be represented through a hippocampal map (Tavares et al., 2015). Having specialized structures for path-finding could thus simplify a variety of computational problems at different levels of abstraction.

#### 4.3.4 Variable binding

Language and reasoning appear to present a problem for neural networks (Minsky, 1991; Hadley, 2009; Marcus, 2001): we seem to be able to apply common grammatical rules to sentences regardless of the content of those sentences, and regardless of whether we have ever seen even remotely similar sentences in the training data. While this is achieved automatically in a computer with fixed registers, location addressable memories, and hard-coded operations, how it could be achieved in a biological brain, or emerge from an optimization algorithm, has been under debate for decades.

As the putative key capability underlying such operations, variable binding has been defined as “the transitory or permanent tying together of two bits of information: a variable (such as an X or Y in algebra, or a placeholder like subject or verb in a sentence) and an arbitrary instantiation of that variable (say, a single number, symbol, vector, or word)” (Marcus et al., 2014a, b). A number of potential biologically plausible binding mechanisms (Hayworth, 2012; Kriete et al., 2013; Eliasmith et al., 2012; Goertzel, 2014) are reviewed in (Marcus et al., 2014a, b). Some, such as vector symbolic architectures^53^, which were proposed in cognitive science (Eliasmith, 2013; Plate, 1995; Stewart and Eliasmith, 2009), are also being considered in the context of efficiently-trainable artificial neural networks (Danihelka et al., 2016) – in effect, these systems learn how to use variable binding.

Variable binding could potentially emerge from simpler memory systems. For example, the Scrub-Jay can remember the place and time of last visit for hundreds of different locations, e.g., to determine whether high-quality food is currently buried at any given location (Clayton and Dickinson, 1998). It is conceivable that such spatially-grounded memory systems enabled a more general binding mechanism to emerge during evolution, perhaps through integration with routing systems or other content-addressable or working memory systems.

#### 4.3.5 Hierarchical syntax

Fixed, static hierarchies (e.g., the hierarchical organization of cortical areas (Felleman and Van Essen, 1991)) only take us so far: to deal with long chains of arbitrary nested references, we need *dynamic* hierarchies that can implement recursion on the fly. Human language syntax has a hierarchical structure, which Berwick et al described as “composition of smaller forms like words and phrases into larger ones” (Berwick et al., 2012; Miyagawa et al., 2013). The extent of recursion in human language and thought may be captured by a class of automata known as higher-order pushdown automata, which can be implemented via finite state machines with access to nested stacks (Rodriguez and Granger, 2016). Specific fronto-temporal networks may be involved in representing and generating such hierarchies (Dehaene et al., 2015), e.g., with the hippocampal system playing a key role in implementing some analog of a pushdown stack (Rodriguez and Granger, 2016)^54^.

Little is known about the underlying circuit mechanisms for such dynamic hierarchies, but it is clear that specific affordances for representing such hierarchies in an efficient way would be beneficial. This may be closely connected with the issue of variable binding, and it is possible that operations similar to pointers could be useful in this context, in both the brain and artificial neural networks (Kriete et al., 2013; Kurach et al., 2015). Augmenting neural networks with a differentiable analog of a push-down stack is another such affordance being pursued in machine learning (Joulin and Mikolov, 2015).

#### 4.3.6 Mental programs and imagination

Humans excel at stitching together sub-actions to form larger actions (Sejnowski and Poizner, 2014; Acuna et al., 2014; Verwey, 1996). Structured, serial, hierarchical probabilistic programs have recently been shown to model aspects of human conceptual representation and compositional learning (Lake et al., 2015). In particular, sequential programs were found to enable one-shot learning of new geometric/visual concepts (Lake et al., 2015). Generative programs have also been proposed in the context of scene understanding (Battaglia et al., 2013). The ability to deal with problems in terms of subproblems is central both in human thought and in many successful algorithms.

One possibility is that the hippocampus supports the rapid construction and learning of sequential programs, e.g., in multi-step planning. An influential idea – known as the “complementary learning systems hypothesis” – is that the hippocampus plays a key role in certain processes where learning must occur quickly on the basis of single episodes, whereas the cortex learns more slowly by aggregating and integrating patterns across large amounts of data (Leibo et al., 2015a; Herd et al., 2013; Blundell et al., 2016; Kumaran et al., 2016). The hippocampus appears to explore, in simulation, possible future trajectories to a goal, even those involving previously unvisited locations (Ólafsdóttir et al., 2015). Hippocampal-prefrontal interaction has been suggested to allow rapid, subconscious evaluation of potential action sequences during decision-making, with the hippocampus in effect simulating the expected outcomes of potential actions that are generated and evaluated in the prefrontal (Wang et al., 2015; Mushiake et al., 2006). The role of the hippocampus in imagination, concept generation (Kumaran et al., 2009), scene construction (Hassabis and Maguire, 2007), mental exploration and goal-directed path planning (Ólafsdóttir et al., 2015; Brown et al., 2016; Hopfield, 2009) suggests that it could help to create generative models to underpin more complex inference such as program induction (Lake et al., 2015) or common-sense world simulation (Battaglia et al., 2013). For example, a sequential, programmatic process, mediated jointly by the basal ganglia, hippocampus and prefrontal cortex might allow one-shot learning of a new concept, as in the sequential computations underlying a process like Bayesian Program Learning (Lake et al., 2015).

Another related possibility is that the cortex itself intrinsically supports the construction and learning of sequential programs (Bach and Herger, 2015). Recurrent neural networks have been used for image generation through a sequential, attention-based process (Gregor et al., 2015), although their correspondence with the brain is unclear^55^.

### 4.4 Other specialized structures

Importantly, there are many other specialized structures known in neuroscience, which arguably receive less attention than they deserve, even for those interested in higher cognition. In the above, in addition to the hippocampus, basal ganglia and cortex, we emphasized the key roles of the thalamus in routing, of the cerebellum as a fast and rapidly trainable control and modeling system, of the amygdala and other areas as a potential source of utility functions, of the retina or early visual areas as a means to generate detectors for motion and other features to bootstrap more complex visual learning, and of the frontal eye fields and other areas as a possible source of attention control. We ignored other structures entirely, whose functions are only beginning to be uncovered, such as the claustrum (Crick and Koch, 2005), which has been speculated to be important for rapidly binding together information from many modalities. Our overall understanding of the functional decomposition of brain circuitry still seems very preliminary.

### 4.5 Relationships with other cognitive frameworks involving specialized systems

A recent analysis (Lake et al., 2016) suggested directions by which to modify and enhance existing neural-net-based machine learning towards more powerful and human-like cognitive capabilities, particularly by introducing new structures and systems which go beyond data-driven optimization. This analysis emphasized that systems should construct generative models of the world that incorporate compositionality (discrete construction from re-usable parts), inductive biases reflecting causality, intuitive physics and intuitive psychology, and the capacity for probabilistic inference over discrete structured models (e.g., structured as graphs, trees, or programs) (Tervo et al., 2016) to harness abstractions and enable transfer learning.

We view these ideas as consistent with and complementary to the framework of cost functions, optimization and specialized systems discussed here. One might seek to understand how optimization and specialized systems could be used to implement some of the mechanisms proposed in (Lake et al., 2016) inside neural networks. Lake et al. (2016) emphasize how incorporating additional structure into trainable neural networks can potentially give rise to systems that use compositional, causal and intuitive inductive biases and that “learn to learn” using structured models and shared data structures. For example, sub-dividing networks into units that can be modularly and dynamically combined, where representations can be copied and routed, may present a path towards improved compositionality and transfer learning (Andreas et al., 2015). The control flow for recombining pre-existing modules and representations could be learned via reinforcement learning (Andreas et al., 2016). How to implement the broad set of mechanisms discussed in (Lake et al., 2016) is a key computational problem, and it remains open at which levels (e.g., cost functions and training procedures vs. specialized computational structures vs. underlying neural primitives) architectural innovations will need to be introduced to capture these phenomena.

Primitives that are more complex than those used in conventional neural networks – for instance, primitives that act as state machines with complex message passing (Bach and Herger, 2015) or networks that intrinsically implement Bayesian inference (George and Hawkins, 2009) – could potentially be useful, and it is plausible that some of these may be found in the brain. Recent findings on the power of generic optimization also do not rule out the idea that the brain may explicitly generate and use particular types of structured representations to constrain its inferences; indeed, the specialized brain systems discussed here might provide a means to enforce such constraints. It might be possible to further map the concepts of Lake et al. (2016) onto neuroscience via an infrastructure of interacting cost functions and specialized brain systems under rich genetic control, coupled to a powerful and generic neurally implemented capacity for optimization. For example, it was recently shown that complex probabilistic population coding and inference can arise automatically from backpropagation-based training of simple neural networks (Orhan and Ma, 2016), without needing to be built in by hand. The nature of the underlying primitives in the brain, on top of which learning can operate, is a key question for neuroscience.

## 5. Machine learning inspired neuroscience

Hypotheses are primarily useful if they lead to concrete, experimentally testable predictions. As such, we now want to go through the hypotheses and see to which level they can be directly tested, as well as refined, through neuroscience.

### 5.1 *Hypothesis 1*– Existence of cost functions

There are multiple general strategies for addressing whether and how the brain optimizes cost functions. A first strategy is based on observing the endpoint of learning. If the brain uses a cost function, and we can guess its identity, then the final state of the brain should be close to optimal for the cost function. We could thus compare (Güçlü and van Gerven, 2015) receptive fields that are optimized in a simulation, according to a particular cost function, with the measured receptive fields. Various techniques exist to carry out such comparisons in fRMI studies, including population receptive field estimation (Güçlü and van Gerven, 2015; Dumoulin and Wandell, 2008) and representational dissimilarity matrices (Kriegeskorte et al., 2008; Khaligh-Razavi and Kriegeskorte, 2014). This strategy is only beginning to be used at the moment, perhaps because it has been difficult to measure the receptive fields or other representational properties across a large population of *individual* neurons (fMRI operates at a much coarser level), but this situation is beginning to improve technologically with the emergence of large-scale recording methods (Hasselmo, 2015).

A second strategy could directly quantify how well a cost function describes learning. If the dynamics of learning minimize a cost function then the underlying vector field should have a strong gradient descent type component and a weak rotational component, i.e., weight changes will primarily move down the gradient rather than drifting in the nullspace. If we could somehow continuously monitor the synaptic strengths, while externally manipulating them, then we could, in principle, measure the vector field in the space of synaptic weights, and calculate its divergence as well as its rotation. For at least the subset of synapses that are being trained via some approximation to gradient descent, the divergence component should be strong relative to the rotational component. This strategy has not been developed yet due to experimental difficulties with monitoring large numbers of synaptic weights^56^.

A third strategy is based on perturbations: cost function based learning should undo the effects of perturbations which disrupt optimality, i.e., the system should return to local minima after a perturbation, and indeed perhaps to the same local minimum after a sufficiently small perturbation. If we change synaptic connections, e.g., in the context of a brain machine interface, we should be able to produce a reorganization that can be predicted based on a guess of the relevant cost function. This strategy is starting to be feasible in motor areas.

Lastly, if we knew structurally which cell types and connections mediated the delivery of error signals vs. input data or other types of connections, then we could stimulate specific connections so as to impose a user-defined cost function. In effect, we would use the brain’s own networks as a trainable deep learning substrate, and then study how the network responds to training. Brain machine interfaces can be used to set up specific local learning problems, in which the brain is asked to create certain user-specified representations, and the dynamics of this process can be monitored (Sadtler et al., 2014). Likewise, brain machine interfaces can be used to give the brain access to new datastreams, and to investigate how those datastreams are incorporated into task performance, and whether such incorporation is governed by optimality principles (Dadarlat et al., 2015). In order to do this kind of experiment fully and optimally, we must first understand more about how the system is wired to deliver cost signals. Much of the structure that would be found in connectomic circuit maps, for example, would not just be relevant for short-timescale computing, but also for creating the infrastructure that supports cost functions and their optimization.

Many of the learning mechanisms that we have discussed in this paper make specific predictions about connectivity or dynamics. For example, the “feedback alignment” approach to biological backpropagation suggests that cortical feedback connections should, at some level of neuronal grouping, be largely sign-concordant with the corresponding feedforward connections, although not necessarily of concordant weight (Liao et al., 2015), and feedback alignment also makes predictions for synaptic normalization mechanisms (Liao et al., 2015). The Kickback model for biologically plausible back-propagation has a specific role for NMDA receptors (Balduzzi et al., 2014). Some models that incorporate dendritic coincidence detection for learning temporal sequences predict that a given axon should make only a small number of synapses on a given dendritic segment (Hawkins and Ahmad, 2016). Models that involve STDP learning will make predictions about the dynamics of changing firing rates (Hinton, 2007, 2016; Bengio et al., 2015a; Bengio and Fischer, 2015; Bengio et al., 2015b), as well as about the particular network structures, such as those based on autoencoders or recirculation, in which STDP can give rise to a form of backpropagation.

It is critical to establish the unit of optimization. We want to know the scale of the modules that are trainable by some approximation of gradient descent optimization. How large are the networks which share a given error signal or cost function? On what scales can appropriate training signals be delivered? It could be that the whole brain is optimized end-to-end, in principle. In this case we would expect to find connections that carry training signals from each layer to the preceding ones. On successively smaller scales, optimization could be within a brain area, a microcircuit^57^, or an individual neuron (Körding and König, 2000, 2001; Mel, 1992; Hawkins and Ahmad, 2016). Importantly, optimization may co-exist across these scales. There may be some slow optimization end-to-end, with stronger optimization within a local area and very efficient algorithms within each cell. Careful experiments should be able to identify the scale of optimization, e.g., by quantifying the extent of learning induced by a local perturbation.

The tightness of the structure-function relationship is the hallmark of molecular and to some extent cellular biology, but in large connectionist learning systems, this relationship can become difficult to extract: the same initial network can be driven to compute many different functions by subjecting it to different training^58^ ^59^. It can be hard to understand the way a neural network solves its problems.

How could one tell the difference, then, between a gradient-descent trained network vs. untrained or random networks vs. a network that has been trained against a different kind of task? One possibility would be to train artificial neural networks against various candidate cost functions, study the resulting neural tuning properties (Todorov, 2002), and compare them with those found in the circuit of interest (Zipser and Andersen, 1988). This has already been done to aid the interpretation of the neural dynamics underlying decision making in the PFC (Sussillo, 2014), working memory in the posterior parietal cortex (Rajan et al., 2016) and object or action representation in the visual system (Yamins and DiCarlo, 2016b, a; Tacchetti et al., 2016). Some have gone on to suggest a direct correspondence between cortical circuits and optimized, appropriately regularized (Sussillo et al., 2015), recurrent neural networks (Liao and Poggio, 2016). In any case, effective analytical methods to reverse engineer complex machine learning systems (Jonas and Kording, 2016), and methods to reverse engineer biological brains, may have some commonalities.

Does this emphasis on function optimization and trainable substrates mean that we should give up on reverse engineering the brain based on detailed measurements and models of its specific connectivity and dynamics? On the contrary: we should use large-scale brain maps to try to better understand a) how the brain implements optimization, b) where the training signals come from and what cost functions they embody, and c) what structures exist, at different levels of organization, to constrain this optimization to efficiently find solutions to specific kinds of problems. The answers may be influenced by diverse local properties of neurons and networks, such as homeostatic rules of neural structure, gene expression and function (Marder and Goaillard, 2006), the diversity of synapse types, cell-type-specific connectivity (Jiang et al., 2015), patterns of inter-laminar projection, distributions of inhibitory neuron types, dendritic targeting and local dendritic physiology and plasticity (Markram et al., 2015; Bloss et al., 2016; Sandler et al., 2016; Morgan et al., 2016) or local glial networks (Perea et al., 2009). They may also be influenced by the integrated nature of higher-level brain systems, including mechanisms for developmental bootstrapping (Ullman et al., 2012), information routing (Gurney et al., 2001; Stocco et al., 2010), attention (Buschman and Miller, 2010) and hierarchical decision making (Lee et al., 2015). Mapping these systems in detail is of paramount importance to understanding how the brain works, down to the nanoscale dendritic organization of ion channels and up to the real-time global coordination of cortex, striatum and hippocampus, all of which are computationally relevant in the framework we have explicated here. We thus expect that large-scale, multi-resolution brain maps would be useful in testing these framework-level ideas, in inspiring their refinements, and in using them to guide more detailed analysis.

### 5.2 *Hypothesis* 2– Biological fine-structure of cost functions

Clearly, we can map differences in structure, dynamics and representation across brain areas. When we find such differences, the question remains as to whether we can interpret these as resulting from differences in the internally-generated cost functions, as opposed to differences in the input data, or from differences that reflect other constraints unrelated to cost functions. If we can directly measure aspects of the cost function in different areas, then we can also compare them across areas. For example, methods from inverse reinforcement learning^60^ might allow backing out the cost function from observed plasticity (Ng and Russell, 2000).

Moreover, as we begin to understand the “neural correlates” of particular cost functions – perhaps encoded in particular synaptic or neuromodulatory learning rules, genetically-guided local wiring patterns, or patterns of interaction between brain areas – we can also begin to understand when differences in observed neural circuit architecture reflect differences in cost functions.

We expect that, for each distinct learning rule or cost function, there may be specific molecularly identifiable types of cells and/or synapses. Moreover, for each specialized system there may be specific molecularly identifiable developmental programs that tune it or otherwise set its parameters. This would make sense if evolution has needed to tune the parameters of one cost function without impacting others.

How many different types of internal training signals does the brain generate? When thinking about error signals, we are not just talking about dopamine and serotonin, or other classical reward-related pathways. The error signals that may be used to train specific sub-networks in the brain, via some approximation of gradient descent or otherwise, are not necessarily equivalent to reward signals. It is important to distinguish between cost functions that may be used to drive optimization of specific sub-circuits in the brain, and what are referred to as “value functions” or “utility functions”, i.e., functions that predict the agent’s aggregate future reward. In both cases, similar reinforcement learning mechanisms may be used, but the interpretation of the cost functions is different. We have not emphasized global utility functions for the animal here, since they are extensively studied elsewhere (e.g., (O’Reilly et al., 2014a; Bach, 2015)), and since we argue that, though important, they are only a part of the picture, i.e., that the brain is not solely an end-to-end reinforcement trained system.

Progress in brain mapping could soon allow us to classify the types of reward signals in the brain, follow the detailed anatomy and connectivity of reward pathways throughout the brain, and map in detail how reward pathways are integrated into striatal, cortical, hippocampal and cerebellar microcircuits. This program is beginning to be carried out in the fly brain, in which twenty specific types of dopamine neuron project to distinct anatomical compartments of the mushroom body to train distinct odor classifiers operating on a set of high-dimensional odor representations (Aso et al., 2014a, b; Caron et al., 2013; Cohn et al., 2015). It is known that, even within the same system, such as the fly olfactory pathway, some neuronal wiring is highly specific and molecularly programmed (Hong and Luo, 2014; Hattori et al., 2007), while other wiring is effectively random (Caron et al., 2013), and yet other wiring is learned (Aso et al., 2014a). The interplay between such design principles could give rise to many forms of “division of labor” between genetics and learning. Likewise, it is believed that birdsong learning is driven by reinforcement learning using a specialized cost function that relies on comparison with a memorized version of a tutor’s song (Fiete et al., 2007), and also that it involves specialized structures for controlling song variability during learning (Aronov et al., 2011). These detailed pathways underlying the construction of cost functions for vocal learning are beginning to be mapped (Mandelblat-Cerf et al., 2014). Starting with simple systems, it should become possible to map the reward pathways and how they evolved and diversified, which would be a step on the way to understanding how the system learns.

These types of mapping efforts would be a first step towards the ability to create a concrete model of the brain’s optimization architecture. Our discussion here has focused on trying to anticipate, based on known neuroscience knowledge and on approaches becoming successful in machine learning, the *kinds* of local cost functions that the brain may rely on, and how specialized brain systems may enable efficient solutions to optimization problems. However, this framework-level discussion is not a formal specification, either of the architecture, or of a notion of biologically applied cost function that could be directly measured based on neural data. In order to move towards a more formal specification of the kind of model we are proposing here, it would be useful to map the architecture of the brains reward systems and to identify other biological pathways that may mediate the generation and delivery of error signals. Based on such maps, one could identify regions which are proposed to be subject to a single cost function. Otherwise, the problem of inference of the cost function, e.g., based on neural dynamics becomes ill-posed: one can define a local cost function for an *arbitrary* dynamics by integrating the trajectory of the system, but this approach in general lacks explanatory power and also, crucially, lacks any circuit-level relationship with the brains actual neural mechanisms of optimization, i.e., such a defined cost function does not necessarily correspond to the cost functions that the biological machinery is actually organized to optimize. Notably, some of the relevant biological pathways mediating cost functions and error signals may involve key biomolecular or gene expression aspects, not just real-time patterns of neural activity.

Another related consideration, in trying to formalize this type approach and to infer cost functions from neural measurements, is that not all neurons in the circuit may be subject to optimization: after all, some neurons may be needed to generate the error signals themselves, or to mediate the optimization process for other neurons, or to perform other unrelated functions. Furthermore, within a given region, there may be multiple sub-circuits subject to different optimization pressures. It is the claim that the brain actually has structured biological machinery to generate, route and apply specific cost functions that gives substance to our proposal, over and above the trivial claim that many kinds of dynamics can be viewed as optimizations, but our knowledge of this machinery is still limited. This is not to mention the difficulties involved in inferring cost functions in the presence of noise or constraints on the dynamics. Thus, one cannot blindly collect the neurons in an arbitrary region, measure their dynamics, and hope to infer their cost function by solving an inverse problem – instead, a rich interplay between structural mapping, dynamic mapping, hypothesis generation, modeling and perturbation is likely to be necessary in order to gain a detailed knowledge of which cost functions the brain uses and how it does so.

### 5.3 *Hypothesis 3*– Embedding within a pre-structured architecture

If different brain structures are performing distinct types of computations with a shared goal, then optimization of a joint cost function will take place with different dynamics in each area. If we focus on a higher level task, e.g., maximizing the probability of correctly detecting something, then we should find that basic feature detection circuits should learn when the features were insufficient for detection, that attentional routing structures should learn when a different allocation of attention would have improved detection and that memory structures should learn when items that matter for detection were not remembered. If we assume that multiple structures are participating in a joint computation, which optimizes an overall cost function (but see **Hypothesis 2**), then an understanding of the computational function of each area leads to a prediction of the measurable plasticity rules.

## 6. Neuroscience inspired machine learning

Machine learning may be equally transformed by neuroscience. Within the brain, a myriad of subsystems and layers work together to produce an agent that exhibits general intelligence. The brain is able to show intelligent behavior across a broad range of problems using only relatively small amounts of data. As such, progress at understanding the brain promises to improve machine learning. In this section, we review our three hypotheses about the brain and discuss how their elaboration might contribute to more powerful machine learning systems.

### 6.1 *Hypothesis 1*– Existence of cost functions

A good practitioner of machine learning should have a broad range of optimization methods at their disposal as different problems ask for different approaches. The brain, we have argued, is an implicit machine learning mechanism which has been evolved over millions of years. Consequently, we should expect the brain to be able to optimize cost functions efficiently, across many domains and kinds of data. Indeed, across different animal phyla, we even see *convergent* evolution of certain brain structures (Shimizu and Karten, 2013; Güntürkün and Bugnyar, 2016), e.g., the bird brain has no cortex yet has developed homologous structures which – as the linguistic feats of the African Grey Parrot demonstrate – can give rise to quite complex intelligence. It seems reasonable to hope to learn how to do truly general-purpose optimization by looking at the brain.

Indeed, there are multiple kinds of optimization that we may expect to discover by looking at the brain. At the hardware level, the brain clearly manages to optimize functions efficiently despite having slow hardware subject to molecular fluctuations, suggesting directions for improving the hardware of machine learning to be more energy efficient. At the level of learning rules, the brain solves an optimization problem in a highly nonlinear, non-differentiable, temporally stochastic, spiking system with massive numbers of feedback connections, a problem that we arguably still do not know how to efficiently solve for neural networks. At the architectural level, the brain can optimize certain kinds of functions based on very few stimulus presentations, operates over diverse timescales, and clearly uses advanced forms of active learning to infer causal structure in the world.

While we have discussed a range of theories (Hinton, 2007, 2016; Bengio et al., 2015a; Balduzzi et al., 2014; Roelfsema et al., 2010; O’Reilly, 1996; O’Reilly et al., 2014a; Körding and König, 2001; Lillicrap et al., 2014) for how the brain can carry out optimization, these theories are still preliminary. Thus, the first step is to understand whether the brain indeed performs multi-layer credit assignment in a manner that approximates full gradient descent, and if so, how it does this. Either way, we can expect that answer to impact machine learning. If the brain does *not* do some form of back-propagation, this suggests that machine learning may benefit from understanding the tricks that the brain uses to avoid having to do so. If, on the other hand, the brain does do backpropagation, then the underlying mechanisms clearly can support a very wide range of efficient optimization processes across many domains, including learning from rich temporal data-streams and via unsupervised mechanisms, and the architectures behind this will likely be of long-term value to machine learning^61^. Moreover, the search for biologically plausible forms of backpropagation has already led to interesting insights, such as the possibility of using random feedback weights (feedback alignment) in backpropagation (Lillicrap et al., 2014), or the unexpected power of internal FORCE learning in chaotic, spontaneously active recurrent networks (Sussillo and Abbott, 2009). This and other findings discussed here suggest that there are still fundamental things we don’t understand about backpropagation – which could potentially lead not only to more biologically plausible ways to train recurrent neural networks, but also to fundamentally simpler and more powerful ones.

### 6.2 *Hypothesis 2*– Biological fine-structure of cost functions

A good practitioner of machine learning has access to a broad range of learning techniques and thus implicitly is able to use many different cost functions. Some problems ask for clustering, others for extracting sparse variables, and yet others for prediction quality to be maximized. The brain also needs to be able to deal with many different kinds of datasets. As such, it makes sense for the brain to use a broad range of cost functions appropriate for the diverse set of tasks it has to solve to thrive in this world.

Many of the most notable successes of deep learning, from language modeling (Sutskever et al., 2011), to vision (Krizhevsky et al., 2012), to motor control (Levine et al., 2015), have been driven by end-to-end optimization of single task objectives. We have highlighted cases where machine learning has opened the door to multiplicities of cost functions that shape network modules into specialized roles. We expect that machine learning will increasingly adopt these practices in the future.

In computer vision, we have begun to see researchers re-appropriate neural networks trained for one task (e.g., ImageNet classification) and then deploy them on new tasks other than the ones they were trained for or for which more limited training data is available (Yosinski et al., 2014; Oquab et al., 2014; Noroozi and Favaro, 2016). We imagine this procedure will be generalized, whereby, in series and in parallel, diverse training problems, each with an associated cost function, are used to shape visual representations. For example, visual data streams can be segmented into elements like foreground vs. background, objects that can move of their own accord vs. those that cannot, all using diverse unsupervised criteria (Ullman et al., 2012; Poggio, 2015). Networks so trained can then be shared, augmented, and retrained on new tasks. They can be introduced as front-ends for systems that perform more complex objectives or even serve to produce cost functions for training other circuits (Watter et al., 2015). As a simple example, a network that can discriminate between images of different kinds of architectural structures (pyramid, staircase, etc.) could act as a critic for a building-construction network.

Scientifically, determining the order in which cost functions are engaged in the biological brain will inform machine learning about how to construct systems with intricate and hierarchical behaviors via divide-and-conquer approaches to learning problems, active learning, and more.

### 6.3 *Hypothesis 3*– Embedding within a pre-structured architecture

A good practitioner of machine learning should have a broad range of algorithms at their disposal. Some problems are efficiently solved through dynamic programming, others through hashing, and yet others through multi-layer back-propagation. The brain needs to be able to solve a broad range of learning problems without the luxury of being reprogrammed. As such, it makes sense for the brain to have specialized structures that allow it to rapidly learn to approximate a broad range of algorithms.

The first neural networks were simple single-layer systems, either linear or with limited nonlinearities (Rashevsky, 1939). The explosion of neural network research in the 1980s (McClelland et al., 1986) saw the advent of multilayer networks, followed by networks with layer-wise specializations as in convolutional networks (Fukushima, 1980; LeCun and Bengio, 1995). In the last two decades, architectures with specializations for holding variables stable in memory like the LSTM (Hochreiter and Schmidhuber, 1997), the control of content-addressable memory (Weston et al., 2014; Graves et al., 2014), and game playing by reinforcement learning (Mnih et al., 2015) have been developed. These networks, though formerly exotic, are now becoming mainstream algorithms in the toolbox of any deep learning practitioner. There is no sign that progress in developing new varieties of structured architectures is halting, and the heterogeneity and modularity of the brain’s circuitry suggests that diverse, specialized architectures are needed to solve the diverse challenges that confront a behaving animal.

The brain combines a jumble of specialized structures in a way that works. Solving this problem *de novo* in machine learning promises to be very difficult, making it attractive to be inspired by observations about how the brain does it. An understanding of the breadth of specialized structures, as well as the architecture that combines them, should be quite useful.

## 7. Did evolution separate cost functions from optimization algorithms?

Deep learning methods have taken the field of machine learning by storm. Driving the success is the separation of the problem of learning into two pieces: (**1**) An algorithm, backpropagation, that allows efficient distributed optimization, and (**2**) Approaches to turn any given problem into an optimization problem, by designing a cost function and training procedure which will result in the desired computation. If we want to apply deep learning to a new domain, e.g., playing Jeopardy, we do not need to change the optimization algorithm – we just need to cleverly set up the right cost function. A lot of work in deep learning, perhaps the majority, is now focused on setting up the right cost functions.

We hypothesize that the brain also acquired such a separation between optimization mechanisms and cost functions. If neural circuits, such as in cortex, implement a general-purpose optimization algorithm, then any improvement to that algorithm will improve function across the cortex. At the same time, different cortical areas solve different problems, so tinkering with each area’s cost function is likely to improve its performance. As such, functionally and evolutionarily separating the problems of optimization and cost function generation could allow evolution to produce better computations, faster. For example, common unsupervised mechanisms could be combined with area-specific reinforcement-based or supervised mechanisms and error signals, much as recent advances in machine learning have found natural ways to combine supervised and unsupervised objectives in a single system (Rasmus and Berglund, 2015).

This suggests interesting questions^62^: When did the division between cost functions and optimization algorithms occur? How is this separation implemented? How did innovations in cost functions and optimization algorithms evolve? And how do our own cost functions and learning algorithms differ from those of other animals?

There are many possibilities for how such a separation might be achieved in the brain. Perhaps the six-layered cortex represents a common optimization algorithm, which in different cortical areas is supplied with different cost functions. This claim is different from the claim that all cortical areas use a single unsupervised learning algorithm and achieve functional specificity by tuning the inputs to that algorithm. In that case, both the optimization mechanism and the implicit unsupervised cost function would be the same across areas (e.g., minimization of prediction error), with only the training data differing between areas, whereas in our suggestion, the optimization mechanism would be the same across areas but the cost function, *as well as* the training data, would differ. Thus the cost function itself would be like an ancillary input to a cortical area, in addition to its input and output data. Some cortical microcircuits could then, perhaps, compute the cost functions that are to be delivered to other cortical microcircuits. Another possibility is that, within the same circuitry, certain aspects of the wiring and learning rules specify an optimization mechanism and are relatively fixed across areas, while others specify the cost function and are more variable. This latter possibility would be similar to the notion of cortical microcircuits as molecularly and structurally configurable elements, akin to the cells in a field-programmable gate array (FPGA) (Marcus et al., 2014a, b), rather than a homogenous substrate. The biological nature of such a separation, if any exists, remains an open question. For example, individual parts of a neuron may separately deal with optimization and with the specification of the cost function, or different parts of a microcircuit may specialize in this way, or there may be specialized types of cells, some of which deal with signal processing and others with cost functions.

## 8. Conclusions

Due to the complexity and variability of the brain, pure “bottom up” analysis of neural data faces potential challenges of interpretation (Robinson, 1992; Jonas and Kording, 2016). Theoretical frameworks can potentially be used to constrain the space of hypotheses being evaluated, allowing researchers to first address higher-level principles and structures in the system, and then “zoom in” to address the details. Proposed “top down” frameworks for understanding neural computation include entropy maximization, efficient encoding, faithful approximation of Bayesian inference, minimization of prediction error, attractor dynamics, modularity, the ability to subserve symbolic operations, and many others (Bialek, 2002; Bialek et al., 2006; Friston, 2010; Knill and Pouget, 2004; Marcus, 2001; Pinker, 1999). Interestingly, many of the “top down” frameworks boil down to assuming that the brain simply optimizes a single, given cost function for a single computational architecture. We generalize these proposals assuming both a heterogeneous combination of cost functions unfolding over development, and a diversity of specialized sub-systems.

Much of neuroscience has focused on the search for “the neural code”, i.e., it has asked which stimuli are good at driving activity in individual neurons, regions, or brain areas. But, if the brain is capable of generic optimization of cost functions, then we need to be aware that rather simple cost functions can give rise to complicated stimulus responses. This potentially leads to a different set of questions. Are differing cost functions indeed a useful way to think about the differing functions of brain areas? How does the optimization of cost functions in the brain actually occur, and how is this different from the implementations of gradient descent in artificial neural networks? What additional constraints are present in the circuitry that remain fixed while optimization occurs? How does optimization interact with a structured architecture, and is this architecture similar to what we have sketched? Which computations are wired into the architecture, which emerge through optimization, and which arise from a mixture of those two extremes? To what extent are cost functions explicitly computed in the brain, versus implicit in its local learning rules? Did the brain evolve to separate the mechanisms involved in cost function generation from those involved in the optimization of cost functions, and if so how? What kinds of meta-level learning might the brain apply, to learn when and how to invoke different cost functions or specialized systems, among the diverse options available, to solve a given task? What crucial mechanisms are left out of this framework? A more in-depth dialog between neuroscience and machine learning could help elucidate some of these questions.

Much of machine learning has focused on finding ever faster ways of doing end-to-end gradient descent in neural networks. Neuroscience may inform machine learning at multiple levels. The optimization algorithms in the brain have undergone a couple of hundred million years of evolution. Moreover, the brain may have found ways of using heterogeneous cost functions that interact over development so as to simplify learning problems by guiding and shaping the outcomes of unsupervised learning. Lastly, the specialized structures evolved in the brain may inform us about ways of making learning efficient in a world that requires a broad range of computational problems to be solved over multiple timescales. Looking at the insights from neuroscience may help machine learning move towards general intelligence in a structured heterogeneous world with access to only small amounts of supervised data.

In some ways our proposal is opposite to many popular theories of neural computation. There is not one mechanism of optimization but (potentially) many, not one cost function but a host of them, not one kind of a representation but a representation of whatever is useful, and not one homogeneous structure but a large number of them. All these elements are held together by the optimization of internally generated cost functions, which allows these systems to make good use of one another. Rejecting simple unifying theories is in line with a broad range of previous approaches in AI. For example, Minsky and Papert’s work on the Society of Mind (Minsky, 1988) – and more broadly on ideas of genetically staged and internally bootstrapped development in connectionist systems (Minsky, 1977) – emphasizes the need for a system of internal monitors and critics, specialized communication and storage mechanisms, and a hierarchical organization of simple control systems.

At the time these early works were written, it was not yet clear that gradient-based optimization could give rise to powerful feature representations and behavioral policies. One can view our proposal as a renewed argument against simple end-to-end training and in favor of a heterogeneous approach. In other words, this framework could be viewed as proposing a kind of “society” of cost functions and trainable networks, permitting internal bootstrapping processes reminiscent of the Society of Mind (Minsky, 1988). In this view, intelligence is enabled by many computationally specialized structures, each trained with its own developmentally regulated cost function, where both the structures and the cost functions are themselves optimized by evolution like the hyperparameters in neural networks.

## Footnotes

1. Hyper-parameter optimization shows that complicated schedules of training, which differ across parts of the network, lead to optimal performance (Maclaurin et al., 2015).

2. In adversarial networks, a generator network is trained to fool a discriminator network into being unable to distinguish generated samples from real data samples, while the discriminator network is trained to prevent the generator network from fooling it in this way.

3. Psychologists have been quantifying the subtleties of many such developmental stagings, e.g., of our perceptual and motor performance, e.g., (Nardini et al., 2010; Dekker and Nardini, 2015; McKone et al., 2009).

4. Our point in this section will not be that all learning in the brain can be captured by cost function optimization, but rather, somewhat more narrowly, our claim is that the algorithms for optimization like backpropagation in deep learning may have correspondences in biological brains. We feel that it is an important task for neuroscience to determine whether and how brains implement these algorithms. The brain may also disclose dynamics that are unlike these algorithms, so we are not disclaiming the possibility of broader theories. In machine learning, many useful algorithms are not explicitly formulated as cost function optimization; for example, many algorithms are based on linear algebra procedures like singular value decomposition, rather than explicit optimization. Such methods can be made nonlinear by using nonlinear kernels – relatedly, some brain circuits run specialized computations using fixed nonlinear basis functions (e.g., in cerebellum). Moreover, while an implicit cost function can be attributed to account for many dynamical processes, as well as many popular learning algorithms, our claim is not merely that the brain uses other learning procedures that lead to solutions which implicitly minimize a cost function, but rather that it actually finds its solutions by performing a powerful form of optimization as such.

5. Of course, some circuits may also be heavily genetically pre-specified to minimize the burden on learning. For instance, particular cell adhesion molecules (Hattori et al., 2007) expressed on particular parts of particular neurons defined by a genetic cell type (Zeisel et al., 2015), and the detailed shapes and placements of neuronal arbors, may constrain connectivity in some cases, though in other cases local connectivity is thought to be only weakly constrained (Kalisman et al., 2005). Genetics is sufficient to specify complex circuits involving hundreds of neurons, such as central pattern generators (Yuste et al., 2005) which create complex self-stabilizing oscillations, or the entire nervous systems of small worms. Genetically guided wiring should not be thought of as fixed “hard-wiring” but rather as a programmatic construction process that can also accept external inputs and interact with learning mechanisms (Marcus, 2004).

6. Hebbian plasticity even has a well-understood biological basis in the form of the NMDA receptors, which are activated by the simultaneous occurrence of chemical transmitter delivered from the pre-synaptic neuron, and voltage depolarization of the post-synaptic neuron.

7. The variance can be mitigated by averaging out many perturbations before making a change to the baseline value of the weights, but this would take significant time for a network of non-trivial size as the variance of weight perturbation’s estimates scales in proportion to the number of synapses in the network.

8. If the error derivatives of the cost function with respect to the last layer of unit activities are unknown, then they can be replaced with node-perturbation-like correlations, as is common in reinforcement learning.

9. Interestingly, STDP is not a unitary phenomenon, but rather a diverse collection of different rules with different timescales and temporal asymmetries (Sjöström and Gerstner, 2010; Mishra et al., 2016). Effects include STDP with the inverse temporal asymmetry, symmetric STDP and STDP with different temporal window sizes. STDP is also frequency dependent, which can be explained by rules that depend on triplets rather than pairs of spikes (Pfister and Gerstner, 2006). In some cortical neurons, STDP even switches its sign as the synapse moves away from the neuron’s soma into the dendritic tree (Letzkus et al., 2006). While STDP is often included explicitly in models, biophysical derivations of STDP from various underlying phenomena are also being attempted, some of which involve the post-synaptic voltage (Clopath and Gerstner, 2010) or a local dendritic voltage (Urbanczik and Senn, 2014). Meanwhile, other theories suggest that STDP may enable the use of precise timing codes based on temporal coincidence of inputs, the generation and unsupervised learning of temporal sequences (Fiete et al., 2010; Abbott and Blum, 1996), enhancements to distal reward processing in reinforcement learning (Izhikevich, 2007), stabilization of neural responses (Kempter et al., 2001), or many other higher-level properties (Nessler et al., 2013; Kappel et al., 2014).

10. Hinton has suggested (Hinton, 2007, 2016) that this could take place in the context of autoencoders and recirculation (Hinton and McClelland, 1988). Bengio and colleagues have proposed (Scellier and Bengio, 2016; Bengio and Fischer, 2015; Bengio, 2014) another context in which the connection between STDP and plasticity rules that depend on the temporal derivative of the post-synaptic firing rate can be exploited for biologically plausible multilayer credit assignment. This setting relies on clamping of outputs and stochastic relaxation in energy-based models (Ackley et al., 1985), which leads to a continuous network dynamics (Hopfield, 1984) in which hidden units are perturbed towards target values (Bengio and Fischer, 2015), loosely similar to that which occurs in XCAL. This dynamics then allows the STDP-based rule to correspond to gradient descent on the energy function with respect to the weights (Scellier and Bengio, 2016). This scheme requires symmetric weights, but in an autoencoder context, Bengio notes that these can arise spontaneously (Arora et al., 2015).

11. Even BPTT has arguably not been completely successful in recurrent networks. The problems of vanishing and exploding gradients led to long short term memory networks with gated memory units. An alternative is to use optimization methods that go beyond first order derivatives (Martens and Sutskever, 2011). This suggests the need for specialized systems and structures in the brain to mitigate problems of temporal credit assignment.

12. Interestingly, the hippocampus seems to “time stamp” memories by encoding them into ensembles with cellular compositions and activity patterns that change gradually as a function of time on the scale of days (Rubin et al., 2015; Cai et al., 2016), and may use “time cells” to mark temporal positions within episodes on a timescale of seconds (Kraus et al., 2013).

13. Control theory concepts also appear to be useful for simplifying optimization problems in certain other settings (Todorov, 2009; Hennequin et al., 2014).

14. In one intriguing study of interval timing, single neurons exhibited response patterns over time which were scaled to the interval duration, and cooling the brain to slow down neural dynamics led to longer intervals being computed by the brain (Xu et al., 2014).

15. Analogs of weight perturbation and node perturbation are known for spiking networks (Seung, 2003; Fiete and Seung, 2006). Seung (2003) also discusses implications of gradient based learning algorithms for neuroscience, echoing some of our considerations here.

16. A related, but more general, question is how to learn over many layers of non-differentiable structures. One option is to perform updates via finite-sized rather than infinitesimal steps, e.g., via target-propagation (Bengio, 2014).

17. Eliasmith and others have shown (Eliasmith, 2013; Eliasmith and Anderson, 2004; Eliasmith et al., 2012) that complex functions and control systems can be compiled onto such networks, using nonlinear encoding and linear decoding of high-dimensional vectors.

18. Dendritic computation may also have other functions, e.g., competitive interactions between dendrites in a single neuron could also allow neurons to contribute to multiple different ensembles (Legenstein and Maass, 2011).

19. Localized activity in dendrites drives localized plasticity, with inhibitory interneurons, and interactions between inputs at different parts of the dendritic tree, controlling the local sign and spatial distribution of this plasticity (Cichon and Gan, 2015; Sjöström and Häusser, 2006).

20. In the model of (Körding and König, 2001), single spikes are used to transmit activations and burst spikes are used to transmit error information. In other models, including the dendritic voltage in a plasticity rule leads to error-driven and predictive learning that can approximate backpropagation *inside* a single complex neuron (in effect backpropagating from the net somatic output, through nonlinearities at the dendritic branch points, all the way back to the individual input synaptic weights) and that generalize to a reinforcement learning context (Urbanczik and Senn, 2014; Schiess et al., 2016). Single neurons with active dendrites and many synapses may also embody learning rules of greater complexity, such as the storage and recall of temporal patterns (Hawkins and Ahmad, 2016).

21. Interestingly, some connectomic studies are finding more obvious connectivity structure at the level of dendritic organization than at the cellular level (Morgan et al., 2016).

22. An interesting recent study explored this idea in the context of a model of modular cortical-column-like units (Piekniewski et al., 2016). Local units are multi-layer perceptrons trained to minimize a prediction error by gradient descent. Within each unit, predictive autoencoders form a data compression in their middle layers, which is then fed up to higher levels as well as laterally. This system is suggestive of the power of using modular units of intermediate complexity, each of which minimizes a prediction error locally, e.g., in a local few-layer network. The system currently uses a fixed format for transmission of vectors from one unit to another, but ideally the inter-module connectons should also be trained by gradient descent as well or by reinforcement learning rather than being fixed. The cortical-column-like modules could also be made more complex and could be organized into higher-order structures like Minsky’s semantic networks, frames and K-lines (Minsky, 1988) rather than in simple hierarchies, or such an architecture could self-organize via reinforcement learning or other mechanisms for defining inter-column connections. Such a system also needs connections with specific kinds of memory and long-range information routing systems.

23. This idea has been used by Hawkins and colleagues to suggest mechanisms for continuous online sequence learning (Hawkins and Ahmad, 2016; Cui et al., 2015) and by Larkum and colleagues for comparison of top-down and bottom-up signals (Larkum, 2013). The Larkum model focuses on the layer 5 (L5) pyramidal neuron type. The cell body of this neuron lies in L5 but extends its “apical” dendritic tree all the way up to a tuft at the top of the cortex in layer 1 (L1), which is a primary target of feedback projections. In the model, interactions between local spiking in these different dendritic zones, which are targeted by different kinds of projections, are crucial to the learning function. The model of Hawkins (Hawkins and Ahmad, 2016; Cui et al., 2015) also focused on the unique dendritic structure of the L5 pyramidal neuron, and distinguishes internal states of the neuron, which impact its responsiveness to other inputs, from activation states, which directly translate into spike rates. Three integration zones in each neuron, and dendritic NMDA spikes (Palmer et al., 2014) acting as local coincidence detectors (Shai et al., 2015), allow temporal patterns of dendritic input to impact the cell’s internal state. Intra-column inhibition is also used in this model. Other cortical models pay less attention to the details of dendritic computation, but still provide detailed interpretations of the inter-laminar projection patterns of the neocortex. For example, in (O’Reilly et al., 2014b), an architecture is presented for continuous learning based on prediction of the next input. Time is discretized into 100 millisecond bins via an alpha oscillation, and the deep vs. shallow layers maintain different information during these time bins, with deep layers maintaining a record of the previous time step, and shallow layers representing the current state. The stored information in the deep layers leads to a prediction of the current state, which is then compared with the actual current state. Periodic bursting locked to the oscillation provides a kind of clock that causes the current state to be shifted into the deep layers for maintenance during the subsequent time step, and recurrent loops with the thalamus allow this representation to remain stable for sufficiently long to be used to generate the prediction. Other theories utilize the biophysics of dendritic computation and spike timing dependent plasticity to explain how neurons could learn to make predictions (Brea et al., 2016) on a timescale of seconds using neurons with intrinsic plasticity time constants of a few tens of milliseconds.

24. I-theory can perhaps be viewed as a generalized alternative paradigm to the online optimization of cost functions via multi-layer gradient descent, as used in deep learning. It exploits similar network architectures as conventional deep learning, e.g., hierarchical convolutional networks for the case of feedforward vision, but rather than backpropagating errors, it uses local circuits and learning rules to store templates against which new inputs are compared. This relies on a theory of generalization in learning based on combinations of tuned units (Poggio and Bizzi, 2004), which has been applied to both vision and motor control. Neurons with the required Gaussian-like tunings to stored templates could be obtained through canonical, local, normalization-based circuits (Kouh and Poggio, 2008), which can also be tweaked to implement other aspects of a vision architecture like softmax operations and pooling.

25. One alternative picture that contrasts with straightforward cost function optimization emphasizes the types of computation that appear most naturally suited to heterogeneous, stochastic, noisy, continually changing neural circuitry (Maass). On this view, network plasticity is viewed as a sampling-based approximation to Bayesian inference (Kappel et al., 2015) where transiently changing synapses sample from a posterior distribution of network configurations, rather than as gradient descent on a cost function. This view emphasizes Monte-Carlo sampling procedures, rather than cost function optimization.

26. Sampling based inference procedures are used widely in Bayesian statistics, and efforts have been made to connect these procedures with circuit-based models of computations (Mansinghka and Jonas, 2014). It currently appears difficult, however, to reconcile generic Marcov Chain Monte Carlo (MCMC) dynamics, which mix slowly, with the fast time scales of human psychophysics. But Bayesian methods are powerful and come with a methodology for model comparison (Ghahramani, 2005). In machine learning, variational Bayesian methods have recently become popular precisely because they are capable of fast though approximate posterior inference (inferring causes from observables), but seem to be powerful enough to create strong models. For example, stochastic gradient descent optimization is beginning to be used for variational Bayesian inference (Kingma and Welling, 2013). Restricted Boltzmann Machines (RBMs) also achieve fast inference in shallow architectures – with only a small number of iterations of mixing required – but they do not mix quickly when stacked into deep hierarchies as deep Boltzmann machines. The greedy, layer-wise pre-training of a deep belief network (Hinton et al., 2006) provides a heuristic way to stack the RBMs by auto-encoding, but these have achieved less competitive results than current variational Bayesian models. The problem of fast inference in MCMC models is the subject of current research, including at the interface with biologically plausible models (Bengio et al., 2016). When these models are made to perform fast inference, they actually become somewhat similar to variational Bayesian methods, since they rely on feedforward approximate inference, at least to initialize the system.

27. Heuristics are widely used to simplify motor planning and control, e.g., (McLeod and Dienes, 1996).

28. Many other reinforcement learning algorithms, including REINFORCE (Williams, 1992), can be implemented as fully online algorithms using “eligibility traces”, which accumulate the sensitivity of action distributions to parameters in a temporally local manner (Sutton and Barto, 1998).

29. Zaremba and Sutskever (2015) also bridges reinforcement learning and backpropagation learning in the same system, in the context of a neural network controlling discrete interfaces, and illustrates some of the challenges of this approach: compared to an end-to-end backpropagation-trained Neural Turing Machine (Graves et al., 2014), reinforcement based training allows training of only relatively simple algorithmic tasks. Special measures need to be taken to make reinforcement efficient, including limiting the number of possible actions, subtracting a baseline reward, and training the network using a curriculum schedule.

30. This is distinct from a game-theoretic scenario in which multiple actors can achieve an equilibrium, e.g., (Gemp and Mahadevan, 2015).

31. Single neurons act as comparators in the motor system, e.g., (Brownstone et al., 2015), and networks in the retina adapt so as to report local differences in space or time rather than absolute values, a form of predictive coding (Hosoya et al., 2005).

32. Beginning with Hopfield’s definition of an energy function for inference in certain classes of symmetric network (Hopfield, 1982), researchers have discovered networks with inherent dynamics that implicitly optimizes certain objectives even while the connection weights are fixed, such as statistical reconstruction of the input via stochastic relaxation in Boltzmann machines (Ackley et al., 1985). Fast approximations of some of these inference procedures are perhaps biologically plausible and could rely on dendritic computation (Bengio et al., 2016). Iterative local Hebbian-like *learning* rules are often used to train the *weights* of such networks, without explicitly propagating error derivatives in the manner of backpropagation. In an appropriate network context, many other combinations of network dynamics and plasticity rules can give rise to inference and learning procedures that implicitly descend cost functions in activity space and/or weight space.

33. Dreams arguably illustrate that the brain uses generative models which also involve selective recall and recombination of episodic memories.

34. Much is known about the architecture of cortical feedback vs. feedforward connections. For example, canonically, feedforward connections project from superficial cortical layers to layer 4 of the recipient layer, while feedback connections terminate outside layer 4 and often originate in deeper layers. These types of relationships can be used anatomically to define the hierarchical organization of visual areas, as in (Felleman and Van Essen, 1991), although the original studies were performed in primates and the precise generalization to rodent cortex is not fully clear (Berezovskii et al., 2011), and there may be various alternate or overlapping anatomical pathways (Callaway, 2004), e.g., with some pathways involved in specific functions like gain control, others routed through specific gating mechanisms, and so forth. Advances in connectomics should allow this architecture to be studied more directly. The study of receptive field properties in the visual cortical hierarchy has led to many insights into this hierarchical system. For example, while each neuron in V1 has a classical local receptive field, neural responses at a given location in V1 also depend on visual locations far from the classical receptive field, e.g., through various forms of surround suppression. These studies have allowed an understanding of the spatial scales over which feedback connections operate in the early visual system (Angelucci et al., 2002). In particular, feedback connections are invoked to account for longer-range receptive field interactions, whereas horizontal connections are invoked to account for shorter-range receptive field interactions (Schwabe et al., 2006). Feedforward and feedback pathways are also distinguished dynamically, e.g., by propagating different oscillatory frequencies (Van Kerkoerle et al., 2014; Bastos et al., 2015), and moleculary, e.g., with NMDA receptors playing an important role in feedback processing.

35. Temporal continuity is exploited in Poggio (2015), which analyzes many properties of deep convolutional networks with respect to their biological plausibility, including their apparent need for large amounts of supervised training data, and concludes that the environment may in fact provide a sufficient number of “implicitly”, though not explicitly, labeled examples to train a deep convolutional network for object recognition. Implicit labeling of object identity, in this case, arises from temporal continuity: successive frames of a video are likely to have the same objects in similar places and orientations. This allows the brain to derive an invariant signature of object identity which is independent of transformations like translations and rotations, but which does not yet associate the object with a specific name or label. Once such an invariant signature is established, however, it becomes basically trivial to associate the signature with a label for classification (Anselmi et al., 2015). Poggio (2015) also suggests specific means, in the context of I-theory (Anselmi et al., 2015), by which this training could occur via the storage of image templates using Hebbian mechanisms among simple and complex cells in the visual cortex. Thus, in this model, the brain has used its implicit knowledge of the temporal continuity of object motion to provide a kind of minimal labeling that is sufficient to bootstrap object recognition. Although not formulated as a cost function, this shows how usefully the assumption of temporal continuity could be exploited by the brain.

36. Although, some multi-sensory integration appears to occur even in the early sensory cortices (Murray et al., 2012).

37. Other brain-inspired unsupervised objectives are being developed for unsupervised visual learning. One recent paper (Higgins et al., 2016) uses an objective function that seeks representations of statistically independent factors in images, by introducing a regularization term that pushes the distribution of latent factors learned in a generative model to be close to a unit Gaussian. This is based on a theory that the ventral visual stream is optimized to disentangle factors of variation in images.

38. In the visual system, it is still unknown why a clustered spatial pattern of representational categories arises, e.g., a physically localized “area” that seems to correspond to representations of faces (Kanwisher et al., 1997), another area for representations of visual word forms (McCandliss et al., 2003), and so on. It is also unknown why this spatial pattern seems to be largely reproducible across individuals. Some theories are based on bottom-up correlation-based clustering or neuronal competition mechanisms, which generate category-selective regions as a byproduct. Other theories suggest a computational reason for this organization, in the context of I-theory (Anselmi et al., 2015), involving the limited ability to generalize transformation-invariances learned for one class of objects to other classes (Leibo et al., 2015b). Areas for abstract culture-dependent concepts, like the visual word form area, suggest that the decomposition cannot be “purely genetic”. But it is conceivable that these areas could at least in part reflect different local cost functions.

39. Psychologists have postulated other innate heuristics, e.g., in the context of object tracking (Franconeri et al., 2012). That infant object concepts are trainable but only along certain dimensions (Scholl, 2004) also suggests the notion of a heuristically “guided” or “bootstrapped” learning process in this context.

40. Of course, specialized architecture also enters the picture at the level of the pre-structuring of trainable/optimizable modules themselves. Just as in deep learning, convolutional networks, LSTMs, residual networks and other specific architectures are used to make learning efficient and fast, even though more generic architectures like multilayer perceptrons or generally RNNs are universal function approximators.

41. It is interesting to consider how standard neural network models of vision would fit into this categorization. Consider convolutional neural networks, for example, with the convolutional filters optimized via supervised backpropagation. This is by no means a completely unstructured prior to backpropagation-based training. Indeed, these networks typically contain max-pooling and normalization layers with fixed computations that are not altered during learning, as well as fixed architectural features such as number and arrangement of layers, size and stride of the sliding window, and so forth. Likewise “hierarchical max-pooling” (HMAX) models (Serre et al., 2007) of the ventral stream are so-named because of these fixed architectural aspects. Thus, in a hypothetical biological implementation of such systems, these aspects would be pre-structured by genetics even if the convolutional weights would be trained via some kind of gradient descent optimization. There are some plausible neural circuits that would implement these standardized normalization and max pooling operations (Kouh and Poggio, 2008). Moreover, in a biological implementation, the machinery necessary to carry out the optimization itself would need to be embodied by appropriate, genetically structured circuitry.

42. Attractor models of memory in neuroscience tend to have the property that only one memory can be accessed at a time (although a brain can have many such memories that can be accessed in parallel). Recent machine learning systems, however, have constructed differentiable addressable memory (Graves et al., 2014) and gating (Whitney et al., 2016) systems by allowing weighted superpositions of memory registers or gates to be queried – it is unclear whether the brain uses such mechanisms.

43. Computational analogies have also been drawn between associative memory storage and object recognition (Leibo et al., 2015a), suggesting the possibility of closely related computations occurring in parts of neocortex and hippocampus. Indeed, the hippocampus and olfactory cortex (a more ancient and simpler structure than the neocortex (Shepherd, 2014; Fournier et al., 2015)) are few-layer structures described in comparative anatomy as “allocortex”, as opposed to the six-layered “neocortex”, and both types of cortex have some anatomical similarities (particularly for CA1 and subiculum, though less so for CA3 and dentate gyrus) such as the presence of pyramidal neurons. It has been suggested that the hippocampus can be thought of as the top of the cortical hierarchy (Hawkins and Blakeslee, 2007), responsible for handling and remembering information that could not be fully explained by lower levels of the hierarchy. These computational connections are still tentative.

44. Attention also arguably solves certain types of perceptual binding problem (Reynolds and Desimone, 1999).

45. The precise roles of synchrony in information routing and other processes, and when it should be viewed as a causal factor versus as an epiphenomenon of other mechanisms, is still being worked out. In some theories, oscillations occur as consequences of certain recurrent processing loops, e.g., thalamo-cortico-striatal loops (Eliasmith et al., 2012). In other models, so-called “dynamic circuit motifs”, involving specific combinations of cellular and synaptic sub-types, both generate synchronies (e.g., in part via intrinsically rhythmic pacemaker neurons) and exploit them for specific computational roles, particularly in the rapid dynamic formation of communication networks (Womelsdorf et al., 2014).

46. One idea for achieving such transferability is that of a partitionable (Hayworth, 2012) or annexable (Bostrom, 1996) network. These models posit that a large associative memory network links all the different buffers. This large associative memory network has a number of stable attractor states. These are called “global” attractor states since they link across all the buffers. Forcing a given buffer into an activity pattern resembling that of its corresponding “piece” of an attractor state will cause the entire global network to enter that global attractor state. During training, all of the connections between buffers are turned on, so that their learned contents, though not identical, are kept in correspondence by being part of the same attractor. Later, the connections between specific buffers can be turned off to allow them to store different information. Copy and paste is then implemented by turning on the connections between a source buffer and a destination buffer (Hayworth, 2012). Copying between a source and destination buffer can also be implemented, i.e., learned, in a deep learning system using methods similar to the addressing mechanisms of the Neural Turing Machine (Graves et al., 2014).

47. Micro-stimulation experiments, in which an animal learns to behaviorally report stimulation of electrode channels located in diverse cortical regions, suggest that many areas can be routed or otherwise linked to behavioral “outputs” (Histed et al., 2013), although the mechanisms behind this – e.g., whether this stimulation gives rise to a high-level percept that the animal then uses to make a decision – are unclear. Likewise, it is possible to reinforcement-train an animal to control the activity of individual neurons (Fetz, 1969, 2007).

48. Conventionally, models of the basal ganglia involve all or none gating of an action, but recent evidence suggests that the basal ganglia may also have continuous, analog outputs (Yttri and Dudman, 2016).

49. It has been suggested that the basic role of the BG is to provide tonic inhibition to other circuits (Grillner et al., 2005). Release of this inhibition can then activate a “discrete” action, such as a motor command. A core function of the BG is thus to choose, based on patterns detected in its input, which of a finite set of actions to initiate via such release of inhibition. In many models of the basal ganglias role in cognitive control, the targets of inhibition are thalamic relays (Sherman, 2005), which are set in a default “off” state by tonic inhibition from the basal ganglia. Upon disinhibition of a relay, information is transferred from one cortical location to another – a form of conditional “gating” of information transfer. For example, the BG might be able to selectively “clamp” particular groups of cortical neurons in a fixed state, while leaving others free to learn and adapt. It could thereby enforce complex training routines, perhaps similar to those used to force the emergence of disentangled representations in (Kulkarni et al., 2015). The idea that the basal ganglia can train the cortex is not new, and already appears to have considerable experimental and anatomical support (Ashby et al., 2007, 2010; Pasupathy and Miller, 2005; Turner and Desmurget, 2010).

50. Like many brain areas, the hippocampus is richly innervated by a variety of reward-related and other neuromodulatory systems (Hasselmo and Wyble, 1997; Verney et al., 1985; Colino and Halliwell, 1987).

51. It remains unclear whether place cells take input from the grid cell system or vice versa (Hasselmo, 2015).

52. Other spatial problems such as mental rotation may require learning architectures specialized for geometric coordinate transformations (Hinton et al., 2011; Jaderberg et al., 2015a) or binding mechanisms that support structural, compositional, parametric descriptions of a scene (Hayworth et al., 2011).

53. There is some direct fMRI evidence for anatomically separate registers representing the contents of different sentence roles in the human brain (Frankland and Greene, 2015), which is suggestive of a possible anatomical binding mechanism, but also consistent with other mechanisms like vector symbolic architectures. More generally, the substrates of symbolic processing in the brain may bear an intimate connection with the representation of objects in working memory in the prefrontal cortex, and specifically with the question of how the PFC represents multiple objects in working memory simultaneously. This question is undergoing extensive study in primates (Warden and Miller, 2007, 2010; Siegel et al., 2009; Rigotti et al., 2013).

54. There is controversy around claims that recursive syntax is also present in songbirds (Van Heijningen et al., 2009).

55. The above mechanisms are spontaneous and subconscious. In conscious thought, too, the brain can clearly visit the multiple layers of a program one after the other. We make high-level plans that we fill with lower-level plans. Humans also have memory for their own thought processes. We have some ability to put “on hold” our current state of mind, start a new train of thought, and then come back to our original thought. We also are able to ask, introspectively, whether we have had a given thought before. The neural basis of these processes is unclear, although one may speculate that the hippocampus is involved.

56. Fluorescent techniques like (Hayashi-Takagi et al., 2015) might be helpful.

57. The use of structured microcircuits rather than individual neurons as the units of learning can ease the burden on the learning rules possessed by individual neurons, as exemplified by a study implementing Helmholtz machine learning in a network of spiking neurons using conventional plasticity rules (Sountsov and Miller, 2015; Roudi and Taylor, 2015). As a simpler example, the classical problem of how neurons with only one output axon could communicate both activation and error derivatives for backpropagation ceases to be a problem if the unit of optimization is not a single neuron. Similar considerations hold for the issue of weight symmetry, or approximate sign-concordance in the case of feedback alignment (Liao et al., 2015).

58. Within this framework, networks that adhere to the basic statistics of neural connectivity, electrophysiology and morphology, such as the initial cortical column models from the Blue Brain Project (Markram et al., 2015), would recapitulate some properties of the cortex, but – just like untrained neural networks – would not spontaneously generate complex functional computation without being subjected to a multi-stage training process, naturalistic sensory data, signals arising from other brain areas and action-driven reinforcement signals.

59. Not only in applied machine learning, but also in today’s most advanced neuro-cognitive models such as SPAUN (Eliasmith, 2013; Eliasmith et al., 2012), the detailed local circuit connectivity is obtained through an optimization process of some kind to achieve a particular functionality. In the case of modern machine learning, training is often done via end-to-end backpropagation through an architecture that is only structured at the level of higher-level “blocks” of units, whereas in SPAUN each block is optimized (Eliasmith and Anderson, 2004) separately according to a procedure that allows the blocks to subsequently be stitched together in a coherent way. Technically, the Neural Engineering Framework (Eliasmith and Anderson, 2004) used in SPAUN uses singular value decomposition, rather than gradient descent, to compute the connections weights as optimal linear decoders. This is possible because of a nonlinear mapping into a high-dimensional space, in which approximating any desired function can be done via a hyperplane regression (Tapson and van Schaik, 2013).

60. There is a rich tradition of trying to estimate the cost function used by human beings (Ng and Russell, 2000; Finn et al., 2016; Ho and Ermon, 2016). The idea is that we observe (by stipulation) behavior that is optimal for the human’s cost function. We can then search for the cost function that makes the observed behavior most probable and simultaneously makes the behaviors that could have been observed, but were not, least probable. Extensions of such approaches could perhaps be used to ask which cost functions the brain is optimizing.

61. Successes of deep learning are already being used, speculatively, to rationalize features of the brain. It has been suggested that large networks, with many more neurons available than are strictly needed for the target computation, make learning easier (Goodfellow et al., 2014b). In concordance with this, visual cortex appears to be a 100-fold over-complete representation of the retinal output (Lewicki and Sejnowski, 2000). Likewise, it has been suggested that biological neurons stabilized (Turrigiano, 2012) to operate far below their saturating firing rates mirror the successful use of rectified linear units in facilitating the training of artificial neural networks (Roudi and Taylor, 2015). Hinton and others have also suggested a biological motivation (Roudi and Taylor, 2015) for “dropout” regularization (Srivastava et al., 2014), in which a fraction of hidden units is stochastically set to zero during each round of training: such a procedure may correspond to the noisiness of neural spike trains, although other theories interpret spikes as sampling in probabilistic inference (Buesing et al., 2011), or in many other ways. Randomness of spiking has some support in neuroscience (Softky and Koch, 1993), although recent experiments suggest that spike trains in certain areas may be less noisy than previously thought (Hires et al., 2015). The key role of proper initialization in enabling effective gradient descent is an important recent finding (Saxe et al., 2013; Sutskever and Martens, 2013) which may also be reflected by biological mechanisms of neural homeostasis or self-organization that would enforce appropriate initial conditions for learning. Retinal fixation has been tentatively connected with robustness of convolutional networks to adversarial perturbations in images (Luo et al., 2015). But making these speculative claims of biological relevance more rigorous will require researchers to first evaluate *whether* biological neural circuits are performing multi-layer optimization of cost functions in the first place.

62. It would be interesting to study these questions in specific brain systems. The primary visual cortex, for example, is still only understood very incompletely (Olshausen and Field, 2004). It serves as a key input modality to both the ventral and dorsal visual pathways, one of which seems to specialize in object identity and the other in motion and manipulation. Higher-level areas like STP draw on both streams to perform tasks like complex action recognition. In some models (e.g., (Jhuang et al., 2007)), both ventral and dorsal streams are structured hierarchically, but the ventral stream primarily makes use of the spatial filtering properties of V1, whereas the dorsal stream primarily makes use of its spatio-*temporal* filtering properties, e.g., temporal frequency filtering by the space-time receptive fields of V1 neurons. Given this, we can ask interesting questions about V1. Within a framework of multilayer optimization, do both dorsal and ventral pathways impose cost functions that help to shape V1’s response properties? Or is V1 largely pre-structured by genetics and local self-organization, with different optimization principles in the ventral and dorsal streams only having effects at higher levels of the hierarchy? Or, more likely, is there some interplay between pre-structuring of the V1 circuitry and optimization according to multiple cost functions? Relatedly, what establishes the differing roles of the downstream ventral vs. dorsal cortical areas, and can their differences be attributed to differing cost functions? This relates to ongoing questions about the basic nature of cortical circuitry. For example, DiCarlo et al. (2012) suggests that visual cortical regions containing on the order of 10000 neurons are locally optimized to perform disentangling of the manifolds corresponding to their local views of the transformations of an object, allowing these manifolds to be linearly separated by readout areas. Yet, DiCarlo et al. (2012) also emphasizes the possibility that certain computations such as normalization are pre-initialized in the circuitry prior to learning-based optimization.

## Acknowledgments

We thank Ken Hayworth for key discussions that led to this paper. We thank Ed Boyden, Chris Eliasmith, Gary Marcus, Shimon Ullman, Tomaso Poggio, Josh Tenenbaum, Dario Amodei, Alex Williams, Erik Peterson, Tom Dean, David Sussillo, Matthew Botvinick, Joscha Bach, Mohammad Gheshlaghi Azar, Joshua Glaser, Marco Nardini, Ali Hummos, David Markowitz, David Rolnick, Sam Rodriques, Nick Barry, Matthew Larkum, Walter Senn, Eric Drexler, Vikash Mansinghka, Darcy Wayne, Lyra and Neo Marblestone, and all of the participants of a Kavli Salon on Cortical Computation (Feb/Oct 2015) for helpful comments. We thank Miles Brundage for an excellent Twitter feed of deep learning papers.

## References

LF Abbott and KI Blum. Functional significance of long-term potentiation for sequence learning and prediction. Cerebral Cortex, 1996. URL http://cercor.oxfordjournals.org/content/6/3/406.short.

LF Abbott, B DePasquale, and RM Memmesheimer. Building Functional Networks of Spiking Model Neurons. neurotheory.columbia.edu, 2016. URL http://www.neurotheory.columbia.edu/Larry/SpikingNetworkReview.pdf.

DH Ackley, GE Hinton, and TJ Sejnowski. A learning algorithm for Boltzmann machines. Cognitive science, 1985. URL http://www.sciencedirect.com/science/article/pii/S0364021385800124.

Daniel E Acuna, Nicholas F Wymbs, Chelsea A Reynolds, Nathalie Picard, Robert S Turner, Peter L Strick, Scott T Grafton, and Konrad P Kording. Multifaceted aspects of chunking enable robust algorithms. Journal of neurophysiology, 112(8): 1849–1856, 2014.

Guillaume Alain, Alex Lamb, Chinnadhurai Sankar, Aaron Courville, and Yoshua Bengio. Variance reduction in sgd by distributed importance sampling. arXiv preprint arXiv:1511.06481, 2015.

K Allen, S Ibara, and A Seymour. Abstract structural representations of goal-directed behavior. Psychological …, 2010. URL http://pss.sagepub.com/content/21/10/1518.short.

Charles H Anderson and David C Van Essen. Shifter circuits: a computational strategy for dynamic aspects of visual processing. Proceedings of the National Academy of Sciences, 84(17): 6297–6301, 1987.

John R Anderson. How Can the Human Mind Occur in the Physical Universe? Oxford University Press, 2007. ISBN 0198043538. URL http://books.google.com/books?id=eYZXEtfplyAC{&}pgis=1.

Jacob Andreas, Marcus Rohrbach, Trevor Darrell, and Dan Klein. Deep compositional question answering with neural module networks. arXiv preprint arXiv:1511.02799, 2015.

Jacob Andreas, Marcus Rohrbach, Trevor Darrell, and Dan Klein. Learning to compose neural networks for question answering. arXiv preprint arXiv:1601.01705, 2016.

Alessandra Angelucci, Jonathan B Levitt, Emma JS Walton, Jean-Michel Hupe, Jean Bullier, and Jennifer S Lund. Circuits for local and global signal integration in primary visual cortex. The Journal of Neuroscience, 22(19): 8633–8646, 2002.

Fabio Anselmi, Joel Z. Leibo, Lorenzo Rosasco, Jim Mutch, Andrea Tacchetti, and Tomaso Poggio. Unsupervised learning of invariant representations. Theoretical Computer Science, jun 2015. ISSN 03043975. doi: 10.1016/j.tcs.2015.06.048. URL http://www.sciencedirect.com/science/article/pii/S0304397515005587.

Srdjan D Antic, Wen-Liang Zhou, Anna R Moore, Shaina M Short, and Katerina D Ikonomu. The decade of the dendritic nmda spike. Journal of neuroscience research, 88(14): 2991–3001, 2010.

O Arancio, M Kiebler, C J Lee, V Lev-Ram, R Y Tsien, E R Kandel, and R D Hawkins. Nitric oxide acts directly in the presynaptic neuron to produce long-term potentiation in cultured hippocampal neurons. Cell, 87(6): 1025–35, dec 1996. ISSN 0092–8674. URL http://www.ncbi.nlm.nih.gov/pubmed/8978607.

Dmitriy Aronov, Lena Veit, Jesse H Goldberg, and Michale S Fee. Two distinct modes of forebrain circuit dynamics underlie temporal patterning in the vocalizations of young songbirds. The Journal of neuroscience : the official journal of the Society for Neuroscience, 31(45): 16353–68, nov 2011. ISSN 1529–2401. doi: 10.1523/JNEUROSCI.3009-11. 2011.

Sanjeev Arora, Yingyu Liang, and Tengyu Ma. Why are deep nets reversible: A simple theory, with implications for training. nov 2015. URL http://arxiv.org/abs/1511.05653.

F Gregory Ashby, John M Ennis, and Brian J Spiering. A neurobiological theory of automaticity in perceptual categorization. Psychological review, 114(3): 632, 2007.

F Gregory Ashby, Benjamin O Turner, and Jon C Horvitz. Cortical and basal ganglia contributions to habit learning and automaticity. Trends in cognitive sciences, 14(5): 208–15, may 2010. ISSN 1879–307X. doi: 10.1016/j.tics.2010.02.001. URL http://www.cell.com/article/S1364661310000306/fulltext.

Yoshinori Aso, Daisuke Hattori, Yang Yu, Rebecca M Johnston, Nirmala A Iyer, Teri-T B Ngo, Heather Dionne, L F Abbott, Richard Axel, Hiromu Tanimoto, and Gerald M Rubin. The neuronal architecture of the mushroom body provides a logic for associative learning. eLife, 3:e04577, jan 2014a. ISSN 2050–084X. doi: 10.7554/eLife.04577. URL http://elifesciences.org/content/3/e04577.abstract.

Yoshinori Aso, Divya Sitaraman, Toshiharu Ichinose, Karla R Kaun, Katrin Vogt, Ghislain Belliart-Guérin, Pierre-Yves Plaçais, Alice A Robie, Nobuhiro Yamagata, Christopher Schnaitmann, William J Rowell, Rebecca M Johnston, Teri-T B Ngo, Nan Chen, Wyatt Korff, Michael N Nitabach, Ulrike Heberlein, Thomas Preat, Kristin M Branson, Hiromu Tanimoto, and Gerald M Rubin. Mushroom body output neurons encode valence and guide memory-based action selection in Drosophila. eLife, 3:e04580, jan 2014b. ISSN 2050–084X. doi: 10.7554/eLife.04580. URL http://elifesciences.org/content/3/e04580.abstract.

J Ba and R Caruana. Do deep nets really need to be deep? Advances in neural information processing…, 2014. URL http://papers.nips.cc/paper/5484-do-deep-nets-really-need-to-be-deep.

Joscha Bach. Modeling Motivation in MicroPsi 2. Artificial General Intelligence, 2015. URL http://link.springer.com/chapter/10.1007/978–3–319–21365–1{_}1.

Joscha Bach and Priska Herger. Request confirmation networks for neuro-symbolic script execution. In Tarek Besold, Artur dAvila Garcez, Gary Marcus, and Risto Miikkulainen, editors, Workshop on Cognitive Computation: Integrating Neural and Symbolic Approaches at NIPS, 2015. URL http://www.neural-symbolic.org/CoCo2015/.

Renée Baillargeon, Rose M Scott, and Lin Bian. Psychological reasoning in infancy. Annual review of psychology, 67:159–186, 2016.

Jan Balaguer, Hugo Spiers, Demis Hassabis, and Christopher Summerfield. Neural Mechanisms of Hierarchical Planning in a Virtual Subway Network. Neuron, 90(4): 893–903, may 2016. ISSN 08966273. doi: 10.1016/j.neuron.2016.03.037. URL http://www.cell.com/article/S0896627316300575/fulltext.

Daniel Baldauf and Robert Desimone. Neural mechanisms of object-based attention. Science (New York, N.Y.), 344(6182): 424–7, apr 2014. ISSN 1095–9203. doi: 10.1126/science.1247003. URL http://www.ncbi.nlm.nih.gov/pubmed/24763592.

Pierre Baldi and Peter Sadowski. The Ebb and Flow of Deep Learning: a Theory of Local Learning. jun 2015. URL http://arxiv.org/abs/1506.06472.

D Balduzzi. Cortical prediction markets. Proceedings of the 2014 international conference on …, 2014. URL http://dl.acm.org/citation.cfm?id=2617449.

David Balduzzi, Hastagiri Vanchinathan, and Joachim Buhmann. Kickback cuts Backprop’s red-tape: Biologically plausible credit assignment in neural networks. page 7, nov 2014. URL http://arxiv.org/abs/1411.6191.

Cornelia I Bargmann. Beyond the connectome: how neuromodulators shape neural circuits. BioEssays : news and reviews in molecular, cellular and developmental biology, 34(6): 458–65, jun 2012. ISSN 1521–1878. doi: 10.1002/bies.201100185. URL http://www.ncbi.nlm.nih.gov/pubmed/22396302.

Cornelia I Bargmann and Eve Marder. From the connectome to brain function. Nature Methods, 10(6): 483–490, may 2013. ISSN 1548–7091. doi: 10.1038/nmeth.2451. URL http://dx.doi.org/10.1038/nmeth.2451.

Andre Moraes Bastos, Julien Vezoli, Conrado Arturo Bosman, Jan-Mathijs Schoffelen, Robert Oostenveld, Jarrod Robert Dowdall, Peter De Weerd, Henry Kennedy, and Pascal Fries. Visual areas exert feedforward and feedback influences through distinct frequency channels. Neuron, 85(2): 390–401, 2015.

Peter W Battaglia, Jessica B Hamrick, and Joshua B Tenenbaum. Simulation as an engine of physical scene understanding. Proceedings of the National Academy of Sciences of the United States of America, 110(45): 18327–32, nov 2013. ISSN 1091–6490. doi: 10.1073/pnas.1306572110. URL http://www.pnas.org/content/110/45/18327.short.

Suzanna Becker and Geoffrey E Hinton. Self-organizing neural network that discovers surfaces in random-dot stereograms. Nature, 355(6356): 161–163, 1992.

Trevor Bekolay, Carter Kolbeck, and Chris Eliasmith. Simultaneous unsupervised and supervised learning of cognitive functions in biologically plausible spiking neural networks. In Proceedings of the 35th annual conference of the cognitive science society, pages 169–174, 2013.

Yoshua Bengio. How Auto-Encoders Could Provide Credit Assignment in Deep Networks via Target Propagation. jul 2014. URL http://arxiv.org/abs/1407.7906.

Yoshua Bengio and Asja Fischer. Early Inference in Energy-Based Models Approximates Back-Propagation. oct 2015. URL http://arxiv.org/abs/1510.02777.

Yoshua Bengio, Jérôme Louradour, Ronan Collobert, and Jason Weston. Curriculum learning. In Proceedings of the 26th annual international conference on machine learning, pages 41–48. ACM, 2009.

Yoshua Bengio, Dong-Hyun Lee, Jorg Bornschein, and Zhouhan Lin. Towards Biologically Plausible Deep Learning. feb 2015a. URL http://arxiv.org/abs/1502.04156.

Yoshua Bengio, Thomas Mesnard, Asja Fischer, Saizheng Zhang, and Yuhuai Wu. STDP as presynaptic activity times rate of change of postsynaptic activity. sep 2015b. URL http://arxiv.org/abs/1509.05936.

Yoshua Bengio, Benjamin Scellier, Olexa Bilaniuk, Joao Sacramento, and Walter Senn. Feedforward initialization for fast inference of deep generative networks is biologically plausible. arXiv preprint arXiv:1606.01651, 2016.

Vladimir K Berezovskii, Jonathan J Nassi, and Richard T Born. Segregation of feedforward and feedback projections in mouse visual cortex. Journal of Comparative Neurology, 519(18): 3672–3683, 2011.

Robert C Berwick, Gabriël J L Beckers, Kazuo Okanoya, and Johan J Bolhuis. A Bird’s Eye View of Human Language Evolution. Frontiers in evolutionary neuroscience, 4:5, jan 2012. ISSN 1663–070X. doi: 10.3389/fnevo.2012.00005. URL http://journal.frontiersin.org/article/10.3389/fnevo.2012.00005/abstract.

W Bialek. Thinking about the brain. Physics of bio-molecules and cells. Physique des …, 2002. URL http://link.springer.com/chapter/10.1007/3–540–45701–1{_}12.

William Bialek, Rob De Ruyter Van Steveninck, and Naftali Tishby. Efficient representation as a design principle for neural coding and computation. In 2006 IEEE International Symposium on Information Theory, pages 659–663. IEEE, jul 2006. ISBN 1–4244–0505-X. doi: 10.1109/ISIT.2006.261867. URL http://ieeexplore.ieee.org/articleDetails.jsp?arnumber=4036045.

H Biilthoff, J Little, and T Poggio. A parallel algorithm for real-time computation of optical flow. Nature, 337(6207): 549–553, 1989.

ErikB. Bloss, MarkS. Cembrowski, Bill Karsh, Jennifer Colonell, RichardD. Fetter, and Nelson Spruston. Structured Dendritic Inhibition Supports Branch-Selective Integration in CA1 Pyramidal Cells. Neuron, 89(5): 1016–30, feb 2016. ISSN 08966273. doi: 10.1016/j.neuron.2016.01.029. URL http://www.cell.com/article/S0896627316000544/fulltext.

Charles Blundell, Benigno Uria, Alexander Pritzel, Yazhe Li, Avraham Ruderman, Joel Z Leibo, Jack Rae, Daan Wierstra, and Demis Hassabis. Model-free episodic control. arXiv preprint arXiv:1606.04460, 2016.

Bruce Bobier, Terrence C Stewart, and Chris Eliasmith. A unifying mechanistic model of selective attention in spiking neurons. PLoS Comput Biol, 10(6):e1003577, 2014.

Nick Bostrom. Cortical integration: Possible solutions to the binding and linking problems in perception, reasoning and long term memory. 1996. URL http://philpapers.org/rec/BOSCIP.

Matthew Botvinick and Ari Weinstein. Model-based hierarchical reinforcement learning and human action control. Philosophical transactions of the Royal Society of London. Series B, Biological sciences, 369(1655):20130480–, nov 2014. ISSN 1471–2970. doi: 10.1098/rstb.2013.0480. URL http://rstb.royalsocietypublishing.org/content/369/1655/20130480.

MM Botvinick, Y Niv, and AC Barto. Hierarchically organized behavior and its neural foundations: A reinforcement learning perspective. Cognition, 2009. URL http://www.sciencedirect.com/science/article/pii/S0010027708002059.

Guillaume Bouchard, Théo Trouillon, Julien Perez, and Adrien Gaidon. Accelerating stochastic gradient descent via online learning to sample. arXiv preprint arXiv:1506.09016, 2015.

Ralph Bourdoukan and Sophie Denève. Enforcing balance allows local supervised learning in spiking recurrent networks. In Advances in Neural Information Processing Systems, pages 982–990, 2015.

V Braitenberg and A Schutz. Anatomy of the cortex: studies of brain function, 1991.

Johanni Brea and Wulfram Gerstner. Does computational neuroscience need new synaptic learning paradigms? Current Opinion in Behavioral Sciences, 11:61–66, 2016.

Johanni Brea, Alexisz Tamás Gaál, Robert Urbanczik, and Walter Senn. Prospective coding by spiking neurons. PLOS Comput Biol, 12(6):e1005003, 2016.

J. Gavin Bremner, Alan M. Slater, and Scott P. Johnson. Perception of Object Persistence: The Origins of Object Permanence in Infancy. Child Development Perspectives, 9(1): 7–13, mar 2015. ISSN 17508592. doi: 10.1111/cdep.12098. URL http://doi.wiley.com/10.1111/cdep.12098.

Carlos SN Brito and Wulfram Gerstner. Nonlinear hebbian learning as a unifying principle in receptive field formation. arXiv preprint arXiv:1601.00701, 2016.

Tobias Brosch, Heiko Neumann, and Pieter R Roelfsema. Reinforcement Learning of Linking and Tracing Contours in Recurrent Neural Networks. PLoS computational biology, 11(10):e1004489, oct 2015. ISSN 1553–7358. doi: 10.1371/journal.pcbi.1004489.

Thackery I. Brown, Valerie A. Carr, Karen F. LaRocque, Serra E. Favila, Alan M. Gordon, Ben Bowles, Jeremy N. Bailenson, and Anthony D. Wagner. Prospective representation of navigational goals in the human hippocampus. Science, 352(6291): 1323–1326, 2016. ISSN 0036–8075. doi: 10.1126/science.aaf0784. URL http://science.sciencemag.org/content/352/6291/1323.

Robert M Brownstone, Tuan V Bui, and Nicolas Stifani. Spinal circuits for motor learning. Current opinion in neurobiology, 33:166–173, 2015.

Lars Buesing, Johannes Bill, Bernhard Nessler, and Wolfgang Maass. Neural dynamics as sampling: a model for stochastic computation in recurrent networks of spiking neurons. PLoS computational biology, 7(11):e1002211, nov 2011. ISSN 1553–7358. doi: 10.1371/journal.pcbi.1002211.

D V Buonomano and M M Merzenich. Temporal information transformed into a spatial code by a neural network with realistic properties. Science (New York, N.Y.), 267(5200): 1028–30, feb 1995. ISSN 0036–8075. URL http://www.ncbi.nlm.nih.gov/pubmed/7863330.

Timothy J Buschman and Earl K Miller. Shifting the spotlight of attention: evidence for discrete computations in cognition. Frontiers in human neuroscience, 4:194, jan 2010. ISSN 1662–5161. doi: 10.3389/fnhum.2010.00194.

Timothy J Buschman and Earl K Miller. Goal-direction and top-down control. Philosophical transactions of the Royal Society of London. Series B, Biological sciences, 369(1655), nov 2014. ISSN 1471–2970. doi: 10.1098/rstb. 2013.0471. URL http://www.ncbi.nlm.nih.gov/pubmed/25267814.

Keith A Bush. An echo state model of non-markovian reinforcement learning. 2007.

György Buzsáki. Neural syntax: cell assemblies, synapsembles, and readers. Neuron, 68(3): 362–385, 2010.

György Buzsáki and Edvard I Moser. Memory, navigation and theta rhythm in the hippocampal-entorhinal system. Nature neuroscience, 16(2): 130–138, feb 2013. ISSN 1546–1726. doi: 10.1038/nn.3304. URL http://dx.doi.org/10.1038/nn.3304.

Denise J Cai, Daniel Aharoni, Tristan Shuman, Justin Shobe, Jeremy Biane, Weilin Song, Brandon Wei, Michael Veshkini, Mimi La-Vu, Jerry Lou, et al. A shared neural ensemble links distinct contextual memories encoded close in time. Nature, 534(7605): 115–118, 2016.

EM Callaway. Feedforward, feedback and inhibitory connections in primate visual cortex. Neural Networks, 2004. URL http://www.sciencedirect.com/science/article/pii/S0893608004000887.

Sophie J C Caron, Vanessa Ruta, L F Abbott, and Richard Axel. Random convergence of olfactory inputs in the Drosophila mushroom body. Nature, 497(7447): 113–7, may 2013. ISSN 1476–4687. doi: 10.1038/nature12063. URL http://dx.doi.org/10.1038/nature12063.

Liang-Chieh Chen, Alexander G. Schwing, Alan L. Yuille, and Raquel Urtasun. Learning Deep Structured Models. page 11, jul 2014. URL http://arxiv.org/abs/1407.2538.

Sharat Chikkerur, Thomas Serre, Cheston Tan, and Tomaso Poggio. What and where: a Bayesian inference theory of attention. Vision research, 50 (22):2233–47, oct 2010. ISSN 1878–5646. doi: 10.1016/j.visres.2010.05.013. URL http://www.ncbi.nlm.nih.gov/pubmed/20493206.

Choo and Eliasmith. A Spiking Neuron Model of Serial-Order Recall. 32nd Annual Conference of the Cognitive Science Society, 2010. URL http://citeseerx.ist.psu.edu/viewdoc/summary?doi=10.1.1.353.1190.

Alexander A Chubykin, Emma B Roach, Mark F Bear, and Marshall G Hussain Shuler. A cholinergic mechanism for reward timing within primary visual cortex. Neuron, 77(4): 723–35, feb 2013. ISSN 1097–4199. doi: 10.1016/j.neuron.2012.12.039. URL http://www.cell.com/article/S0896627313000470/fulltext.

Junyoung Chung, Caglar Gulcehre, KyungHyun Cho, and Yoshua Bengio. Empirical Evaluation of Gated Recurrent Neural Networks on Sequence Modeling. dec 2014. URL http://arxiv.org/abs/1412.3555.

Joseph Cichon and Wen-Biao Gan. Branch-specific dendritic ca2+ spikes cause persistent synaptic plasticity. Nature, 520(7546): 180–185, 2015.

Nicola S Clayton and Anthony Dickinson. Episodic-like memory during cache recovery by scrub jays. Nature, 395(6699): 272–274, 1998.

Claudia Clopath and Wulfram Gerstner. Voltage and spike timing interact in stdp–a unified model. Spike-timing dependent plasticity, page 294, 2010.

Raphael Cohn, Ianessa Morantte, and Vanessa Ruta. Coordinated and Compartmentalized Neuromodulation Shapes Sensory Processing in Drosophila. Cell, 163(7): 1742–1755, dec 2015. ISSN 00928674. doi: 10.1016/j.cell.2015.11.019. URL http://www.cell.com/article/S0092867415014993/fulltext.

A Colino and JV Halliwell. Differential modulation of three separate k-conductances in hippocampal ca1 neurons by serotonin. Nature, 328(6125): 73–77, 1987.

Alexandra O Constantinescu, Jill X OReilly, and Timothy EJ Behrens. Organizing conceptual knowledge in humans with a gridlike code. Science, 352(6292): 1464–1468, 2016.

David Daniel Cox and Thomas Dean. Neural networks and neuroscience-inspired computer vision. Current biology : CB, 24(18):R921–9, sep 2014. ISSN 1879–0445. doi: 10.1016/j.cub.2014.08.026. URL http://www.sciencedirect.com/science/article/pii/S0960982214010392.

F Crick. The recent excitement about neural networks. Nature, 337(6203): 129–32, jan 1989. ISSN 0028–0836. doi: 10.1038/337129a0. URL http://dx.doi.org/10.1038/337129a0.

Francis C Crick and Christof Koch. What is the function of the claustrum? Philosophical Transactions of the Royal Society of London B: Biological Sciences, 360(1458):1271–1279, 2005.

Sébastien M Crouzet and Simon J Thorpe. Low-level cues and ultra-fast face detection. Frontiers in psychology, 2:342, jan 2011. ISSN 1664–1078. doi: 10.3389/fpsyg.2011.00342.

Yuwei Cui, Chetan Surpur, Subutai Ahmad, and Jeff Hawkins. Continuous online sequence learning with an unsupervised neural network model. dec 2015. URL http://arxiv.org/abs/1512.05463.

Maria C Dadarlat, Joseph E O’Doherty, and Philip N Sabes. A learning-based approach to artificial sensory feedback leads to optimal integration. Nature neuroscience, 18(1): 138–144, 2015.

Ivo Danihelka, Greg Wayne, Benigno Uria, Nal Kalchbrenner, and Alex Graves. Associative Long Short-Term Memory. feb 2016. URL http://arxiv.org/abs/1602.03032.

Nathaniel D Daw, Yael Niv, and Peter Dayan. Actions, policies, values and the basal ganglia. Recent breakthroughs in basal ganglia research, pages 91–106, 2006.

Peter Dayan. Twenty-five lessons from computational neuromodulation. Neuron, 76(1): 240–56, oct 2012. ISSN 1097–4199. doi: 10.1016/j.neuron.2012.09.027. URL http://www.sciencedirect.com/science/article/pii/S0896627312008628.

T Dean. A computational model of the cerebral cortex. Proceedings of the National Conference on Artificial …, 2005. URL http://www.aaai.org/Papers/AAAI/2005/AAAI05–148.pdf.

Stanislas Dehaene, Florent Meyniel, Catherine Wacongne, Liping Wang, and Christophe Pallier. The Neural Representation of Sequences: From Transition Probabilities to Algebraic Patterns and Linguistic Trees. Neuron, 88(1): 2–19, oct 2015. ISSN 08966273. doi: 10.1016/j.neuron.2015.09.019. URL http://www.sciencedirect.com/science/article/pii/S089662731500776X.

Tessa M Dekker and Marko Nardini. Risky visuo-motor choices during rapid reaching in childhood. Developmental science, 2015.

Olivier Delalleau and Yoshua Bengio. Shallow vs. Deep Sum-Product Networks. In Advances in Neural Information Processing Systems, pages 666–674, 2011. URL http://papers.nips.cc/paper/4350-shallow-vs-deep-sum-product-networks.

B DePasquale, MM Churchland, and LF Abbott. Using firing-rate dynamics to train recurrent networks of spiking model neurons. arXiv preprint arXiv: …, 2016. URL http://arxiv.org/abs/1601.07620.

T DeWolf and C Eliasmith. The neural optimal control hierarchy for motor control. Journal of neural engineering, 8(6): 065009, dec 2011. ISSN 1741–2552. doi: 10.1088/1741-2560/8/6/065009. URL http://www.ncbi.nlm.nih.gov/pubmed/22056418.

James J DiCarlo, Davide Zoccolan, and Nicole C Rust. How does the brain solve visual object recognition? Neuron, 73(3): 415–434, 2012.

Rodney J Douglas and Kevan A C Martin. Neuronal circuits of the neocortex. Annual review of neuroscience, 27:419–51, jan 2004. ISSN 0147–006X. doi: 10.1146/annurev.neuro.27.070203.144152. URL http://www.ncbi.nlm.nih.gov/pubmed/15217339.

Kenji Doya. What are the computations of the cerebellum, the basal ganglia and the cerebral cortex? Neural networks, 12(7): 961–974, 1999.

Joshua T Dudman, David Tsay, and Steven A Siegelbaum. A role for synaptic inputs at distal dendrites: instructive signals for hippocampal long-term plasticity. Neuron, 56(5): 866–879, 2007.

Serge O Dumoulin and Brian A Wandell. Population receptive field estimates in human visual cortex. Neuroimage, 39(2): 647–660, 2008.

Chris Eliasmith. How to Build a Brain: A Neural Architecture for Biological Cognition. Oxford University Press, 2013. ISBN 0199794545. URL http://books.google.com/books?id=BK0YRJPmuzgC{&}pgis=1.

Chris Eliasmith and Charles H. Anderson. Neural Engineering: Computation, Representation, and Dynamics in Neurobiological Systems. MIT Press, 2004. ISBN 0262550601. URL http://books.google.com/books?id=J6jz9s4kbfIC{&}pgis=1.

Chris Eliasmith and James Martens. Normalization for probabilistic inference with neurons. Biological cybernetics, 104(4–5):251–262, 2011.

Chris Eliasmith, Terrence C Stewart, Xuan Choo, Trevor Bekolay, Travis DeWolf, Yichuan Tang, Charlie Tang, and Daniel Rasmussen. A large-scale model of the functioning brain. Science (New York, N.Y.), 338(6111): 1202–5, nov 2012. ISSN 1095–9203. doi: 10.1126/science.1225266. URL http://www.sciencemag.org/content/338/6111/1202.

Stephen T Emlen. Migratory orientation in the indigo bunting, passerina cyanea: part i: evidence for use of celestial cues. The Auk, 84(3): 309–342, 1967.

Pierre Enel, Emmanuel Procyk, René Quilodran, and Peter Ford Dominey. Reservoir computing properties of neural dynamics in prefrontal cortex. PLOS Comput Biol, 12(6):e1004967, 2016.

D Erhan and PA Manzagol. The difficulty of training deep architectures and the effect of unsupervised pre-training. International …, 2009. URL http://machinelearning.wustl.edu/mlpapers/paper{_}files/AISTATS09{_}ErhanMBBV.pdf.

SM Eslami, N Heess, and T Weber. Attend, Infer, Repeat: Fast Scene Understanding with Generative Models. arXiv preprint arXiv: …, 2016. URL http://arxiv.org/abs/1603.08575.

Caitlin M Fausey, Swapnaa Jayaraman, and Linda B Smith. From faces to hands: Changing visual input in the first two years. Cognition, 152:101–107, 2016.

Daniel J Felleman and David C Van Essen. Distributed hierarchical processing in the primate cerebral cortex. Cerebral cortex, 1(1): 1–47, 1991.

David Ferster and Kenneth D. Miller. Neural Mechanisms of Orientation Selectivity in the Visual Cortex. nov 2003. URL http://www.annualreviews.org/doi/abs/10.1146/annurev.neuro.23.1.441.

Eberhard E Fetz. Operant conditioning of cortical unit activity. Science, 163(3870): 955–958, 1969.

Eberhard E Fetz. Volitional control of neural activity: implications for brain–computer interfaces. The Journal of physiology, 579(3): 571–579, 2007.

Ila R Fiete and H Sebastian Seung. Gradient learning in spiking neural networks by dynamic perturbation of conductances. Physical review letters, 97(4): 048104, jul 2006. ISSN 0031–9007. doi: 10.1103/PhysRevLett.97.048104. URL http://journals.aps.org/prl/abstract/10.1103/PhysRevLett.97.048104.

Ila R Fiete, Michale S Fee, and H Sebastian Seung. Model of birdsong learning based on gradient estimation by dynamic perturbation of neural conductances. Journal of neurophysiology, 98 (4):2038–57, oct 2007. ISSN 0022–3077. doi: 10.1152/jn.01311.2006. URL http://www.ncbi.nlm.nih.gov/pubmed/17652414.

Ila R Fiete, Walter Senn, Claude Z H Wang, and Richard H R Hahnloser. Spike-time-dependent plasticity and heterosynaptic competition organize networks to produce long scale-free sequences of neural activity. Neuron, 65(4): 563–76, feb 2010. ISSN 1097–4199. doi: 10.1016/j.neuron. 2010.02.003. URL http://www.ncbi.nlm.nih.gov/pubmed/20188660.

Chelsea Finn, Sergey Levine, and Pieter Abbeel. Guided cost learning: Deep inverse optimal control via policy optimization. arXiv preprint arXiv:1603.00448, 2016.

GT Finnerty and MN Shadlen. Time in Cortical Circuits. The Journal of …, 2015. URL https://www.jneurosci.org/content/35/41/13912.full.

Janet Dean Fodor and Carrie Crowther. Understanding stimulus poverty arguments. The Linguistic Review, 18(1–2):105–145, 2002.

Peter Földiák. Learning Invariance from Transformation Sequences, mar 2008. URL http://www.mitpressjournals.org/doi/abs/10.1162/neco.1991.3.2.194{#}.VofznO8rKHo.

D. J. Foster, R. G. M. Morris, Peter Dayan, and Centre For Neuroscience. Models of Hippocampally Dependent Navigation, Using The Temporal Difference Learning Rule. sep 2000.

Julien Fournier, Christian M Müller, and Gilles Laurent. Looking for the roots of cortical sensory computation in three-layered cortices. Current opinion in neurobiology, 31:119–126, 2015.

Steven L Franconeri, Zenon W Pylyshyn, and Brian J Scholl. A simple proximity heuristic allows tracking of multiple objects through occlusion. Attention, Perception, & Psychophysics, 74(4): 691–702, 2012.

Michael J Frank and David Badre. Mechanisms of hierarchical reinforcement learning in corticostriatal circuits 1: computational analysis. Cerebral cortex (New York, N.Y. : 1991), 22(3): 509–26, mar 2012. ISSN 1460–2199. doi: 10.1093/cercor/bhr114.

Steven M Frankland and Joshua D Greene. An architecture for encoding sentence meaning in left mid-superior temporal cortex. Proceedings of the National Academy of Sciences of the United States of America, 112(37): 11732–11737, aug 2015. ISSN 1091–6490. doi: 10.1073/pnas.1421236112. URL http://www.pnas.org/content/112/37/11732.full.

Mathias Franzius, Henning Sprekeler, and Laurenz Wiskott. Slowness and sparseness lead to place, head-direction, and spatial-view cells. PLoS computational biology, 3(8):e166, aug 2007. ISSN 1553–7358. doi: 10.1371/journal.pcbi.0030166. URL http://journals.plos.org/ploscompbiol/article?id=10.1371/journal.pcbi.0030166.

K Friston. The free-energy principle: a unified brain theory? Nature Reviews Neuroscience, 2010. URL http://www.nature.com/nrn/journal/v11/n2/abs/nrn2787.html.

KJ Friston and KE Stephan. Free-energy and the brain. Synthese, 2007. URL http://link.springer.com/article/10.1007/s11229–007–9237-y.

Kunihiko Fukushima. Neocognitron: A self-organizing neural network model for a mechanism of pattern recognition unaffected by shift in position. Biological cybernetics, 36(4): 193–202, 1980.

Mathieu N Galtier and Gilles Wainrib. A biological gradient descent for prediction through a combination of stdp and homeostatic plasticity. Neural computation, 25(11): 2815–2832, 2013.

Tao Gao, Daniel Harari, Joshua Tenenbaum, and Shimon Ullman. When Computer Vision Gazes at Cognition. dec 2014. URL http://arxiv.org/abs/1412.2672.

Ian Gemp and Sridhar Mahadevan. Modeling context in cognition using variational inequalities. 2015.

Dileep George and Jeff Hawkins. Towards a mathematical theory of cortical micro-circuits. PLoS computational biology, 5(10):e1000532, oct 2009. ISSN 1553–7358. doi: 10.1371/journal.pcbi.1000532.

Samuel J Gershman and Jeffrey M Beck. Complex probabilistic inference: From cognition to neural computation. 2016.

Samuel J Gershman, Christopher D Moore, Michael T Todd, Kenneth A Norman, and Per B Sederberg. The successor representation and temporal context. Neural Computation, 24(6): 1553–1568, 2012.

Samuel J Gershman, Ahmed A Moustafa, and Elliot A Ludvig. Time representation in reinforcement learning models of the basal ganglia. 2014.

Iain Murray Zoubin Ghahramani. A note on the evidence and bayesian occams razor. 2005.

Lisa M Giocomo, Eric A Zilli, Erik Fransén, and Michael E Hasselmo. Temporal frequency of sub-threshold oscillations scales with entorhinal grid cell field spacing. Science, 315(5819): 1719–1722, 2007.

Nicolas Giret, Joergen Kornfeld, Surya Ganguli, and Richard H R Hahnloser. Evidence for a causal inverse model in an avian cortico-basal ganglia circuit. Proceedings of the National Academy of Sciences of the United States of America, 111(16): 6063–6068, apr 2014. ISSN 1091–6490. doi: 10.1073/pnas.1317087111. URL http://www.pnas.org/content/111/16/6063.full.

B Goertzel. How might the brain represent complex symbolic knowledge? Neural Networks (IJCNN), 2014 International Joint …, 2014. URL http://ieeexplore.ieee.org/xpls/abs{_}all.jsp?arnumber=6889662.

Mark S Goldman, Joseph H Levine, Guy Major, David W Tank, and HS Seung. Robust persistent neural activity in a model integrator with multiple hysteretic dendrites per neuron. Cerebral cortex, 13(11): 1185–1195, 2003.

S L Gonzalez Andino and R Grave de Peralta Menendez. Coding of saliency by ensemble bursting in the amygdala of primates. Frontiers in behavioral neuroscience, 6:38, jan 2012. ISSN 1662–5153. doi: 10.3389/fnbeh.2012.00038. URL http://journal.frontiersin.org/article/10.3389/fnbeh.2012.00038/abstract.

Cynthia M Gooch, Martin Wiener, Elaine B Wencil, and H Branch Coslett. Interval timing disruptions in subjects with cerebellar lesions. Neuropsychologia, 48(4): 1022–31, mar 2010. ISSN 1873–3514. doi: 10.1016/j.neuropsychologia.2009.11.028.

Ian J. Goodfellow, Jean Pouget-Abadie, Mehdi Mirza, Bing Xu, David Warde-Farley, Sherjil Ozair, Aaron Courville, and Yoshua Bengio. Generative Adversarial Networks. jun 2014a. URL http://arxiv.org/abs/1406.2661.

Ian J. Goodfellow, Oriol Vinyals, and Andrew M. Saxe. Qualitatively characterizing neural network optimization problems. dec 2014b. URL http://arxiv.org/abs/1412.6544.

Alison Gopnik, Andrew N Meltzoff, and Patricia K Kuhl. The scientist in the crib: What early learning tells us about the mind. Harper Paperbacks, 2000.

Alex Graves, Greg Wayne, and Ivo Danihelka. Neural Turing Machines. ArXiv, oct 2014. URL http://arxiv.org/abs/1410.5401.

Ann M Graybiel. The basal ganglia and chunking of action repertoires. Neurobiology of learning and memory, 70(1): 119–136, 1998.

Karol Gregor, Ivo Danihelka, Alex Graves, Danilo Jimenez Rezende, and Daan Wierstra. DRAW: A Recurrent Neural Network For Image Generation. feb 2015. URL http://arxiv.org/abs/1502.04623.

Sten Grillner, Jeanette Hellgren, Ariane Ménard, Kazuya Saitoh, and Martin A Wikström. Mechanisms for selection of basic motor programs–roles for the striatum and pallidum. Trends in neurosciences, 28(7): 364–70, jul 2005. ISSN 0166–2236. doi: 10.1016/j.tins.2005.05.004. URL http://www.ncbi.nlm.nih.gov/pubmed/15935487.

Stephen Grossberg. Adaptive Resonance Theory: how a brain learns to consciously attend, learn, and recognize a changing world. Neural networks : the official journal of the International Neural Network Society, 37:1–47, jan 2013. ISSN 1879–2782. doi: 10.1016/j.neunet.2012.09.017. URL http://www.sciencedirect.com/science/article/pii/S0893608012002584.

Umut Güçlü and Marcel AJ van Gerven. Deep neural networks reveal a gradient in the complexity of neural representations across the ventral stream. The Journal of Neuroscience, 35(27): 10005–10014, 2015.

Arthur Guez, David Silver, and Peter Dayan. Efficient bayes-adaptive reinforcement learning using sample-based search. In Advances in Neural Information Processing Systems, pages 1025–1033, 2012.

Çalar Gülçehre and Yoshua Bengio. Knowledge Matters: Importance of Prior Information for Optimization. Journal of Machine Learning Research, 17(8): 1–32, 2016. URL http://jmlr.org/papers/v17/gulchere16a.html.

Onur Güntürkün and Thomas Bugnyar. Cognition without cortex. Trends in cognitive sciences, 20 (4):291–303, 2016.

K Gurney, T J Prescott, and P Redgrave. A computational model of action selection in the basal ganglia. I. A new functional anatomy. Biological cybernetics, 84(6): 401–10, jun 2001. ISSN 0340–1200. URL http://www.ncbi.nlm.nih.gov/pubmed/11417052.

Robert F Hadley. The problem of rapid variable creation. Neural computation, 21(2): 510–32, mar 2009. ISSN 0899–7667. doi: 10.1162/neco.2008.07-07-572. URL http://www.ncbi.nlm.nih.gov/pubmed/19431268.

J Kiley Hamlin, Karen Wynn, and Paul Bloom. Social evaluation by preverbal infants. Nature, 450 (7169):557–9, nov 2007. ISSN 1476–4687. doi: 10.1038/nature06288. URL http://dx.doi.org/10.1038/nature06288.

Balázs Hangya, SachinP. Ranade, Maja Lorenc, and Adam Kepecs. Central Cholinergic Neurons Are Rapidly Recruited by Reinforcement Feedback. Cell, 162(5): 1155–1168, aug 2015. ISSN 00928674. doi: 10.1016/j.cell.2015.07.057. URL http://www.sciencedirect.com/science/article/pii/S0092867415009733.

A Hanuschkin, S Ganguli, and R H R Hahnloser. A Hebbian learning rule gives rise to mirror neurons and links them to control theoretic inverse models. Frontiers in neural circuits, 7:106, jan 2013. ISSN 1662–5110. doi: 10.3389/fncir.2013.00106.

Christopher M Harris and Daniel M Wolpert. Signal-dependent noise determines motor planning. Nature, 394(6695): 780–784, 1998.

KD Harris. Stability of the fittest: organizing learning through retroaxonal signals. Trends in neurosciences, 2008. URL http://www.sciencedirect.com/science/article/pii/S0166223608000180.

D Hassabis and EA Maguire. The construction system of the brain. … of the Royal …, 2009. URL http://rstb.royalsocietypublishing.org/content/364/1521/1263.short.

Demis Hassabis and Eleanor A Maguire. Deconstructing episodic memory with construction. Trends in cognitive sciences, 11(7): 299–306, jul 2007. ISSN 1364–6613. doi: 10.1016/j.tics.2007.05.001. URL http://www.sciencedirect.com/science/article/pii/S1364661307001258.

Michael E Hasselmo. The role of acetylcholine in learning and memory. Current opinion in neurobiology, 16(6): 710–5, dec 2006. ISSN 0959–4388. doi: 10.1016/j.conb.2006.09.002.

Michael E Hasselmo. If i had a million neurons: Potential tests of cortico-hippocampal theories. Progress in brain research, 219:1–19, 2015.

Michael E Hasselmo and Chantal E Stern. Current questions on space and time encoding. Hippocampus, 25(6): 744–752, 2015.

Michael E Hasselmo and Bradley P Wyble. Free recall and recognition in a network model of the hippocampus: simulating effects of scopolamine on human memory function. Behavioural Brain Research, 89(1–2):1–34, dec 1997. ISSN 01664328. doi: 10.1016/S0166-4328(97)00048-X. URL http://www.sciencedirect.com/science/article/pii/S016643289700048X.

Daisuke Hattori, Ebru Demir, Ho Won Kim, Erika Viragh, S Lawrence Zipursky, and Barry J Dickson. Dscam diversity is essential for neuronal wiring and self-recognition. Nature, 449(7159): 223–227, sep 2007. ISSN 1476–4687. doi: 10.1038/nature06099.

Jeff Hawkins and Subutai Ahmad. Why neurons have thousands of synapses, a theory of sequence memory in neocortex. Frontiers in neural circuits, 10, 2016.

Jeff Hawkins and Sandra Blakeslee. On Intelligence. Henry Holt and Company, 2007. ISBN 1429900458. URL http://books.google.com/books?id=Qg2dmntfxmQC{&}pgis=1.

Akiko Hayashi-Takagi, Sho Yagishita, Mayumi Nakamura, Fukutoshi Shirai, Yi I Wu, Amanda L Loshbaugh, Brian Kuhlman, Klaus M Hahn, and Haruo Kasai. Labelling and optical erasure of synaptic memory traces in the motor cortex. Nature, 2015.

Simon S. Haykin. Neural networks: a comprehensive foundation. Macmillan, 1994. ISBN 0023527617. URL https://books.google.com/books/about/Neural{_}networks.html?id=OZJKAQAAIAAJ{&}pgis=1.

Kenneth J Hayworth. Dynamically partitionable autoassociative networks as a solution to the neural binding problem. Frontiers in computational neuroscience, 6:73, jan 2012. ISSN 1662–5188. doi: 10.3389/fncom.2012.00073.

Kenneth J Hayworth, Mark D Lescroart, and Irving Biederman. Neural encoding of relative position. Journal of experimental psychology. Human perception and performance, 37(4): 1032–50, aug 2011. ISSN 1939–1277. doi: 10.1037/a0022338. URL http://www.ncbi.nlm.nih.gov/pubmed/21517211.

Guillaume Hennequin, Tim P Vogels, and Wulfram Gerstner. Optimal control of transient dynamics in balanced networks supports generation of complex movements. Neuron, 82(6): 1394–1406, 2014.

SA Herd, KA Krueger, and TE Kriete. Strategic cognitive sequencing: a computational cognitive neuroscience approach. Computational …, 2013. URL http://dl.acm.org/citation.cfm?id=2537972.

I. Higgins, L. Matthey, X. Glorot, A. Pal, B. Uria, C. Blundell, S. Mohamed, and A. Lerchner. Early Visual Concept Learning with Unsupervised Deep Learning. ArXiv e-prints, June 2016.

G Hinton. How to do backpropagation in a brain. Invited talk at the NIPS’2007 Deep Learning Workshop, 2007. URL http://www.cs.utoronto.ca/{˜}hinton/backpropincortex2014.pdf.

G Hinton. Can the brain do back-propagation? Invited talk at Stanford University Colloquium on Computer Systems, 2016. URL https://www.youtube.com/watch?v=VIRCybGgHts.

GE Hinton. Connectionist learning procedures. Artificial intelligence, 1989. URL http://www.sciencedirect.com/science/article/pii/0004370289900490.

GE Hinton and JL McClelland. Learning representations by recirculation. Neural information processing …, 1988.

GE Hinton, P Dayan, BJ Frey, and RM Neal. The” wake-sleep” algorithm for unsupervised neural networks. Science, 1995. URL http://www.sciencemag.org/content/268/5214/1158.short.

GE Hinton, A Krizhevsky, and SD Wang. Transforming auto-encoders. Artificial Neural Networks and …, 2011. URL http://link.springer.com/chapter/10.1007/978–3–642–21735–7{_}6.

Geoffrey E Hinton, Simon Osindero, and Yee-Whye Teh. A fast learning algorithm for deep belief nets. Neural computation, 18(7): 1527–54, jul 2006. ISSN 0899–7667. doi: 10.1162/neco.2006.18.7.1527. URL http://www.ncbi.nlm.nih.gov/pubmed/16764513.

Samuel Andrew Hires, Diego A Gutnisky, Jianing Yu, Daniel H O’Connor, and Karel Svoboda. Low-noise encoding of active touch by layer 4 in the somatosensory cortex. eLife, 4, jan 2015. ISSN 2050–084X. doi: 10.7554/eLife.06619.

Mark H Histed, Amy M Ni, and John HR Maunsell. Insights into cortical mechanisms of behavior from microstimulation experiments. Progress in neurobiology, 103:115–130, 2013.

Jonathan Ho and Stefano Ermon. Generative adversarial imitation learning. arXiv preprint arXiv:1606.03476, 2016.

S Hochreiter and J Schmidhuber. Long short-term memory. Neural computation, 9(8): 1735–80, nov 1997. ISSN 0899–7667. URL http://www.ncbi.nlm.nih.gov/pubmed/9377276.

Gregor M Hoerzer, Robert Legenstein, and Wolfgang Maass. Emergence of complex computational structures from chaotic neural networks through reward-modulated Hebbian learning. Cerebral cortex (New York, N.Y. : 1991), 24(3): 677–90, mar 2014. ISSN 1460–2199. doi: 10.1093/cercor/bhs348. URL http://www.ncbi.nlm.nih.gov/pubmed/23146969.

Weizhe Hong and Liqun Luo. Genetic control of wiring specificity in the fly olfactory system. Genetics, 196(1): 17–29, jan 2014. ISSN 1943–2631. doi: 10.1534/genetics.113.154336.

J J Hopfield. Neural networks and physical systems with emergent collective computational abilities. Proceedings of the National Academy of Sciences of the United States of America, 79(8): 2554–8, apr 1982. ISSN 0027–8424.

J J Hopfield. Neurons with graded response have collective computational properties like those of two-state neurons. Proceedings of the National Academy of Sciences of the United States of America, 81(10): 3088–92, may 1984. ISSN 0027–8424. URL http://www.pnas.org/content/81/10/3088.abstract.

J. J. Hopfield. Neurodynamics of mental exploration. Proceedings of the National Academy of Sciences, 107(4): 1648–1653, dec 2009. ISSN 0027–8424. doi: 10.1073/pnas.0913991107. URL http://www.pnas.org/content/107/4/1648.abstract.

Toshihiko Hosoya, Stephen A Baccus, and Markus Meister. Dynamic predictive coding by the retina. Nature, 436(7047): 71–77, 2005.

Y Huang and RPN Rao. Predictive coding. Wiley Interdisciplinary Reviews: Cognitive …, 2011. URL http://onlinelibrary.wiley.com/doi/10.1002/wcs.142/pdf.

Quentin JM Huys, Níall Lally, Paul Faulkner, Neir Eshel, Erich Seifritz, Samuel J Gershman, Peter Dayan, and Jonathan P Roiser. Interplay of approximate planning strategies. Proceedings of the National Academy of Sciences, 112(10): 3098–3103, 2015.

Leyla Isik, Joel Z Leibo, and Tomaso Poggio. Learning and disrupting invariance in visual recognition with a temporal association rule. Frontiers in computational neuroscience, 6:37, jan 2012. ISSN 1662–5188. doi: 10.3389/fncom.2012.00037. URL http://journal.frontiersin.org/article/10.3389/fncom.2012.00037/abstract.

Eugene M Izhikevich. Polychronization: computation with spikes. Neural computation, 18(2): 245–282, 2006.

Eugene M Izhikevich. Solving the distal reward problem through linkage of STDP and dopamine signaling. Cerebral cortex (New York, N.Y. : 1991), 17(10): 2443–52, oct 2007. ISSN 1047–3211. doi: 10.1093/cercor/bhl152. URL http://www.ncbi.nlm.nih.gov/pubmed/17220510.

Gilad A Jacobson and Rainer W Friedrich. Neural circuits: random design of a higher-order olfactory projection. Current biology : CB, 23(10):R448–51, may 2013. ISSN 1879–0445. doi: 10.1016/j.cub.2013.04.016. URL http://www.sciencedirect.com/science/article/pii/S0960982213004247.

M Jaderberg, K Simonyan, and A Zisserman. Spatial transformer networks. Advances in Neural…, 2015a. URL http://papers.nips.cc/paper/5854-spatial-transformer-networks.

M. Jaderberg, W. M. Czarnecki, S. Osindero, O. Vinyals, A. Graves, and K. Kavukcuoglu. Decoupled Neural Interfaces using Synthetic Gradients. ArXiv e-prints, August 2016.

Max Jaderberg, Karen Simonyan, Andrew Zisserman, and Koray Kavukcuoglu. Spatial Transformer Networks. jun 2015b. URL http://arxiv.org/abs/1506.02025.

Herbert Jaeger and Harald Haas. Harnessing nonlinearity: predicting chaotic systems and saving energy in wireless communication. Science (New York, N.Y.), 304(5667): 78–80, apr 2004. ISSN 1095–9203. doi: 10.1126/science.1091277. URL http://www.sciencemag.org/content/304/5667/78.abstract.

Julian Jara-Ettinger, Hyowon Gweon, Laura E Schulz, and Joshua B Tenenbaum. The naïve utility calculus: Computational principles underlying commonsense psychology. Trends in Cognitive Sciences, 20(8): 589–604, 2016.

Santiago Jaramillo and Barak A Pearlmutter. A normative model of attention: Receptive field modulation. Neurocomputing, 58:613–618, 2004.

Hueihan Jhuang, Thomas Serre, Lior Wolf, and Tomaso Poggio. A biologically inspired system for action recognition. In Computer Vision, 2007. ICCV 2007. IEEE 11th International Conference on, pages 1–8. Ieee, 2007.

Daoyun Ji and Matthew A Wilson. Coordinated memory replay in the visual cortex and hippocampus during sleep. Nature neuroscience, 10(1): 100–7, jan 2007. ISSN 1097–6256. doi: 10.1038/nn1825. URL http://dx.doi.org/10.1038/nn1825.

X. Jiang, S. Shen, C. R. Cadwell, P. Berens, F. Sinz, A. S. Ecker, S. Patel, and A. S. Tolias. Principles of connectivity among morphologically defined cell types in adult neocortex. Science, 350(6264):aac9462–aac9462, nov 2015. ISSN 0036–8075. doi: 10.1126/science.aac9462. URL http://science.sciencemag.org/content/350/6264/aac9462.abstract.

Fredrik Johansson, Dan-Anders Jirenhed, Anders Rasmussen, Riccardo Zucca, and Germund Hesslow. Memory trace and timing mechanism localized to cerebellar Purkinje cells. Proceedings of the National Academy of Sciences of the United States of America, 111(41): 14930–4, oct 2014. ISSN 1091–6490. doi: 10.1073/pnas.1415371111. URL http://www.pnas.org/content/111/41/14930.abstract.

Eric Jonas and Konrad Kording. Could a neuroscientist understand a microprocessor? bioRxiv, 2016. doi: 10.1101/055624. URL http://biorxiv.org/content/early/2016/05/26/055624.

Armand Joulin and Tomas Mikolov. Inferring algorithmic patterns with stack-augmented recurrent nets. In Advances in Neural Information Processing Systems, pages 190–198, 2015.

Nir Kalisman, Gilad Silberberg, and Henry Markram. The neocortical microcircuit as a tabula rasa. Proceedings of the National Academy of Sciences of the United States of America, 102(3): 880–5, jan 2005. ISSN 0027–8424. doi: 10.1073/pnas.0407088102.

Nancy Kanwisher, Josh McDermott, and Marvin M Chun. The fusiform face area: a module in human extrastriate cortex specialized for face perception. The Journal of Neuroscience, 17(11): 4302–4311, 1997.

David Kappel, Bernhard Nessler, and Wolfgang Maass. STDP installs in Winner-Take-All circuits an online approximation to hidden Markov model learning. PLoS computational biology, 10 (3):e1003511, mar 2014. ISSN 1553–7358. doi: 10.1371/journal.pcbi.1003511.

David Kappel, Stefan Habenschuss, Robert Legenstein, and Wolfgang Maass. Network plasticity as bayesian inference. PLoS Comput Biol, 11(11): e1004485, 2015.

R Kempter, W Gerstner, and J L van Hemmen. Intrinsic stabilization of output rates by spike-based Hebbian learning. Neural computation, 13 (12):2709–41, dec 2001. ISSN 0899–7667. doi: 10.1162/089976601317098501. URL http://www.ncbi.nlm.nih.gov/pubmed/11705408.

Seyed-Mahdi Khaligh-Razavi and Nikolaus Kriegeskorte. Deep supervised, but not unsupervised, models may explain it cortical representation. PLoS Comput Biol, 10(11): e1003915, 2014.

Diederik P Kingma and Max Welling. Auto-Encoding Variational Bayes. dec 2013. URL http://arxiv.org/abs/1312.6114.

DC Knill and A Pouget. The Bayesian brain: the role of uncertainty in neural coding and computation. TRENDS in Neurosciences, 2004. URL http://www.sciencedirect.com/science/article/pii/S0166223604003352.

Etienne Koechlin and Thomas Jubault. Broca’s area and the hierarchical organization of human behavior. Neuron, 50(6): 963–974, 2006.

Brent Komer and Chris Eliasmith. A unified theoretical approach for biological cognition and learning. Current Opinion in Behavioral Sciences, 11:14–20, mar 2016. ISSN 23521546. doi: 10.1016/j.cobeha.2016.03.006. URL http://www.sciencedirect.com/science/article/pii/S2352154616300651.

K Körding. Decision theory: what” should” the nervous system do? Science, 2007. URL http://science.sciencemag.org/content/318/5850/606.short.

Konrad P Körding, Christoph Kayser, Wolfgang Einhäuser, and Peter König. How are complex cell properties adapted to the statistics of natural stimuli? Journal of neurophysiology, 91(1): 206–12, jan 2004. ISSN 0022–3077. doi: 10.1152/jn.00149. 2003. URL http://jn.physiology.org/content/91/1/206.short.

K.P Körding and P König. A learning rule for dynamic recruitment and decorrelation. Neural Networks, 13(1): 1–9, jan 2000. ISSN 08936080. doi: 10.1016/S0893-6080(99)00088-X. URL http://www.sciencedirect.com/science/article/pii/S089360809900088X.

KP Körding and P König. Supervised and unsupervised learning with two sites of synaptic integration. Journal of Computational Neuroscience, 2001. URL http://link.springer.com/article/10.1023/A:1013776130161.

Minjoon Kouh and Tomaso Poggio. A canonical neural circuit for cortical nonlinear operations. Neural computation, 20(6): 1427–51, jun 2008. ISSN 0899–7667. doi: 10.1162/neco.2008.02-07-466. URL http://www.ncbi.nlm.nih.gov/pubmed/18254695.

Benjamin J Kraus, Robert J Robinson, John A White, Howard Eichenbaum, and Michael E Hasselmo. Hippocampal time cells: time versus path integration. Neuron, 78(6): 1090–1101, 2013.

Nikolaus Kriegeskorte, Marieke Mur, and Peter A Bandettini. Representational similarity analysis-connecting the branches of systems neuroscience. Frontiers in systems neuroscience, 2:4, 2008.

Trenton Kriete, David C Noelle, Jonathan D Cohen, and Randall C O’Reilly. Indirection and symbol-like processing in the prefrontal cortex and basal ganglia. Proceedings of the National Academy of Sciences of the United States of America, 110(41): 16390–16395, oct 2013. ISSN 1091–6490. doi: 10.1073/pnas.1303547110. URL http://www.pnas.org/content/110/41/16390.short.

Ramnandan Krishnamurthy, Aravind S. Lakshminarayanan, Peeyush Kumar, and Balaraman Ravindran. Hierarchical Reinforcement Learning using Spatio-Temporal Abstractions and Deep Neural Networks. page 13, may 2016. URL http://arxiv.org/abs/1605.05359.

Alex Krizhevsky, Ilya Sutskever, and Geoffrey E Hinton. Imagenet classification with deep convolutional neural networks. In Advances in neural information processing systems, pages 1097–1105, 2012.

Tejas D. Kulkarni, Will Whitney, Pushmeet Kohli, and Joshua B. Tenenbaum. Deep Convolutional Inverse Graphics Network. mar 2015. URL http://arxiv.org/abs/1503.03167.

Tejas D. Kulkarni, Karthik R. Narasimhan, Ardavan Saeedi, and Joshua B. Tenenbaum. Hierarchical Deep Reinforcement Learning: Integrating Temporal Abstraction and Intrinsic Motivation. page 13, apr 2016. URL http://arxiv.org/abs/1604.06057.

Dharshan Kumaran, Jennifer J Summerfield, Demis Hassabis, and Eleanor A Maguire. Tracking the emergence of conceptual knowledge during human decision making. Neuron, 63(6): 889–901, sep 2009. ISSN 1097–4199. doi: 10.1016/j.neuron.2009.07.030. URL http://www.cell.com/article/S0896627309006187/fulltext.

Dharshan Kumaran, Demis Hassabis, and James L McClelland. What learning systems do intelligent agents need? complementary learning systems theory updated. Trends in Cognitive Sciences, 20(7): 512–534, 2016.

Karol Kurach, Marcin Andrychowicz, and Ilya Sutskever. Neural Random-Access Machines. page 13, nov 2015. URL http://arxiv.org/abs/1511.06392.

Brenden M. Lake, Ruslan Salakhutdinov, and Joshua B. Tenenbaum. Human-level concept learning through probabilistic program induction. Science, 350(6266): 1332–1338, dec 2015. doi: 10.1126/science.aab3050. URL http://www.sciencemag.org/content/350/6266/1332.full.

Brenden M Lake, Tomer D Ullman, Joshua B Tenenbaum, and Samuel J Gershman. Building machines that learn and think like people. arXiv preprint arXiv:1604.00289, 2016.

Matthew Larkum. A cellular mechanism for cortical associations: an organizing principle for the cerebral cortex. Trends in neurosciences, 36(3): 141–151, mar 2013. ISSN 1878–108X. doi: 10.1016/j.tins.2012.11.006. URL http://www.ncbi.nlm.nih.gov/pubmed/23273272.

Quoc V. Le, Marc’Aurelio Ranzato, Rajat Monga, Matthieu Devin, Kai Chen, Greg S. Corrado, Jeff Dean, and Andrew Y. Ng. Building high-level features using large scale unsupervised learning. dec 2011. URL http://arxiv.org/abs/1112.6209.

Yann LeCun and Yoshua Bengio. Convolutional networks for images, speech, and time series. The handbook of brain theory and neural networks, 3361 (10):1995, 1995.

Yann LeCun, Yoshua Bengio, and Geoffrey Hinton. Deep learning. Nature, 521(7553): 436–444, may 2015. ISSN 0028–0836. doi: 10.1038/nature14539. URL http://dx.doi.org/10.1038/nature14539.

A M Lee, L-H Tai, A Zador, and L Wilbrecht. Between the primate and ‘reptilian’ brain: Rodent models demonstrate the role of corticostriatal circuits in decision making. Neuroscience, 296:66–74, jun 2015. ISSN 1873–7544. doi: 10.1016/j.neuroscience.2014.12.042. URL http://www.sciencedirect.com/science/article/pii/S0306452214010914.

Tai Sing Lee and David Mumford. Hierarchical Bayesian inference in the visual cortex. Journal of the Optical Society of America. A, Optics, image science, and vision, 20(7): 1434–48, jul 2003. ISSN 1084–7529. URL http://www.ncbi.nlm.nih.gov/pubmed/12868647.

TS Lee and AL Yuille. Efficient coding of visual scenes by grouping and segmentation: theoretical predictions and biological evidence. Department of Statistics, UCLA, 2011. URL http://escholarship.org/uc/item/1mc5v1b6.pdf.

Robert Legenstein and Wolfgang Maass. Branch-specific plasticity enables self-organization of nonlinear computation in single neurons. The Journal of neuroscience : the official journal of the Society for Neuroscience, 31(30): 10787–802, jul 2011. ISSN 1529–2401. doi: 10.1523/JNEUROSCI.5684-10.2011. URL http://www.jneurosci.org.libproxy.mit.edu/content/31/30/10787.short.

Joel Z. Leibo, Julien Cornebise, Sergio Gómez, and Demis Hassabis. Approximate Hubel-Wiesel Modules and the Data Structures of Neural Computation. page 13, dec 2015a. URL http://arxiv.org/abs/1512.08457.

Joel Z Leibo, Qianli Liao, Fabio Anselmi, and Tomaso Poggio. The invariance hypothesis implies domain-specific regions in visual cortex. PLoS Comput Biol, 11(10):e1004390, 2015b.

J. Lettvin, H. Maturana, W. McCulloch, and W. Pitts. What the Frog’s Eye Tells the Frog’s Brain. Proceedings of the IRE, 47 (11):1940–1951, nov 1959. ISSN 0096–8390. doi: 10.1109/JRPROC.1959.287207. URL http://ieeexplore.ieee.org/articleDetails.jsp?arnumber=4065609.

Johannes J Letzkus, Björn M Kampa, and Greg J Stuart. Learning rules for spike timing-dependent plasticity depend on dendritic synapse location. The Journal of neuroscience, 26(41): 10420–10429, 2006.

Sergey Levine, Chelsea Finn, Trevor Darrell, and Pieter Abbeel. End-to-end training of deep visuo-motor policies. arXiv preprint arXiv:1504.00702, 2015.

Michael S. Lewicki and Terrence J. Sejnowski. Learning Overcomplete Representations. Neural Computation, 12(2): 337–365, feb 2000. ISSN 0899–7667. doi: 10.1162/089976600300015826.

S. N. Lewis and K. D. Harris. The Neural Marketplace: I. General Formalismand Linear Theory. Technical report, dec 2014. URL http://biorxiv.org/content/early/2014/12/23/013185.abstract.

Nuo Li and James J Dicarlo. Neuronal learning of invariant object representation in the ventral visual stream is not dependent on reward. The Journal of neuroscience : the official journal of the Society for Neuroscience, 32(19): 6611–20, may 2012. ISSN 1529–2401. doi: 10.1523/JNEUROSCI.3786-11.2012. URL http://www.jneurosci.org/content/32/19/6611.full.

Qianli Liao and Tomaso Poggio. Bridging the Gaps Between Residual Learning, Recurrent Neural Networks and Visual Cortex. apr 2016. URL http://arxiv.org/abs/1604.03640.

Qianli Liao, Joel Z. Leibo, and Tomaso Poggio. How Important is Weight Symmetry in Backpropagation? oct 2015. URL http://arxiv.org/abs/1510.05067.

Timothy P. Lillicrap, Daniel Cownden, Douglas B. Tweed, and Colin J. Akerman. Random feedback weights support learning in deep neural networks. page 14, nov 2014. URL http://arxiv.org/abs/1411.0247.

Jian K Liu and Dean V Buonomano. Embedding multiple trajectories in simulated recurrent neural networks in a self-organizing manner. The Journal of neuroscience : the official journal of the Society for Neuroscience, 29(42): 13172–81, oct 2009. ISSN 1529–2401. doi: 10.1523/JNEUROSCI.2358-09.2009. URL http://www.jneurosci.org/content/29/42/13172.short.

Roi Livni, Shai Shalev-Shwartz, and Ohad Shamir. An Algorithm for Training Polynomial Networks. apr 2013. URL http://arxiv.org/abs/1304.7045.

William Lotter, Gabriel Kreiman, and David Cox. Unsupervised Learning of Visual Structure using Predictive Generative Networks. nov 2015. URL http://arxiv.org/abs/1511.06380.

William Lotter, Gabriel Kreiman, and David Cox. Deep predictive coding networks for video prediction and unsupervised learning. arXiv preprint arXiv:1605.08104, 2016.

Steven J Luck and Edward K Vogel. The capacity of visual working memory for features and conjunctions. Nature, 390(6657): 279–281, 1997.

Yan Luo, Xavier Boix, Gemma Roig, Tomaso Poggio, and Qi Zhao. Foveation-based mechanisms alleviate adversarial examples. arXiv preprint arXiv:1511.06292, 2015.

Ashley B Lyons and Erik W Cheries. Inferring social disposition by sound and surface appearance in infancy. Journal of Cognition and Development, (just-accepted), 2016.

Wei Ji Ma, Jeffrey M Beck, Peter E Latham, and Alexandre Pouget. Bayesian inference with probabilistic population codes. Nature neuroscience, 9(11): 1432–8, nov 2006. ISSN 1097–6256. doi: 10.1038/nn1790. URL http://dx.doi.org/10.1038/nn1790.

Wolfgang Maass. Searching for principles of brain computation.

Wolfgang Maass, Thomas Natschläger, and Henry Markram. Real-time computing without stable states: a new framework for neural computation based on perturbations. Neural computation, 14 (11):2531–60, nov 2002. ISSN 0899–7667. doi: 10.1162/089976602760407955.

Wolfgang Maass, Prashant Joshi, and Eduardo D Sontag. Computational aspects of feedback in neural circuits. PLoS computational biology, 3(1):e165, jan 2007. ISSN 1553–7358. doi: 10.1371/journal.pcbi.0020165. URL http://journals.plos.org/ploscompbiol/article?id=10.1371/journal.pcbi.0020165.

Christopher J MacDonald, Kyle Q Lepage, Uri T Eden, and Howard Eichenbaum. Hippocampal time cells bridge the gap in memory for discontiguous events. Neuron, 71(4): 737–749, 2011.

D Maclaurin, D Duvenaud, and RP Adams. Gradient-based hyperparameter optimization through reversible learning. arXiv preprint arXiv:1502.03492, 2015. URL http://arxiv.org/abs/1502.03492.

Joseph G Makin, Matthew R Fellows, and Philip N Sabes. Learning multisensory integration and coordinate transformation via density estimation. PLoS Comput Biol, 9(4):e1003035, 2013.

Joseph G Makin, Benjamin K Dichter, and Philip N Sabes. Recurrent exponential-family harmoniums without backprop-through-time. arXiv preprint arXiv:1605.05799, 2016.

Yael Mandelblat-Cerf, Liora Las, Natalia Denisenko, and Michale S Fee. A role for descending auditory cortical projections in songbird vocal learning. eLife, 3:e02152, jan 2014. ISSN 2050–084X. doi: 10.7554/eLife.02152. URL http://elifesciences.org/content/3/e02152.abstract.

Vikash Mansinghka and Eric Jonas. Building fast bayesian computing machines out of intentionally stochastic, digital parts. arXiv preprint arXiv:1402.4914, 2014.

Adam H Marblestone and Edward S Boyden. Designing tools for assumption-proof brain mapping. Neuron, 83(6): 1239–41, sep 2014. ISSN 1097–4199. doi: 10.1016/j.neuron.2014.09.004. URL http://www.cell.com/article/S0896627314007922/fulltext.

Gary Marcus. The Algebraic Mind: Integrating Connectionism and Cognitive Science. MIT Press, 2001. ISBN 0262632683. URL http://books.google.com/books?id=7YpuRUlFLm8C{&}pgis=1.

Gary Marcus. The birth of the mind: How a tiny number of genes creates the complexities of human thought. Basic Books, 2004.

Gary Marcus, Adam Marblestone, and Thomas Dean. Frequently asked question for: the atoms of neural computation. oct 2014a. URL http://arxiv.org/abs/1410.8826.

Gary Marcus, Adam Marblestone, and Thomas Dean. The atoms of neural computation. Science, 346(6209): 551–552, 2014b. doi: 10.1126/science.1261661.

Eve Marder and Jean-Marc Goaillard. Variability, compensation and homeostasis in neuron and network function. Nature reviews. Neuroscience, 7(7): 563–74, jul 2006. ISSN 1471–003X. doi: 10.1038/nrn1949. URL http://dx.doi.org/10.1038/nrn1949.

David A Markowitz, Clayton E Curtis, and Bijan Pesaran. Multiple component networks support working memory in prefrontal cortex. Proceedings of the National Academy of Sciences of the United States of America, 112(35): 11084–11089, aug 2015. ISSN 1091–6490. doi: 10.1073/pnas.1504172112. URL http://www.pnas.org/content/112/35/11084.abstract.

H Markram, J Lübke, M Frotscher, and B Sakmann. Regulation of synaptic efficacy by coincidence of postsynaptic APs and EPSPs. Science (New York, N.Y.), 275(5297): 213–5, jan 1997. ISSN 0036–8075. URL http://www.ncbi.nlm.nih.gov/pubmed/8985014.

Henry Markram, Eilif Muller, Srikanth Ramaswamy, MichaelW. Reimann, Marwan Abdellah, CarlosAguado Sanchez, Anastasia Ailamaki, Lidia Alonso-Nanclares, Nicolas Antille, Selim Arsever, GuyAntoineAtenekeng Kahou, ThomasK. Berger, Ahmet Bilgili, Nenad Buncic, Athanassia Chalimourda, Giuseppe Chindemi, Jean-Denis Courcol, Fabien Delalondre, Vincent Delattre, Shaul Druckmann, Raphael Dumusc, James Dynes, Stefan Eilemann, Eyal Gal, MichaelEmiel Gevaert, Jean-Pierre Ghobril, Albert Gidon, JoeW. Graham, Anirudh Gupta, Valentin Haenel, Etay Hay, Thomas Heinis, JuanB. Hernando, Michael Hines, Lida Kanari, Daniel Keller, John Kenyon, Georges Khazen, Yihwa Kim, JamesG. King, Zoltan Kisvarday, Pramod Kumbhar, Sébastien Lasserre, Jean-Vincent LeBé, BrunoR.C. Magalhães, Angel Merchán-Pérez, Julie Meystre, BenjaminRoy Morrice, Jeffrey Muller, Alberto Muñoz-Céspedes, Shruti Muralidhar, Keerthan Muthurasa, Daniel Nachbaur, TaylorH. Newton, Max Nolte, Aleksandr Ovcharenko, Juan Palacios, Luis Pastor, Rodrigo Perin, Rajnish Ranjan, Imad Riachi, José-Rodrigo Rodríguez, JuanLuis Riquelme, Christian Rössert, Konstantinos Sfyrakis, Ying Shi, JulianC. Shillcock, Gilad Silberberg, Ricardo Silva, Farhan Tauheed, Martin Telefont, Maria Toledo-Rodriguez, Thomas Tränkler, Werner VanGeit, JafetVillafranca Díaz, Richard Walker, Yun Wang, StefanoM. Zaninetta, Javier DeFelipe, SeanL. Hill, Idan Segev, and Felix Schürmann. Reconstruction and Simulation of Neocortical Microcircuitry. Cell, 163(2): 456–492, oct 2015. ISSN 00928674. doi: 10. 1016/j.cell.2015.09.029. URL http://www.cell.com/article/S0092867415011915/fulltext.

D Marr. A theory of cerebellar cortex. The Journal of physiology, 202(2): 437–70, jun 1969. ISSN 0022–3751.

J Martens and I Sutskever. Learning recurrent neural networks with hessian-free optimization. Proceedings of the 28th …, 2011. URL http://machinelearning.wustl.edu/mlpapers/paper{_}files/ICML2011Martens{_}532.pdf.

Bruce D McCandliss, Laurent Cohen, and Stanislas Dehaene. The visual word form area: expertise for reading in the fusiform gyrus. Trends in cognitive sciences, 7(7): 293–299, 2003.

James L McClelland, David E Rumelhart, et al. Parallel distributed processing, vol. 1. Cambridge, Ma., MIT Press. Zipser D.(1986), Feature Discovery by Competitive Learning, in DE Rumel hart-JL McClelland (eds), voi, 1:151–163, 1986.

Warren S. McCulloch and Walter Pitts. A logical calculus of the ideas immanent in nervous activity. The Bulletin of Mathematical Biophysics, 5 (4):115–133, dec 1943. ISSN 0007–4985. doi: 10.1007/BF02478259. URL http://link.springer.com/10.1007/BF02478259.

Jeffrey L McKinstry, Gerald M Edelman, and Jeffrey L Krichmar. A cerebellar model for predictive motor control tested in a brain-based device. Proceedings of the National Academy of Sciences of the United States of America, 103(9): 3387–3392, feb 2006. ISSN 0027–8424. doi: 10.1073/pnas.0511281103. URL http://www.pnas.org/content/103/9/3387.full{#}ref-4.

Elinor McKone, Kate Crookes, Nancy Kanwisher, et al. The cognitive and neural development of face recognition in humans. The cognitive neurosciences, 4:467–482, 2009.

Peter McLeod and Zoltan Dienes. Do fielders know where to go to catch the ball or only how to get there? Journal of Experimental Psychology: Human Perception and Performance, 22(3): 531, 1996.

BW Mel. The clusteron: toward a simple abstraction for a complex neuron. Advances in neural information processing systems, 1992.

Andrew N Meltzoff. Born to learn: What infants learn from watching us. The role of early experience in infant development, pages 145–164, 1999.

Andrew N Meltzoff, Anna Waismeyer, and Alison Gopnik. Learning about causes from people: observational causal learning in 24-month-old infants. Developmental psychology, 48(5): 1215, 2012.

Andrew N Meltzoff, Rebecca A Williamson, and Peter J Marshall. 11 developmental perspectives on action science: Lessons from infant imitation and cognitive neuroscience. Action science: Foundations of an emerging discipline, page 281, 2013.

George A. Miller. The magical number seven, plus or minus two: some limits on our capacity for processing information. 1956.

K. Miller, J. Keller, and M. Stryker. Ocular dominance column development: analysis and simulation. Science, 245(4918): 605–615, aug 1989. ISSN 0036–8075. doi: 10.1126/science.2762813. URL http://www.sciencemag.org/content/245/4918/605.short.

Kenneth D. Miller and David J. C. MacKay. The Role of Constraints in Hebbian Learning. Neural Computation, 6(1): 100–126, jan 1994. ISSN 0899–7667. doi: 10.1162/neco.1994.6.1.100. URL http://ieeexplore.ieee.org/articleDetails.jsp?arnumber=6797012.

M Minsky. Plain talk about neurodevelopmental epistemology. 1977. URL http://dspace.mit.edu/handle/1721.1/5763.

Marvin Minsky. Society Of Mind. Simon and Schuster, 1988. ISBN 0671657135. URL http://books.google.com/books/about/Society{_}Of{_}Mind.html?id=bLDLllfRpdkC{&}pgis=1.

Marvin Minsky. The emotion machine. New York: Pantheon, 2006.

Marvin L. Minsky. Logical Versus Analogical or Symbolic Versus Connectionist or Neat Versus Scruffy, jun 1991. ISSN 0738–4602. URL http://www.aaai.org/ojs/index.php/aimagazine/article/view/894.

Marvin Lee Minsky and Seymour Papert. Perceptrons: An Introduction to Computational Geometry. Mit Press, 1972. ISBN 0262630222. URL https://books.google.com/books/about/Perceptrons.html?id=Ow1OAQAAIAAJ{&}pgis=1.

Rajiv K. Mishra, Sooyun Kim, Segundo J. Guzman, and Peter Jonas. Symmetric spike timing-dependent plasticity at ca3-ca3 synapses optimizes storage and recall in autoassociative networks. Nat Commun, 7, May 2016. URL http://dx.doi.org/10.1038/ncomms11552.Article.

Tom M Mitchell. The need for biases in learning generalizations. Department of Computer Science, Laboratory for Computer Science Research, Rutgers Univ. New Jersey, 1980.

Shigeru Miyagawa, Robert C Berwick, and Kazuo Okanoya. The emergence of hierarchical structure in human language. Frontiers in psychology, 4:71, jan 2013. ISSN 1664–1078. doi: 10.3389/fpsyg.2013.00071.

Volodymyr Mnih, Nicolas Heess, Alex Graves, et al. Recurrent models of visual attention. In Advances in Neural Information Processing Systems, pages 2204–2212, 2014.

Volodymyr Mnih, Koray Kavukcuoglu, David Silver, Andrei A. Rusu, Joel Veness, Marc G. Bellemare, Alex Graves, Martin Riedmiller, Andreas K. Fidjeland, Georg Ostrovski, Stig Petersen, Charles Beattie, Amir Sadik, Ioannis Antonoglou, Helen King, Dharshan Kumaran, Daan Wierstra, Shane Legg, and Demis Hassabis. Human-level control through deep reinforcement learning. Nature, 518 (7540):529–533, feb 2015. ISSN 0028–0836. doi: 10.1038/nature14236. URL http://dx.doi.org/10.1038/nature14236.

Hossein Mobahi, Ronan Collobert, and Jason Weston. Deep learning from temporal coherence in video. In Proceedings of the 26th Annual International Conference on Machine Learning - ICML ’09, pages 1–8, New York, New York, USA, jun 2009. ACM Press. ISBN 9781605585161. doi: 10.1145/1553374.1553469. URL http://dl.acm.org/citation.cfm?id=1553374.1553469.

Torgeir Moberget, Eva Hilland Gullesen, Stein Andersson, Richard B Ivry, and Tor Endestad. Generalized role for the cerebellum in encoding internal models: evidence from semantic processing. The Journal of neuroscience : the official journal of the Society for Neuroscience, 34(8): 2871–8, feb 2014. ISSN 1529–2401. doi: 10.1523/JNEUROSCI.2264-13.2014. URL http://www.jneurosci.org/content/34/8/2871.short.

Shakir Mohamed and Danilo Jimenez Rezende. Variational Information Maximisation for Intrinsically Motivated Reinforcement Learning. sep 2015. URL http://arxiv.org/abs/1509.08731.

Igor Mordatch, Emanuel Todorov, and Zoran Popović. Discovery of complex behaviors through contact-invariant optimization. ACM Transactions on Graphics (TOG), 31(4): 43, 2012.

Josh Lyskowski Morgan, Daniel Raimund Berger, Arthur Willis Wetzel, and Jeff William Lichtman. The fuzzy logic of network connectivity in mouse visual thalamus. Cell, 165(1): 192–206, 2016.

Micah M Murray, Mark T Wallace, Céline Cappe, Eric M Rouiller, and Pascal Barone. Cortical and thalamic pathways for multisensory and sensori-motor interplay. 2012.

Hajime Mushiake, Naohiro Saito, Kazuhiro Sakamoto, Yasuto Itoyama, and Jun Tanji. Activity in the lateral prefrontal cortex reflects multiple steps of future events in action plans. Neuron, 50(4): 631–41, may 2006. ISSN 0896–6273. doi: 10.1016/j.neuron.2006.03.045. URL http://www.ncbi.nlm.nih.gov/pubmed/16701212.

Marko Nardini, Rachael Bedford, and Denis Mareschal. Fusion of visual cues is not mandatory in children. Proceedings of the National Academy of Sciences, 107(39): 17041–17046, 2010.

Arvind Neelakantan, Quoc V. Le, and Ilya Sutskever. Neural Programmer: Inducing Latent Programs with Gradient Descent. nov 2015. URL http://arxiv.org/abs/1511.04834.

Bernhard Nessler, Michael Pfeiffer, Lars Buesing, and Wolfgang Maass. Bayesian computation emerges in generic cortical microcircuits through spike-timing-dependent plasticity. PLoS computational biology, 9(4):e1003037, apr 2013. ISSN 1553–7358. doi: 10.1371/journal.pcbi.1003037.

AY Ng and SJ Russell. Algorithms for inverse reinforcement learning. Icml, 2000. URL http://ai.stanford.edu/{˜}ang/papers/icml00-irl.pdf.

Mehdi Noroozi and Paolo Favaro. Unsupervised learning of visual representations by solving jigsaw puzzles. arXiv preprint arXiv:1603.09246, 2016.

H Freyja Ólafsdóttir, Caswell Barry, Aman B Saleem, Demis Hassabis, and Hugo J Spiers. Hippocampal place cells construct reward related sequences through unexplored space. eLife, 4: e06063, jun 2015. ISSN 2050–084X. doi: 10.7554/eLife.06063. URL http://elifesciences.org/content/4/e06063.abstract.

Y Ollivier and G Charpiat. Training recurrent networks online without backtracking. arXiv preprint arXiv:1507.07680, 2015. URL http://arxiv.org/abs/1507.07680.

B A Olshausen, C H Anderson, and D C Van Essen. A neurobiological model of visual attention and invariant pattern recognition based on dynamic routing of information. The Journal of neuroscience : the official journal of the Society for Neuroscience, 13(11): 4700–19, nov 1993. ISSN 0270–6474. URL http://www.ncbi.nlm.nih.gov/pubmed/8229193.

Bruno A. Olshausen and David J. Field. Emergence of simple-cell receptive field properties by learning a sparse code for natural images. Nature, 381 (6583):607–609, jun 1996. ISSN 0028–0836. doi: 10.1038/381607a0. URL http://www.ncbi.nlm.nih.gov/pubmed/8637596.

Bruno A. Olshausen and David J. Field. Sparse coding with an overcomplete basis set: A strategy employed by V1? Vision Research, 37(23): 3311–3325, dec 1997. ISSN 00426989. doi: 10.1016/S0042-6989(97)00169-7. URL http://www.sciencedirect.com/science/article/pii/S0042698997001697.

Bruno A Olshausen and David J Field. What is the other 85% of v1 doing. Problems in Systems Neuroscience, 4(5): 182–211, 2004.

Maxime Oquab, Leon Bottou, Ivan Laptev, and Josef Sivic. Learning and transferring mid-level image representations using convolutional neural networks. In Proceedings of the IEEE Conference on Computer Vision and Pattern Recognition, pages 1717–1724, 2014.

Randall C. O’Reilly. Biologically Plausible Error-Driven Learning Using Local Activation Differences: The Generalized Recirculation Algorithm. Neural Computation, 8(5): 895–938, jul 1996. ISSN 0899–7667. doi: 10.1162/neco.1996.8.5.895. URL http://ieeexplore.ieee.org/articleDetails.jsp?arnumber=6796552.

Randall C. O’Reilly. Biologically based computational models of high-level cognition. Science (New York, N.Y.), 314(5796): 91–4, oct 2006. ISSN 1095–9203. doi: 10.1126/science.1127242. URL http://www.sciencemag.org/content/314/5796/91.

Randall C O’Reilly and Michael J Frank. Making working memory work: a computational model of learning in the prefrontal cortex and basal ganglia. Neural computation, 18(2): 283–328, mar 2006. ISSN 0899–7667. doi: 10.1162/089976606775093909. URL http://www.ncbi.nlm.nih.gov/pubmed/16378516.

Randall C. O’Reilly, Thomas E. Hazy, Jessica Mollick, Prescott Mackie, and Seth Herd. Goal-Driven Cognition in the Brain: A Computational Framework. apr 2014a. URL http://arxiv.org/abs/1404.7591.

Randall C. O’Reilly, Dean Wyatte, and John Rohrlich. Learning Through Time in the Thalamocortical Loops. page 37, jul 2014b. URL http://arxiv.org/abs/1407.3432.

RC O’Reilly, Y Munakata, MJ Frank, and TE Hazy. Computational cognitive neuroscience. 2012. URL grey.colorado.edu/mediawiki/sites/CompCogNeuro/images/a/a2/Cecn{_}oreilly{_}intro.pdf.

A. Emin Orhan and Wei Ji Ma. The Inevitability of Probability: Probabilistic Inference in Generic Neural Networks Trained with Non-Probabilistic Feedback. page 26, jan 2016. URL http://arxiv.org/abs/1601.03060.

Lucy M Palmer, Adam S Shai, James E Reeve, Harry L Anderson, Ole Paulsen, and Matthew E Larkum. NMDA spikes enhance action potential generation during sensory input. Nature neuroscience, 17(3): 383–90, mar 2014. ISSN 1546–1726. doi: 10.1038/nn.3646. URL http://dx.doi.org/10.1038/nn.3646.

C Parisien, CH Anderson, and C Eliasmith. Solving the problem of negative synaptic weights in cortical models. Neural computation, 2008. URL http://ieeexplore.ieee.org/xpls/abs{_}all.jsp?arnumber=6796691.

Anitha Pasupathy and Earl K Miller. Different time courses of learning-related activity in the pre-frontal cortex and striatum. Nature, 433(7028): 873–876, feb 2005. ISSN 1476–4687. doi: 10.1038/nature03287. URL http://dx.doi.org/10.1038/nature03287.

AB Patel, T Nguyen, and RG Baraniuk. A Probabilistic Theory of Deep Learning. arXiv preprint arXiv:1504.00641, 2015. URL http://arxiv.org/abs/1504.00641.

Cengiz Pehlevan and Dmitri B Chklovskii. Optimization theory of hebbian/anti-hebbian networks for pca and whitening. In 2015 53rd Annual Allerton Conference on Communication, Control, and Computing (Allerton), pages 1458–1465. IEEE, 2015.

Gertrudis Perea, Marta Navarrete, and Alfonso Araque. Tripartite synapses: astrocytes process and control synaptic information. Trends in neurosciences, 32(8): 421–31, aug 2009. ISSN 1878–108X. doi: 10.1016/j.tins.2009.05.001. URL http://www.ncbi.nlm.nih.gov/pubmed/19615761.

Alexander A. Petrov, David J. Jilk, and Randall C. O’Reilly. The Leabra architecture: Specialization without modularity. Behavioral and Brain Sciences, 33(04): 286–287, oct 2010. ISSN 0140–525X. doi: 10.1017/S0140525X10001160. URL http://journals.cambridge.org/abstract{_}S0140525X10001160.

Giovanni Pezzulo, Paul F M J Verschure, Christian Balkenius, and Cyriel M A Pennartz. The principles of goal-directed decision-making: from neural mechanisms to computation and robotics. Philosophical transactions of the Royal Society of London. Series B, Biological sciences, 369(1655):20130470–, nov 2014. ISSN 1471–2970. doi: 10.1098/rstb.2013.0470. URL http://rstb.royalsocietypublishing.org/content/369/1655/20130470.short.

Jean-Pascal Pfister and Wulfram Gerstner. Triplets of spikes in a model of spike timing-dependent plasticity. The Journal of neuroscience, 26(38): 9673–9682, 2006.

Ann T Phillips, Henry M Wellman, and Elizabeth S Spelke. Infants’ ability to connect gaze and emotional expression to intentional action. Cognition, 85(1): 53–78, 2002.

Filip Piekniewski, Patryk Laurent, Csaba Petre, Micah Richert, Dimitry Fisher, and Todd Hylton. Unsupervised learning from continuous video in a scalable predictive recurrent network. arXiv preprint arXiv:1607.06854, 2016.

Fernando J Pineda. Generalization of back-propagation to recurrent neural networks. Physical review letters, 59(19): 2229, 1987.

S Pinker. How the mind works. Annals of the New York Academy of Sciences, 1999. URL http://onlinelibrary.wiley.com/doi/10.1111/j.1749–6632.1999.tb08538.x/full.

T A Plate. Holographic reduced representations. IEEE transactions on neural networks /a publication of the IEEE Neural Networks Council, 6(3): 623–641, jan 1995. ISSN 1045–9227. doi: 10.1109/72.377968. URL http://ieeexplore.ieee.org/articleDetails.jsp?arnumber=377968.

Tomaso Poggio. What if… 2015. URL http://cbmm.mit.edu/views-reviews/article/what-if.

Tomaso Poggio and Emilio Bizzi. Generalization in vision and motor control. Nature, 431(7010): 768–774, 2004.

Filip Ponulak and John J Hopfield. Rapid, parallel path planning by propagating wavefronts of spiking neural activity. Frontiers in computational neuroscience, 7:98, jan 2013. ISSN 1662–5188. doi: 10.3389/fncom.2013.00098.

Alec Radford, Luke Metz, and Soumith Chintala. Unsupervised Representation Learning with Deep Convolutional Generative Adversarial Networks. nov 2015. URL http://arxiv.org/abs/1511.06434.

Kanaka Rajan, Christopher D Harvey, and David W Tank. Recurrent network models of sequence generation and memory. Neuron, 90(1): 128–142, 2016.

Vilayanur S Ramachandran. Mirror neurons and imitation learning as the driving force behind the great leap forward in human evolution, 2000.

Rajesh PN Rao. Bayesian computation in recurrent neural circuits. Neural computation, 16(1): 1–38, 2004.

Nicolas Rashevsky. Mathematical biophysics: physico-mathematical foundations of biology. Bull. Amer. Math. Soc. 45 (1939), 223–224 DOI: http://dx.doi.org/10.1090/S0002–9904–1939–06963–2 PII, pages 0002–9904, 1939.

A Rasmus and M Berglund. Semi-Supervised Learning with Ladder Networks. Advances in Neural …, 2015. URL papers.nips.cc/paper/5947-semi-supervised-learning-with-ladder-networks.

John H. Reynolds and Robert Desimone. The Role of Neural Mechanisms of Attention in Solving the Binding Problem. Neuron, 24(1): 19–29, sep 1999. ISSN 08966273. doi: 10.1016/S0896-6273(00)80819-3. URL http://www.sciencedirect.com/science/article/pii/S0896627300808193.

Danilo Jimenez Rezende, Shakir Mohamed, Ivo Danihelka, Karol Gregor, and Daan Wierstra. One-Shot Generalization in Deep Generative Models. mar 2016. URL http://arxiv.org/abs/1603.05106.

Mattia Rigotti, Omri Barak, Melissa R Warden, Xiao-Jing Wang, Nathaniel D Daw, Earl K Miller, and Stefano Fusi. The importance of mixed selectivity in complex cognitive tasks. Nature, 497 (7451):585–90, may 2013. ISSN 1476–4687. doi: 10.1038/nature12160.

DA Robinson. Implications of neural networks for how we think about brain function. Behavioral and brain sciences, 1992. URL http://journals.cambridge.org/abstract{_}S0140525X00072563.

A Rodriguez and R Granger. The grammar of mammalian brain capacity. Theoretical Computer Science, 633:100–111, 2016.

A Rodriguez, J Whitson, and R Granger. Derivation and analysis of basic computational operations of thalamocortical circuits. Journal of cognitive neuroscience, 16(5): 856–877, 2004.

Pieter R Roelfsema and Arjen van Ooyen. Attention-gated reinforcement learning of internal representations for classification. Neural computation, 17 (10):2176–214, oct 2005. ISSN 0899–7667. doi: 10.1162/0899766054615699. URL http://www.ncbi.nlm.nih.gov/pubmed/16105222.

PR Roelfsema, A van Ooyen, and T Watanabe. Perceptual learning rules based on reinforcers and attention. Trends in cognitive sciences, 2010. URL http://www.sciencedirect.com/science/article/pii/S1364661309002617.

Edmund T Rolls. The mechanisms for pattern completion and pattern separation in the hippocampus. Frontiers in systems neuroscience, 7:74, jan 2013. ISSN 1662–5137. doi: 10.3389/fnsys.2013.00074. URL http://journal.frontiersin.org/article/10.3389/fnsys.2013.00074/abstract.

Jaldert O Rombouts, Sander M Bohte, and Pieter R Roelfsema. How attention can create synaptic tags for the learning of working memories in sequential tasks. PLoS computational biology, 11(3):e1004060, mar 2015. ISSN 1553–7358. doi: 10.1371/journal.pcbi.1004060. URL http://journals.plos.org/ploscompbiol/article?id=10.1371/journal.pcbi.1004060.

Adriana Romero, Nicolas Ballas, Samira Ebrahimi Kahou, Antoine Chassang, Carlo Gatta, and Yoshua Bengio. Fitnets: Hints for thin deep nets. arXiv preprint arXiv:1412.6550, 2014.

Y Roudi and G Taylor. Learning with hidden variables. Current opinion in neurobiology, 2015. URL http://www.sciencedirect.com/science/article/pii/S0959438815001245.

Christopher J Rozell, Don H Johnson, Richard G Baraniuk, and Bruno A Olshausen. Sparse coding via thresholding and local competition in neural circuits. Neural computation, 20(10): 2526–63, oct 2008. ISSN 0899–7667. doi: 10.1162/neco.2008.03-07-486. URL http://www.ncbi.nlm.nih.gov/pubmed/18439138.

Alon Rubin, Nitzan Geva, Liron Sheintuch, and Yaniv Ziv. Hippocampal ensemble dynamics timestamp events in long-term memory. eLife, page e12247, 2015.

David E. Rumelhart, Geoffrey E. Hinton, and Ronald J. Williams. Learning representations by back-propagating errors. Nature, 323(6088): 533–536, oct 1986. ISSN 0028–0836. doi: 10.1038/323533a0. URL http://www.nature.com/nature/journal/v323/n6088/pdf/323533a0.pdf.

Patrick T Sadtler, Kristin M Quick, Matthew D Golub, Steven M Chase, Stephen I Ryu, Elizabeth C Tyler-Kabara, Byron M Yu, and Aaron P Batista. Neural constraints on learning. Nature, 512(7515): 423–6, aug 2014. ISSN 1476–4687. doi: 10.1038/nature13665. URL http://dx.doi.org/10.1038/nature13665.

Maneesh Sahani and Peter Dayan. Doubly distributional population codes: simultaneous representation of uncertainty and multiplicity. Neural Computation, 15(10): 2255–2279, 2003.

Maya Sandler, Yoav Shulman, and Jackie Schiller. A novel form of local plasticity in tuft dendrites of neocortical somatosensory layer 5 pyramidal neurons. Neuron, 2016.

Adam Santoro, Sergey Bartunov, Matthew Botvinick, Daan Wierstra, and Timothy Lillicrap. One-shot Learning with Memory-Augmented Neural Networks. page 13, may 2016. URL http://arxiv.org/abs/1605.06065.

Andrew M. Saxe, James L. McClelland, and Surya Ganguli. Exact solutions to the nonlinear dynamics of learning in deep linear neural networks. dec 2013. URL http://arxiv.org/abs/1312.6120.

Benjamin Scellier and Yoshua Bengio. Towards a Biologically Plausible Backprop. feb 2016. URL http://arxiv.org/abs/1602.05179.

Mathieu Schiess, Robert Urbanczik, and Walter Senn. Somato-dendritic synaptic plasticity and error-backpropagation in active dendrites. PLoS Comput Biol, 12(2):e1004638, 2016.

J Schmidhuber. Formal theory of creativity, fun, and intrinsic motivation (19902010). Autonomous Mental Development, IEEE …, 2010. URL http://ieeexplore.ieee.org/xpls/abs{_}all.jsp?arnumber=5508364.

Jürgen Schmidhuber. Deep learning in neural networks: An overview. Neural Networks, 61:85–117, 2015.

Brian J Scholl. Can infants’ object concepts be trained? Trends in Cognitive Sciences, 8(2): 49–51, 2004.

Lars Schwabe, Klaus Obermayer, Alessandra Angelucci, and Paul C Bressloff. The role of feedback in shaping the extra-classical receptive field of cortical neurons: a recurrent network model. The Journal of Neuroscience, 26(36): 9117–9129, 2006.

TJ Sejnowski and H Poizner. Prospective Optimization. Proceedings of the …, 2014. URL http://ieeexplore.ieee.org/xpls/abs{_}all.jsp?arnumber=6803897.

P Sermanet and K Kavukcuoglu. Pedestrian detection with unsupervised multi-stage feature learning. Proceedings of the …, 2013.

T. Serre, A. Oliva, and T. Poggio. A feedforward architecture accounts for rapid categorization. Proceedings of the National Academy of Sciences, 104(15): 6424–6429, apr 2007. ISSN 0027–8424. doi: 10.1073/pnas.0700622104. URL http://www.pnas.org/content/104/15/6424.long.

E Servan-Schreiber and JR Anderson. Chunking as a mechanism of implicit learning. Journal of Experimental Psychology: Learning, …, 1990.

H. Sebastian Seung. Continuous attractors and oculomotor control. Neural Networks, 11 (7–8):1253–1258, oct 1998. ISSN 08936080. doi: 10.1016/S0893-6080(98)00064-1. URL http://www.sciencedirect.com/science/article/pii/S0893608098000641.

H. Sebastian Seung. Learning in Spiking Neural Networks by Reinforcement of Stochastic Synaptic Transmission. Neuron, 40(6): 1063–1073, dec 2003. ISSN 08966273. doi: 10.1016/S0896-6273(03)00761-X. URL http://www.sciencedirect.com/science/article/pii/S089662730300761X.

Adam S Shai, Costas A Anastassiou, Matthew E Larkum, and Christof Koch. Physiology of layer 5 pyramidal neurons in mouse primary visual cortex: coincidence detection through bursting. PLoS Comput Biol, 11(3):e1004090, 2015.

Gordon M Shepherd. The microcircuit concept applied to cortical evolution: from three-layer to six-layer cortex. Adaptive Function and Brain Evolution, page 151, 2014.

S Murray Sherman. Thalamic relays and cortical functioning. Progress in brain research, 149:107–26, jan 2005. ISSN 1875–7855. doi: 10.1016/S0079-6123(05)49009-3. URL http://www.ncbi.nlm.nih.gov/pubmed/16226580.

S Murray Sherman. The thalamus is more than just a relay. Current opinion in neurobiology, 17(4): 417–22, aug 2007. ISSN 0959–4388. doi: 10.1016/j.conb.2007.07.003.

Toru Shimizu and Harvey J Karten. Multiple origins of neocortex: Contributions of the dorsal. The Neocortex: Ontogeny and Phylogeny, 200:75, 2013.

Jennie Si. Handbook of learning and approximate dynamic programming, volume 2. John Wiley & Sons, 2004.

Markus Siegel, Melissa R Warden, and Earl K Miller. Phase-dependent neuronal coding of objects in short-term memory. Proceedings of the National Academy of Sciences of the United States of America, 106(50): 21341–6, dec 2009. ISSN 1091–6490. doi: 10.1073/pnas.0908193106. URL http://www.pnas.org/content/106/50/21341.abstract.

Ray Singh and Chris Eliasmith. Higher-dimensional neurons explain the tuning and dynamics of working memory cells. The Journal of neuroscience : the official journal of the Society for Neuroscience, 26(14): 3667–78, apr 2006. ISSN 1529–2401. doi: 10.1523/JNEUROSCI.4864-05.2006. URL http://www.ncbi.nlm.nih.gov/pubmed/16597721.

Jesper Sjöström and Wulfram Gerstner. Spike-timing dependent plasticity. Scholarpedia, 5(2): 1362, feb 2010. ISSN 1941–6016. doi: 10.4249/scholarpedia.1362. URL http://www.scholarpedia.org/article/Spike-timing{_}dependent{_}plasticity.

Per Jesper Sjöström and Michael Häusser. A cooperative switch determines the sign of synaptic plasticity in distal dendrites of neocortical pyramidal neurons. Neuron, 51(2): 227–238, 2006.

Amy E Skerry and Elizabeth S Spelke. Preverbal infants identify emotional reactions that are incongruent with goal outcomes. Cognition, 130(2): 204–216, feb 2014. ISSN 1873–7838. doi: 10.1016/j.cognition.2013.11.002.

WR Softky and C Koch. The highly irregular firing of cortical cells is inconsistent with temporal integration of random EPSPs. The Journal of Neuroscience, 1993. URL http://www.jneurosci.org/content/13/1/334.short.

Soren Van Hout Solari and Rich Stoner. Cognitive consilience: primate non-primary neuroanatomical circuits underlying cognition. Frontiers in neuroanatomy, 5:65, jan 2011. ISSN 1662–5129. doi: 10.3389/fnana.2011.00065. URL http://journal.frontiersin.org/article/10.3389/fnana.2011.00065/abstract.

Soren Van Hout Solari and Rich Stoner. Cognitive consilience: primate non-primary neuroanatomical circuits underlying cognition. Wiring Principles of Cerebral Cortex, page 149, 2015.

Pavel Sountsov and Paul Miller. Spiking neuron network Helmholtz machine. Frontiers in computational neuroscience, 9:46, jan 2015. ISSN 1662–5188. doi: 10.3389/fncom.2015.00046. URL http://journal.frontiersin.org/article/10.3389/fncom.2015.00046/abstract.

Larry R Squire. Memory systems of the brain: a brief history and current perspective. Neurobiology of learning and memory, 82(3): 171–177, 2004.

Nitish Srivastava, Geoffrey Hinton, Alex Krizhevsky, Ilya Sutskever, and Ruslan Salakhutdinov. Dropout: a simple way to prevent neural networks from overfitting. The Journal of Machine Learning Research, 15(1): 1929–1958, jan 2014. ISSN 1532–4435. URL http://dl.acm.org/citation.cfm?id=2627435.2670313.

KL Stachenfeld. Design Principles of the Hippocampal Cognitive Map. Advances in Neural …, 2014.

Terry Stewart and Chris Eliasmith. Compositionality and biologically plausible models. 2009. URL http://philpapers.org/rec/STECAB-2.

Andrea Stocco, Christian Lebiere, and John R Anderson. Conditional routing of information to the cortex: a model of the basal ganglia’s role in cognitive coordination. Psychological review, 117(2): 541–74, apr 2010. ISSN 1939–1471. doi: 10.1037/a0019077.

D. G. Stork. Is backpropagation biologically plausible? In International Joint Conference on Neural Networks, pages 241–246 vol.2. IEEE, 1989. doi: 10.1109/IJCNN.1989.118705. URL http://ieeexplore.ieee.org/articleDetails.jsp?arnumber=118705.

Nicholas J Strausfeld and Frank Hirth. Deep homology of arthropod central complex and vertebrate basal ganglia. Science (New York, N.Y.), 340(6129): 157–61, apr 2013. ISSN 1095–9203. doi: 10.1126/science.1231828. URL http://www.sciencemag.org/content/340/6129/157.short.

Liviu Stnior, Chris van der Togt, Cyriel M A Pennartz, and Pieter R Roelfsema. A unified selection signal for attention and reward in primary visual cortex. Proceedings of the National Academy of Sciences of the United States of America, 110 (22):9136–41, may 2013. ISSN 1091–6490. doi: 10.1073/pnas.1300117110.

Sainbayar Sukhbaatar, Joan Bruna, Manohar Paluri, Lubomir Bourdev, and Rob Fergus. Training convolutional networks with noisy labels. arXiv preprint arXiv:1406.2080, 2014.

Yi Sun, Faustino Gomez, and Jürgen Schmidhuber. Planning to be surprised: Optimal bayesian exploration in dynamic environments. In Artificial General Intelligence, pages 41–51. Springer, 2011.

D Sussillo. Neural circuits as computational dynamical systems. Current opinion in neurobiology, 2014. URL http://www.sciencedirect.com/science/article/pii/S0959438814000166.

D Sussillo and LF Abbott. Generating coherent patterns of activity from chaotic neural networks. Neuron, 2009. URL http://www.sciencedirect.com/science/article/pii/S0896627309005479.

David Sussillo, Mark M Churchland, Matthew T Kaufman, and Krishna V Shenoy. A neural network that finds a naturalistic solution for the production of muscle activity. Nature neuroscience, 18(7): 1025–33, jul 2015. ISSN 1546–1726. doi: 10.1038/nn.4042. URL http://dx.doi.org/10.1038/nn.4042.

I Sutskever and J Martens. On the importance of initialization and momentum in deep learning. Proceedings of the …, 2013. URL http://machinelearning.wustl.edu/mlpapers/papers/icml2013{_}sutskever13.

Ilya Sutskever, James Martens, and Geoffrey E Hinton. Generating text with recurrent neural networks. In Proceedings of the 28th International Conference on Machine Learning (ICML-11), pages 1017–1024, 2011.

Richard S Sutton and Andrew G Barto. Reinforcement learning: An introduction. MIT press, 1998.

Andrea Tacchetti, Leyla Isik, and Tomaso Poggio. Spatio-temporal convolutional neural networks explain human neural representations of action recognition. arXiv preprint arXiv:1606.04698, 2016.

Norio Takata, Tsuneko Mishima, Chihiro Hisatsune, Terumi Nagai, Etsuko Ebisui, Katsuhiko Mikoshiba, and Hajime Hirase. Astrocyte calcium signaling transforms cholinergic modulation to cortical plasticity in vivo. The Journal of neuroscience : the official journal of the Society for Neuroscience, 31(49): 18155–65, dec 2011. ISSN 1529–2401. doi: 10.1523/JNEUROSCI.5289-11.2011. URL http://www.ncbi.nlm.nih.gov/pubmed/22159127.

Aviv Tamar, Sergey Levine, and Pieter Abbeel. Value iteration networks. arXiv preprint arXiv:1602.02867, 2016.

Y Tang, R Salakhutdinov, and G Hinton. Tensor analyzers. Proceedings of The 30th International …, 2013. URL http://jmlr.org/proceedings/papers/v28/tang13.html.

Yichuan Tang, Ruslan Salakhutdinov, and Geoffrey Hinton. Deep Mixtures of Factor Analysers. jun 2012. URL http://arxiv.org/abs/1206.4635.

Jonathan Tapson and André van Schaik. Learning the pseudoinverse solution to network weights. Neural Networks, 45:94–100, 2013.

Rita Morais Tavares, Avi Mendelsohn, Yael Grossman, Christian Hamilton Williams, Matthew Shapiro, Yaacov Trope, and Daniela Schiller. A map for social navigation in the human brain. Neuron, 87(1): 231–243, 2015.

Scott V Taylor and Aldo A Faisal. Does the cost function of human motor control depend on the internal metabolic state? BMC Neuroscience, 12(Suppl 1):P99, 2011. ISSN 1471–2202. doi: 10.1186/1471-2202-12-S1-P99. URL http://www.ncbi.nlm.nih.gov/pmc/articles/PMC3240571/.

Chris Eliasmith Terrence Stewart, Xuan Choo. Symbolic reasoning in spiking neurons: A model of the cortex/basal ganglia/thalamus loop. 32nd Annual Meeting of the Cognitive Science Society, 2010.

D Gowanlock R Tervo, Joshua B Tenenbaum, and Samuel J Gershman. Toward the neural implementation of structure learning. Current opinion in neurobiology, 37:99–105, feb 2016. ISSN 1873–6882. doi: 10.1016/j.conb.2016.01.014. URL http://www.ncbi.nlm.nih.gov/pubmed/26874471.

G Tesauro. Temporal difference learning and TDGammon. Communications of the ACM, 1995.

Dominik Thalmeier, Marvin Uhlmann, Hilbert J. Kappen, and Raoul-Martin Memmesheimer. Learning universal computations with spikes. page 24, may 2015. URL http://arxiv.org/abs/1505.07866.

N Tinbergen. Behavior and natural selection. 1965.

Emanuel Todorov. Cosine tuning minimizes motor errors. Neural Computation, 14(6): 1233–1260, 2002.

Emanuel Todorov. Efficient computation of optimal actions. Proceedings of the national academy of sciences, 106(28): 11478–11483, 2009.

Emanuel Todorov and Michael I Jordan. Optimal feedback control as a theory of motor coordination. Nature neuroscience, 5(11): 1226–1235, 2002.

Bryan Tripp and Chris Eliasmith. Function approximation in inhibitory networks. Neural networks : the official journal of the International Neural Network Society, 77:95–106, may 2016. ISSN 1879–2782. doi: 10.1016/j.neunet.2016.01.010. URL http://www.sciencedirect.com/science/article/pii/S0893608016000113.

Robert S Turner and Michel Desmurget. Basal ganglia contributions to motor control: a vigorous tutor. Current opinion in neurobiology, 20(6): 704–716, 2010.

Gina Turrigiano. Homeostatic synaptic plasticity: local and global mechanisms for stabilizing neuronal function. Cold Spring Harbor perspectives in biology, 4(1):a005736, jan 2012. ISSN 1943–0264. doi: 10.1101/cshperspect.a005736.

Shimon Ullman, Daniel Harari, and Nimrod Dorfman. From simple innate biases to complex visual concepts. Proceedings of the National Academy of Sciences of the United States of America, 109(44): 18215–18220, oct 2012. ISSN 1091–6490. doi: 10.1073/pnas.1207690109. URL http://www.pnas.org/content/109/44/18215.full.

Robert Urbanczik and Walter Senn. Learning by the dendritic prediction of somatic spiking. Neuron, 81(3): 521–8, feb 2014. ISSN 1097–4199. doi: 10.1016/j.neuron.2013.11.030. URL http://www.ncbi.nlm.nih.gov/pubmed/24507189.

H Valpola. From neural PCA to deep unsupervised learning. Adv. in Independent Component Analysis and …, 2015.

Aaron van den Oord, Nal Kalchbrenner, and Koray Kavukcuoglu. Pixel Recurrent Neural Networks. jan 2016. URL http://arxiv.org/abs/1601.06759.

Caroline AA Van Heijningen, Jos De Visser, Willem Zuidema, and Carel Ten Cate. Simple rules can explain discrimination of putative recursive syntactic structures by a songbird species. Proceedings of the National Academy of Sciences, 106(48): 20538–20543, 2009.

Timo Van Kerkoerle, Matthew W Self, Bruno Dagnino, Marie-Alice Gariel-Mathis, Jasper Poort, Chris Van Der Togt, and Pieter R Roelfsema. Alpha and gamma oscillations characterize feedback and feedforward processing in monkey visual cortex. Proceedings of the National Academy of Sciences, 111(40): 14332–14341, 2014.

Andreas Veit, Michael Wilber, and Serge Belongie. Residual Networks are Exponential Ensembles of Relatively Shallow Networks. may 2016. URL http://arxiv.org/abs/1605.06431.

C Verney, M Baulac, B Berger, C Alvarez, A Vigny, and KB Helle. Morphological evidence for a dopaminergic terminal field in the hippocampal formation of young and adult rat. Neuroscience, 14(4): 1039–1052, 1985.

Willem B Verwey. Buffer loading and chunking in sequential keypressing. Journal of Experimental Psychology: Human Perception and Performance, 22(3): 544, 1996.

Jane X Wang, Neal J Cohen, and Joel L Voss. Covert rapid action-memory simulation (CRAMS): a hypothesis of hippocampal-prefrontal interactions for adaptive behavior. Neurobiology of learning and memory, 117:22–33, jan 2015. ISSN 1095–9564. doi: 10.1016/j.nlm.2014.04.003.

Jianyu Wang and Alan Yuille. Semantic Part Segmentation using Compositional Model combining Shape and Appearance. dec 2014. URL http://arxiv.org/abs/1412.6124.

Xiao-Jing Wang. The Prefrontal Cortex as a Quintessential Cognitive-Type Neural Circuit : Principles of Frontal Lobe Function - oi, 2012. URL http://oxfordindex.oup.com/view/10.1093/med/9780199837755.003.0018.

Melissa R Warden and Earl K Miller. The representation of multiple objects in prefrontal neuronal delay activity. Cerebral cortex (New York, N.Y. : 1991), 17 Suppl 1:i41–50, sep 2007. ISSN 1047–3211. doi: 10.1093/cercor/bhm070. URL http://www.ncbi.nlm.nih.gov/pubmed/17726003.

Melissa R Warden and Earl K Miller. Task-dependent changes in short-term memory in the prefrontal cortex. The Journal of neuroscience : the official journal of the Society for Neuroscience, 30(47): 15801–10, nov 2010. ISSN 1529–2401. doi: 10.1523/JNEUROSCI.1569-10. 2010. URL http://www.jneurosci.org/content/30/47/15801.short.

Manuel Watter, Jost Springenberg, Joschka Boedecker, and Martin Riedmiller. Embed to control: A locally linear latent dynamics model for control from raw images. In Advances in Neural Information Processing Systems, pages 2728–2736, 2015.

Greg Wayne and L F Abbott. Hierarchical control using networks trained with higher-level forward models. Neural computation, 26(10): 2163–93, oct 2014. ISSN 1530–888X.

Paul Werbos. Beyond regression: New tools for prediction and analysis in the behavioral sciences. 1974.

PJ Werbos. Applications of advances in nonlinear sensitivity analysis. System modeling and optimization, 1982. URL http://link.springer.com/chapter/10.1007/BFb0006203.

PJ Werbos. Backpropagation through time: what it does and how to do it. Proceedings of the IEEE, 1990. URL http://ieeexplore.ieee.org/xpls/abs{_}all.jsp?arnumber=58337.

Justin Werfel, Xiaohui Xie, and H Sebastian Seung. Learning curves for stochastic gradient descent in linear feedforward networks. Neural computation, 17(12): 2699–718, dec 2005. ISSN 0899–7667. doi: 10.1162/089976605774320539. URL http://ieeexplore.ieee.org/articleDetails.jsp?arnumber=6790355.

Jason Weston, Sumit Chopra, and Antoine Bordes. Memory Networks. oct 2014. URL http://arxiv.org/abs/1410.3916.

William F. Whitney, Michael Chang, Tejas Kulkarni, and Joshua B. Tenenbaum. Understanding Visual Concepts with Continuation Learning. feb 2016. URL http://arxiv.org/abs/1602.06822.

Ronald J. Williams. Simple statistical gradient-following algorithms for connectionist reinforcement learning. Machine Learning, 8(3–4):229–256, may 1992. ISSN 0885–6125. doi: 10.1007/BF00992696. URL http://link.springer.com/10.1007/BF00992696.

Ronald J Williams and Leemon C Baird. Tight performance bounds on greedy policies based on imperfect value functions. Technical report, Citeseer, 1993.

S R Williams and G J Stuart. Backpropagation of physiological spike trains in neocortical pyramidal neurons: implications for temporal coding in dendrites. The Journal of neuroscience : the official journal of the Society for Neuroscience, 20(22): 8238–8246, nov 2000. ISSN 1529–2401. URL http://www.ncbi.nlm.nih.gov/pubmed/11069929.

R I Wilson and R A Nicoll. Endogenous cannabinoids mediate retrograde signalling at hippocampal synapses. Nature, 410(6828): 588–92, mar 2001. ISSN 0028–0836. doi: 10.1038/35069076. URL http://www.ncbi.nlm.nih.gov/pubmed/11279497.

Patrick Henry Winston. The strong story hypothesis and the directed perception hypothesis. 2011.

Laurenz Wiskott and Terrence J Sejnowski. Slow feature analysis: unsupervised learning of invariances. Neural computation, 14(4): 715–70, apr 2002. ISSN 0899–7667. doi: 10.1162/089976602317318938. URL http://www.ncbi.nlm.nih.gov/pubmed/11936959.

Daniel M Wolpert and J Randall Flanagan. Computations underlying sensorimotor learning. Current opinion in neurobiology, 37:7–11, 2016.

Thilo Womelsdorf, Taufik A Valiante, Ned T Sahin, Kai J Miller, and Paul Tiesinga. Dynamic circuit motifs underlying rhythmic gain control, gating and integration. Nature neuroscience, 17(8): 1031–1039, 2014.

Reto Wyss, Peter König, and Paul F M J Verschure. A model of the ventral visual system based on temporal stability and local memory. PLoS biology, 4(5):e120, may 2006. ISSN 1545–7885. doi: 10.1371/journal.pbio.0040120. URL http://journals.plos.org/plosbiology/article?id=10.1371/journal.pbio.0040120.

X Xie and HS Seung. Spike-based learning rules and stabilization of persistent neural activity. Advances in neural information processing …, 2000.

Xiaohui Xie and H Sebastian Seung. Equivalence of backpropagation and contrastive Hebbian learning in a layered network. Neural computation, 15(2): 441–54, feb 2003. ISSN 0899–7667. doi: 10.1162/089976603762552988. URL http://www.ncbi.nlm.nih.gov/pubmed/12590814.

Caiming Xiong, Stephen Merity, and Richard Socher. Dynamic memory networks for visual and textual question answering. arXiv preprint arXiv:1603.01417, 2016.

Min Xu, Si-yu Zhang, Yang Dan, and Mu-ming Poo. Representation of interval timing by temporally scalable firing patterns in rat prefrontal cortex. Proceedings of the National Academy of Sciences, 111(1): 480–485, 2014.

Daniel LK Yamins and James J DiCarlo. Eight open questions in the computational modeling of higher sensory cortex. Current opinion in neurobiology, 37:114–120, 2016a.

DLK Yamins and JJ DiCarlo. Using goal-driven deep learning models to understand sensory cortex. Nature neuroscience, 2016b. URL http://www.nature.com/neuro/journal/v19/n3/abs/nn.4244.html.

Jason Yosinski, Jeff Clune, Yoshua Bengio, and Hod Lipson. How transferable are features in deep neural networks? In Advances in Neural Information Processing Systems, pages 3320–3328, 2014.

Eric A Yttri and Joshua T Dudman. Opponent and bidirectional control of movement velocity in the basal ganglia. Nature, 533(7603): 402–406, 2016.

Chen Yu and Linda B Smith. Joint attention without gaze following: Human infants and their parents coordinate visual attention to objects through eye-hand coordination. PloS one, 8(11):e79659, 2013.

Rafael Yuste, Jason N MacLean, Jeffrey Smith, and Anders Lansner. The cortex as a central pattern generator. Nature reviews. Neuroscience, 6 (6):477–83, jun 2005. ISSN 1471–003X. doi: 10. 1038/nrn1686. URL http://www.ncbi.nlm.nih.gov/pubmed/15928717.

Wojciech Zaremba and Ilya Sutskever. Reinforcement learning neural turing machines. arXiv preprint arXiv:1505.00521, 2015.

A. Zeisel, A. B. M. Manchado, S. Codeluppi, P. Lonnerberg, G. La Manno, A. Jureus, S. Marques, H. Munguba, L. He, C. Betsholtz, C. Rolny, G. Castelo-Branco, J. Hjerling-Leffler, and S. Linnarsson. Cell types in the mouse cortex and hippocampus revealed by single-cell RNAseq. Science, 347(6226): 1138–42, feb 2015. ISSN 0036–8075. doi: 10.1126/science.aaa1934. URL http://science.sciencemag.org/content/347/6226/1138.abstract.

Richard S Zemel and Peter Dayan. Combining probabilistic population codes. In IJCAI, pages 1114–1119, 1997.

Eric A Zilli and Michael E Hasselmo. Coupled noisy spiking neurons as velocity-controlled oscillators in a model of grid cell spatial firing. The Journal of neuroscience, 30(41): 13850–13860, 2010.

D Zipser and RA Andersen. A back-propagation programmed network that simulates response properties of a subset of posterior parietal neurons. Nature, 1988. URL https://cortex.vis.caltech.edu/Papers/PDFsofjournalarticles/Nature/V331{_}88.pdf.

